# Machine-learning dissection of Human Accelerated Regions in primate neurodevelopment

**DOI:** 10.1101/256313

**Authors:** Sean Whalen, Fumitaka Inoue, Hane Ryu, Tyler Fairr, Eirene Markenscoff-Papadimitriou, Kathleen Keough, Martin Kircher, Beth Martin, Beatriz Alvarado, Orry Elor, Dianne Laboy Cintron, Alex Williams, Md. Abul Hassan Samee, Sean Thomas, Robert Krencik, Erik M. Ullian, Arnold Kriegstein, John L. Rubenstein, Jay Shendure, Alex A. Pollen, Nadav Ahituv, Katherine S. Pollard

## Abstract

Using machine learning (ML), we interrogated the function of all human-chimpanzee variants in 2,645 Human Accelerated Regions (HARs), some of the fastest evolving regions of the human genome. We predicted that 43% of HARs have variants with large opposing effects on chromatin state and 14% on neurodevelopmental enhancer activity. This pattern, consistent with compensatory evolution, was confirmed using massively parallel reporter assays in human and chimpanzee neural progenitor cells. The species-specific enhancer activity of assayed HARs was accurately predicted from the presence and absence of transcription factor footprints in each species. Despite these striking *cis* effects, activity of a given HAR sequence was nearly identical in human and chimpanzee cells. These findings suggest that HARs did not evolve to compensate for changes in the *trans* environment but instead altered their ability to bind factors present in both species. Thus, ML prioritized variants with functional effects on human neurodevelopment and revealed an unexpected reason why HARs may have evolved so rapidly.

## Introduction

Human accelerated regions (HARs) are highly conserved sequences that acquired many nucleotide substitutions in humans since we diverged from our common ancestor with chimpanzees and, more recently, from archaic hominins (Franchini and Pollard, 2017; Hubisz and Pollard, 2014). The human-specific accelerated substitution rates in HARs suggest that they are important and that their functions changed during human evolution, perhaps altering traits that distinguish us from chimpanzees and other animals such as morphological differences, our unique diet, reproductive challenges, and cognitive skills (Franchini and Pollard, 2017). Furthermore, it has been hypothesized that HARs and other uniquely human genomic regions could be responsible for our high rates of psychiatric disorders, such as schizophrenia and autism spectrum disorder (ASD), which might be maladaptive by-products of the same changes in the human brain that enabled our unique linguistic and cognitive skills (Burns, 2004; Crow, 1997). Indeed, HARs are enriched in disease-associated loci and nearby genes expressed during embryonic development, especially neurodevelopment (Babbitt et al., 2011; Capra et al., 2013; Doan et al., 2016; Kamm et al., 2013; Pollard et al., 2006a; Prabhakar et al., 2006). Therefore, HARs are exciting candidates for understanding human-specific traits, including our unique susceptibilities to disease.

The majority of HARs (96%) reside in noncoding regions. We previously used machine learning (ML) to predict that at least 30% are developmental enhancers based on their epigenetic and sequence features (Capra et al., 2013). Indeed, several human HAR sequences have been shown to alter evolutionarily conserved enhancer activity, driving changes to transcription factor (TF) expression and uniquely human phenotypes in the limb (HAR2/HACNS1: *GBX2* target gene) (Dutrow et al., 2022), testes (2xHAR.238: *GLI2*) (Norman et al., 2021), skin (2xHAR.20: *EN1*) (Aldea et al., 2021), and brain (HARE5/ANC516: *FZD8*) (Boyd et al., 2015). In addition, fifty-two prioritized HARs have been analyzed for their regulatory activity via enhancer assays in transgenic mice (Boyd et al., 2015; Capra et al., 2013; Franchini and Pollard, 2017; Kamm et al., 2013; Prabhakar et al., 2008), with 31 (60%) functioning as enhancers at the tested points in embryonic development. Nineteen (37%) are active in neurodevelopment, with 14 (27%) driving gene expression in the telencephalon. Of 29 HARs where the human and chimpanzee sequence were both tested, nine (31%) show differences in their expression patterns. These findings indicate that sequence changes in HARs during human evolution can alter developmental gene regulation and phenotypes.

However, the forces that drove the many human-specific sequence variants in HARs after millions of years of conservation remain largely unknown. Most HARs appear to have undergone positive selection on the human lineage prior to our divergence from archaic hominins, while ∼20% have substitution patterns that are consistent with GC-biased gene conversion and a few others show population genetic signatures of ongoing adaptation (Kostka et al., 2012; Pollard et al., 2006a). But it is unknown how much of the accelerated substitution rate in HARs can be attributed to genetic hitchhiking, recurrent positive selection, compensatory evolution to maintain ancestral functions, or other evolutionary forces. Suppose it was adaptive for a HAR to evolve 50% lower enhancer activity in neural progenitor cells. If the HAR was subject to hitchhiking, one of its human-specific variants would have decreased enhancer activity by ∼50%, while the others (the hitchhikers) had little or no effect on enhancer activity. In contrast, if the HAR experienced recurrent positive selection, each of the variants would have incrementally decreased enhancer activity summing to a 50% reduction. In compensatory evolution, the HAR would contain some variants that increase and others that decrease enhancer activity. Thus, learning the contributions of individual variants within a HAR to its enhancer activity could reveal how the HAR evolved to have so many human-specific variants.

Measuring the function of human, chimpanzee, and resurrected ancestral sequences is a powerful approach to these questions. We hypothesized that recently developed ML methods that model the gene regulatory activity of non-coding sequences (Avsec et al., 2021; Vaishnav et al., 2022; Zhou et al., 2018) are capable of predicting how sequence changes alter HAR function. ML has the advantage of being able to leverage massive amounts of epigenetic data and learn complex sequence grammars, while being relatively scalable and cost-effective compared to experimental strategies. Massively parallel reporter assays (MPRAs) are a complementary approach to dissecting variant effects. They measure enhancer function *en masse* with a quantitative readout based on RNA sequencing (Inoue and Ahituv, 2015) and can be applied to real or synthetic sequences, including detection of interactions between variants (Uebbing et al., 2021). In recent years, episomal and lentivirus based MPRAs have been applied to human polymorphisms (Doan et al., 2016), human-chimpanzee fixed differences (Girskis et al., 2021; Uebbing et al., 2021), and modern human-specific substitutions in HARs (Weiss et al., 2021). They have also been used to study human-chimpanzee variants in human-gained enhancers (Uebbing et al., 2021) and introgressed Neanderthal variants (Jagoda et al., 2022). However, none of these MPRAs tested HAR enhancers in non-human primate cells to evaluate how the *trans* environment (Pollen et al., 2019) interacts with *cis* regulatory changes. We therefore saw an opportunity to combine ML and MPRAs in chimpanzee and human neural progenitor cells (NPCs) to decipher the evolutionary forces that drove rapid substitutions in HARs after millions of years of strong negative selection.

In this study, we deeply interrogated the enhancer function of 2645 non-coding HARs from prior studies (Hubisz and Pollard, 2014). By generating 19 new epigenomics datasets and combining them with ML and MPRA, we discovered that human-chimpanzee differences in HAR enhancer activity are primarily determined by nucleotide changes rather than differences in the cellular environment and can be predicted from the presence/absence of transcription factor footprints in each species. We also found striking computational and experimental evidence for compensatory evolution, with multiple variants in the same HAR having opposite effects on enhancer activity, in some cases negatively interacting to maintain ancestral enhancer activity. This new functional understanding is important because almost all HARs showing enhancer activity in NPCs are genetically and physically linked to neurodevelopmental genes and/or neuropsychiatric disease.

## Results

### Chimpanzee and human neural progenitor cells model gene regulation in early forebrain development

In order to establish an *in vitro* system to produce data for modeling the *cis* effects of variants on HAR enhancer function, we generated NPCs from induced pluripotent stem cell (iPSC) lines of two human and two chimpanzee individuals. Though general species-specific regulatory regions have been identified in organoids (Kanton et al., 2019), chimpanzee NPCs have not been used in prior HAR-focused research and are essential for quantifying *trans* effects of the cellular environment. Neural induction was initiated with noggin, a BMP inhibitor, and cells were cultured in retinoic acid-free media supplemented with growth factors FGF and EGF in order to generate early (N2; 12-18 passages) and late (N3: 20-28 passages) telencephalon-fated neural progenitors (**Figure 1**). All lines exhibited normal cell morphology (**Figure 1A,G**) and normal karyotypes (**Figure 1D,J**), as well as neural rosette morphology at an early induction stage and neural progenitor cell morphology at later stages of differentiation (**Figure 1B-C,H-I**). Characterization through immunohistochemistry assays showed that both human and chimpanzee NPCs express neural and glial progenitor proteins such as PAX6 and GFAP (**Figure 1E-F,K-L**). We assessed cell heterogeneity through single-cell RNA-sequencing (scRNA-seq; **Supplemental Table S1**) and observed comparable patterns of telencephalon and radial glia marker expression in human and chimpanzee NPCs (**Figure 1M**). Next, we performed chromatin immunoprecipitation sequencing (ChIP-seq) for the active enhancer-associated histone H3K27ac in N2 and N3 cells from both species, observing high genome-wide concordance between human and chimpanzee NPCs (R^2^ = 0.862 in N2, 0.712 in N3). Just over half of our peaks overlap previously published H3K27ac peaks from developing human (54.9%) and adult chimpanzee (54.3%) cortical tissues (Castelijns et al., 2020; Markenscoff-Papadimitriou et al., 2020). This indicates the relevance of our NPCs to *in vivo* biology while also suggesting that we could discover substantial numbers of novel enhancers in this model of early neurodevelopment.

**Figure 1.**
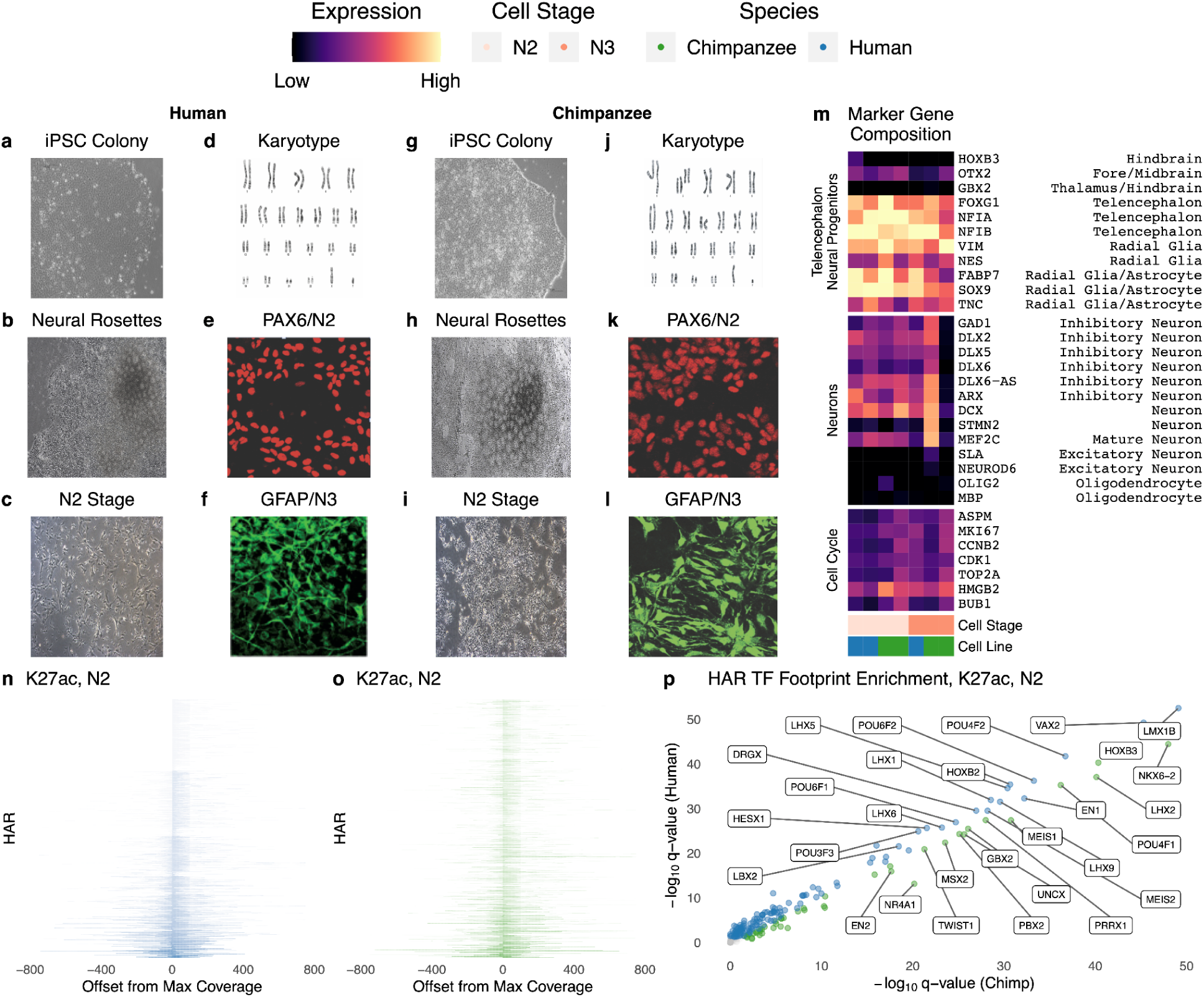
Characterization of chimpanzee and human neural progenitor cells. (**A-C**) Brightfield images of human iPSCs (**A**). iPSC differentiated into neural rosettes (**B**) and N2 cells (**C**) demonstrating typical morphology. (**D**) Human iPSCs demonstrate normal karyotypes. (**E**) Human N2 cells express Paired Box 6 (PAX6), a neural marker. (**F**) Human N3 cells express Glial Fibrillary Acidic Protein (GFAP), a glial marker. (**G-I**) Brightfield images of chimpanzee iPSCs (G). iPSC differentiated into neural rosettes (**H**) and N2 cells (**I**) demonstrating typical morphology. (**J**) Chimpanzee iPSCs demonstrate normal karyotypes. (**K**) Chimpanzee N2 cells express PAX6. (**L**) Chimpanzee N3 cells express GFAP. (**M**) Percentage of cells in scRNA-seq expressing genes that are markers for the cell cycle or telencephalon and neuronal cell types. Human and chimpanzee N2 and N3 cells show comparable marker expression for radial glia and telencephalon. For example, 50-90% of cells expressed FOXG1, a marker of the telencephalon. (**N-O**) Coverage (CPM) of H3K27ac ChIP-seq reads at HARs, sorted by maximum CPM, in human (**N**) and chimpanzee (**O**) N2 cells. (**P**) Human and chimpanzee N2 H3K27ac TF footprints are largely concordant, but some TF families with LIM, POU and homeodomains show species-biased enrichment. Select TFs expressed in NPCs (Schwartz et al., 2015) with large differences in q-value between species are labeled.

### Machine-learning delineation of HAR enhancers using hundreds of epigenetic features

Having established parallel chimpanzee and human iPSC-derived NPCs as a cell culture system for characterizing primate telencephalon development, we undertook a large-scale epigenetic characterization of HARs. We augmented our H3K27ac ChIP-seq by performing the assay for transposase-accessible chromatin (ATAC-seq) and H3K27me3 ChIP-seq for repressed chromatin in human N2 and N3 cells, an early NPC stage (N1; eleven days after initiating neural induction, passage 1), and astrocyte progenitors (**Table 1**). The majority of HARs overlap H3K27ac marks in human (**Figure 1N**) and chimpanzee (**Figure 1O**) NPCs. HARs with high H3K27ac tend to lie in open chromatin (ATAC-seq peaks), whereas those with low H3K27ac tend to overlap peaks of the repression-associated histone H3K27me3, though some HARs have both marks. However, H3K27ac signal at HARs is not highly correlated between human and chimpanzee NPCs, even though it is similar genome-wide and shows limited differences between human N2 and N3 cells (**Supplemental Figure S1**).

**Table 1.**
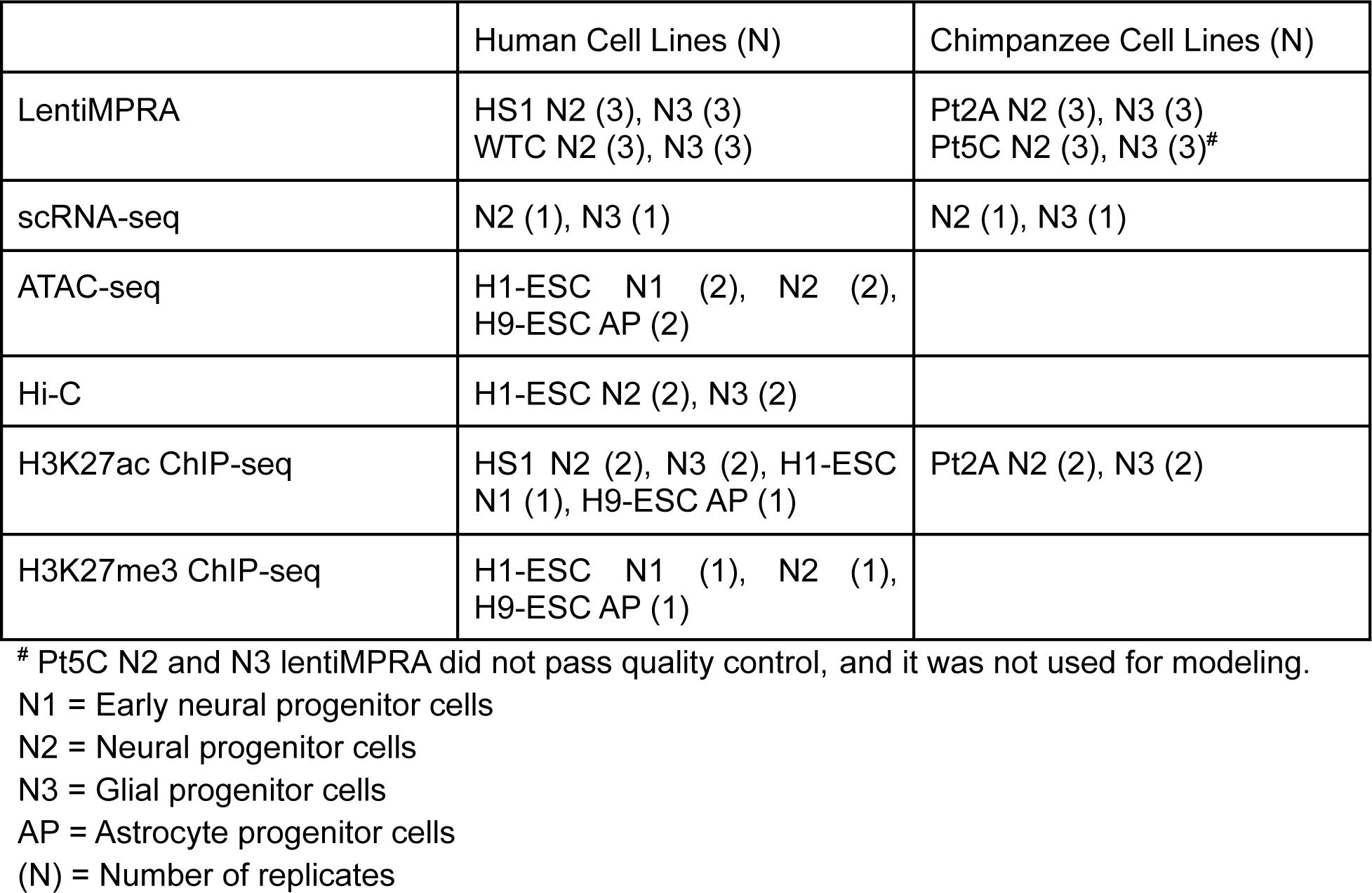
Datasets generated in human and chimpanzee NPCs.

This motivated us to predict transcription factor footprints in HARs with the histone ChIP-seq module of a tool called Hmm-based IdentificatioN of Transcription factor footprints (HINT) (Gusmao et al., 2014), enabling a direct comparison of binding sites in human versus chimpanzee NPCs. We observed many transcription factors whose footprints are differentially enriched between species due to differences in both the sequences and epigenetic profiles of HARs (**Figure 1P**; **Supplemental Table S2**). *EN1, VAX2,* and several other homeobox genes are more enriched in human versus chimpanzee HAR footprints, whereas *NKX6-2* is the most chimpanzee-biased transcription factor. This first epigenetic characterization of early chimpanzee neurodevelopment is consistent with prior observations that many HARs function as enhancers in human cells (Capra et al., 2013; Doan et al., 2016; Girskis et al., 2021; Pollard et al., 2006b; Prabhakar et al., 2008; Uebbing et al., 2021), while revealing differences between human and chimpanzee regulatory potential.

We next sought to relate this *in vitro* epigenetic characterization of HARs to *in vivo* enhancer function using ML. To do so, we leveraged the rich *in vivo* epigenetic data available for many human tissues. Combining our 19 ChIP-seq and ATAC-seq experiments with 254 publicly available epigenetic studies in primary tissue (**Supplemental Table S3**), we found that 70% of HARs (1846/2645) overlap open chromatin and/or active marks in the human brain when considering all developmental stages and brain regions (**Figure 2A**). Significantly fewer HARs (935/2645) have these marks of active regulatory elements in other tissues (Kolmogorov-Smirnov p < 2e-16; **Figure 2B**), despite having similar numbers of datasets. Consistent with multi-tissue enhancer function, 808 HARs have both neural and non-neural marks (**Supplemental Figure S2**). These results emphasize that HARs likely function as enhancers in many contexts beyond neurodevelopment, although brain enhancer-associated epigenetic marks are particularly enriched in HARs, as previously reported in smaller datasets (Bae et al., 2015; Capra et al., 2013; Girskis et al., 2021).

**Figure 2.**
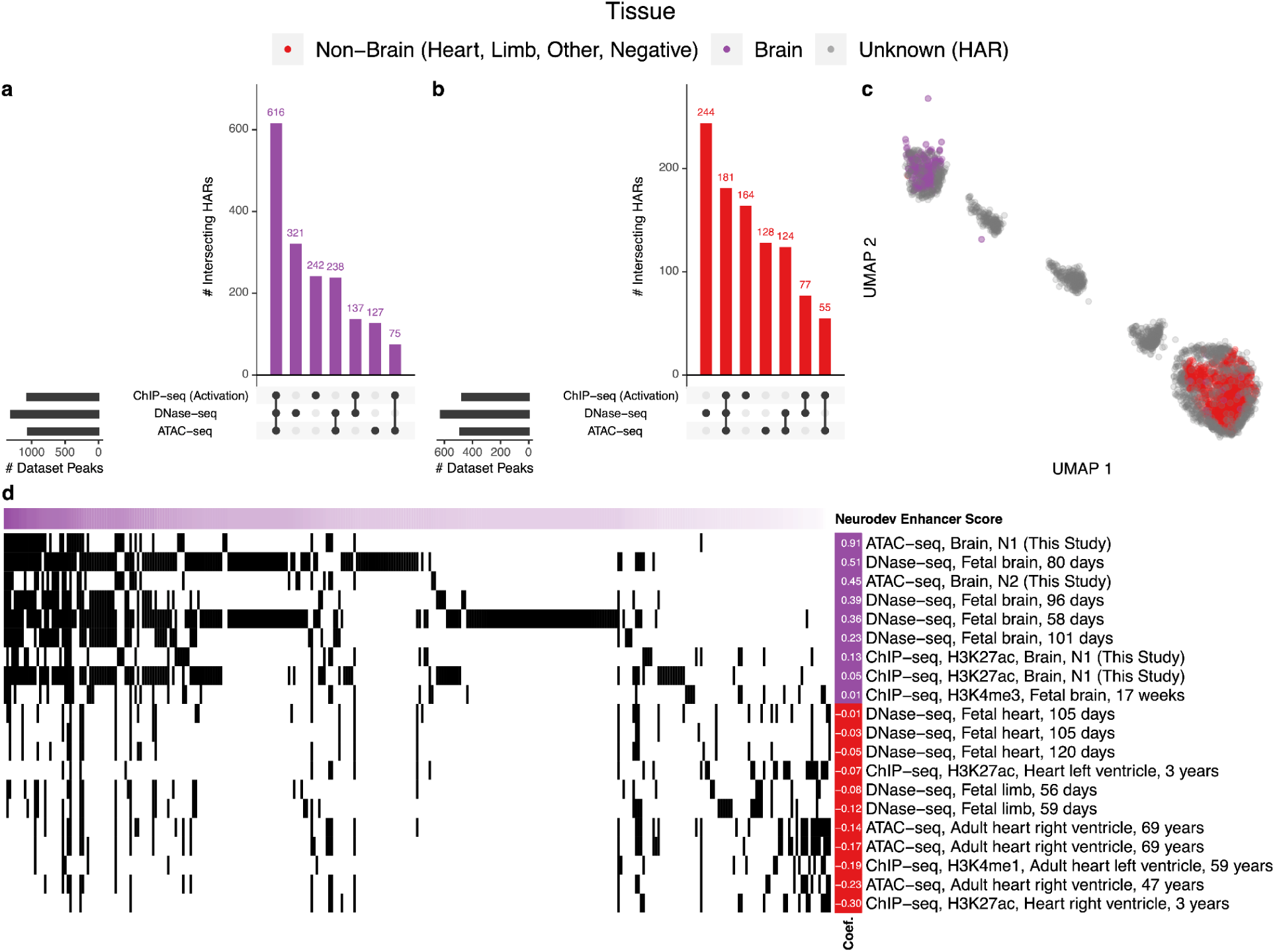
The *in vivo* epigenetic landscape of HARs. A large collection of open chromatin (ATAC-seq, DNase-seq) and ChIP-seq (TF, histone) datasets from human primary tissues (49% brain, 48% heart, 2% limb; GEO and ENCODE accessions in **Supplemental Table S3**) were intersected with HARs. (**A**) Upset plot showing that 616/2645 HARs overlap at least one activating epigenetic mark (H3K4me1, H3K4me3, H3K9ac, H3K27ac, or H3K36me3), one DNase-seq open chromatin region, and one ATAC-seq open chromatin region in brain tissue. A total of 1846/2645 HARs overlap at least one type of open or active chromatin mark. The purple histogram shows the number of HARs with the denoted combination of marks, while the black bars to the left show the number of peaks that overlap a HAR. (**B**) HAR overlaps with activating marks and open chromatin in other tissues. There are significantly more overlaps for the brain compared to non-brain tissues (p-value < 2e-16). Joint heart and brain overlaps are shown in **Supplemental Figure S2**. (**C**) Two-dimensional UMAP projection of HARs (grey) with VISTA heart (red) and brain (purple) enhancers (Visel et al., 2007) showing that some HARs cluster with *in vivo* validated enhancers. (**D**) HARs (horizontal axis, sorted so those most similar to VISTA brain enhancers are on the left) with their epigenetic profiles (vertical axis; black indicates overlapping epigenetic features). Shown are the epigenetic features most predictive in a ML model of VISTA brain enhancers (purple) versus non-brain enhancers (VISTA negatives plus enhancers active in other tissues; red), along with their model coefficients (left).

To connect this epigenetic profiling of HARs to *in vivo* enhancer activity, we visualized the co-embedding of HARs and validated developmental enhancers from the VISTA Enhancer Browser (Visel et al., 2007). We chose VISTA because it measures tissue-specific enhancer activity during embryonic development, and most of the tested sequences are evolutionarily conserved, similar to HARs. First, we annotated each human genomic region in VISTA and each HAR with binary vectors denoting genomic overlap or not with peaks from each dataset in our epigenetic compendium. Embedding the high-dimensional vectors for VISTA enhancers in two dimensions, we observed that neurodevelopmental enhancers have distinct signatures compared to sequences active only in non-brain tissues or without enhancer activity. Then, we co-embedded HARs in this same epigenetic space and discovered that many HARs cluster with *in vivo* validated neurodevelopmental enhancers, while others appear not to function as enhancers or to be active in other tissues and developmental stages (**Figure 2C**).

This clear partitioning of HARs motivated us to score HARs based on having embryonic brain enhancer-like epigenetic profiles or not. To do so, we first trained a supervised ML model using epigenetic signatures to distinguish VISTA brain enhancers from enhancers that are inactive or active in other tissues. This is a more difficult classification problem than simply predicting enhancers versus non-enhancers, especially given the large number of multi-tissue enhancers containing overlapping epigenetic signatures. Nonetheless, we established that a L1-penalized logistic regression model can distinguish neurodevelopmental enhancers in held-out VISTA data (median cross-chromosome auPR 0.69, auROC 0.8). Using this model, we then scored HARs based on how consistent their epigenetic profiles are with neurodevelopmental enhancer function (**Figure 2D**). As expected, HARs with higher scores overlap more neurodevelopmental epigenetic marks and have similar co-embedding coordinates to VISTA brain enhancers (**Supplemental Table S7**). Thus, ML models are able to integrate hundreds of epigenetic signals to prioritize HARs likely to function as enhancers during neurodevelopment.

### 2xHAR.183 is a *ROCK2* neuronal enhancer

Next, we sought to validate a novel HAR enhancer prediction. We generated chromatin capture (Hi-C) data in our human N2 and N3 cells and used it along with Hi-C from primary fetal brain tissue (Song et al., 2019, 2020; Won et al., 2016) to associate HARs with genes they may regulate. This analysis confirmed known regulatory relationships between HARs and developmental genes, including 2xHAR.20 with *EN1* (Aldea et al., 2021) and 2xHAR.238 with *GLI2* (Norman et al., 2021). Based on chromatin contacts with the neurodevelopmental gene *ROCK2* in NPCs (**Figure 3A**) as well as a PLAC-seq loop to *ROCK2* in excitatory neurons (Song et al., 2020), we selected 2xHAR.183 for functional characterization. This HAR has a high neurodevelopmental enhancer score in our epigenetics-based ML model and overlaps enhancer annotations from ChromHMM (Ernst and Kellis, 2017) and FANTOM5 (Lizio et al., 2015). Consistent with *ROCK2*’s increasing expression in later stages of embryonic development and at postnatal time points in mice (Lein et al., 2007) (mid-gestation in humans), we observed progressively more open chromatin and greater H3K27ac signal at 2xHAR.183 over the developmental stages in our epigenetic compendium, with a slightly larger activation signature in chimpanzee compared to human cells/tissue (**Figure 3A**). Supporting the hypothesis that 2xHAR.183 is a neurodevelopmental enhancer, footprint analysis in our human NPCs and ENCODE fetal brain tissue (Funk et al., 2020) identified binding sites for C/EBPBeta, PRDM1, BCL11A, and RFX2 (**Figure 3B**). To test if 2xHAR.183 indeed regulates *ROCK2* or other genes in the locus, we performed CRISPR activation (CRISPRa) in human NGN2 induced iPSC (WTC11) derived neurons (Wang et al., 2017) and observed increased expression of *ROCK2* but not *E2F6,* a nearby gene in the adjacent chromatin domain (**Figure 3C**). These findings indicate that 2xHAR.183 is a *ROCK2* enhancer in developing neurons.

**Figure 3.**
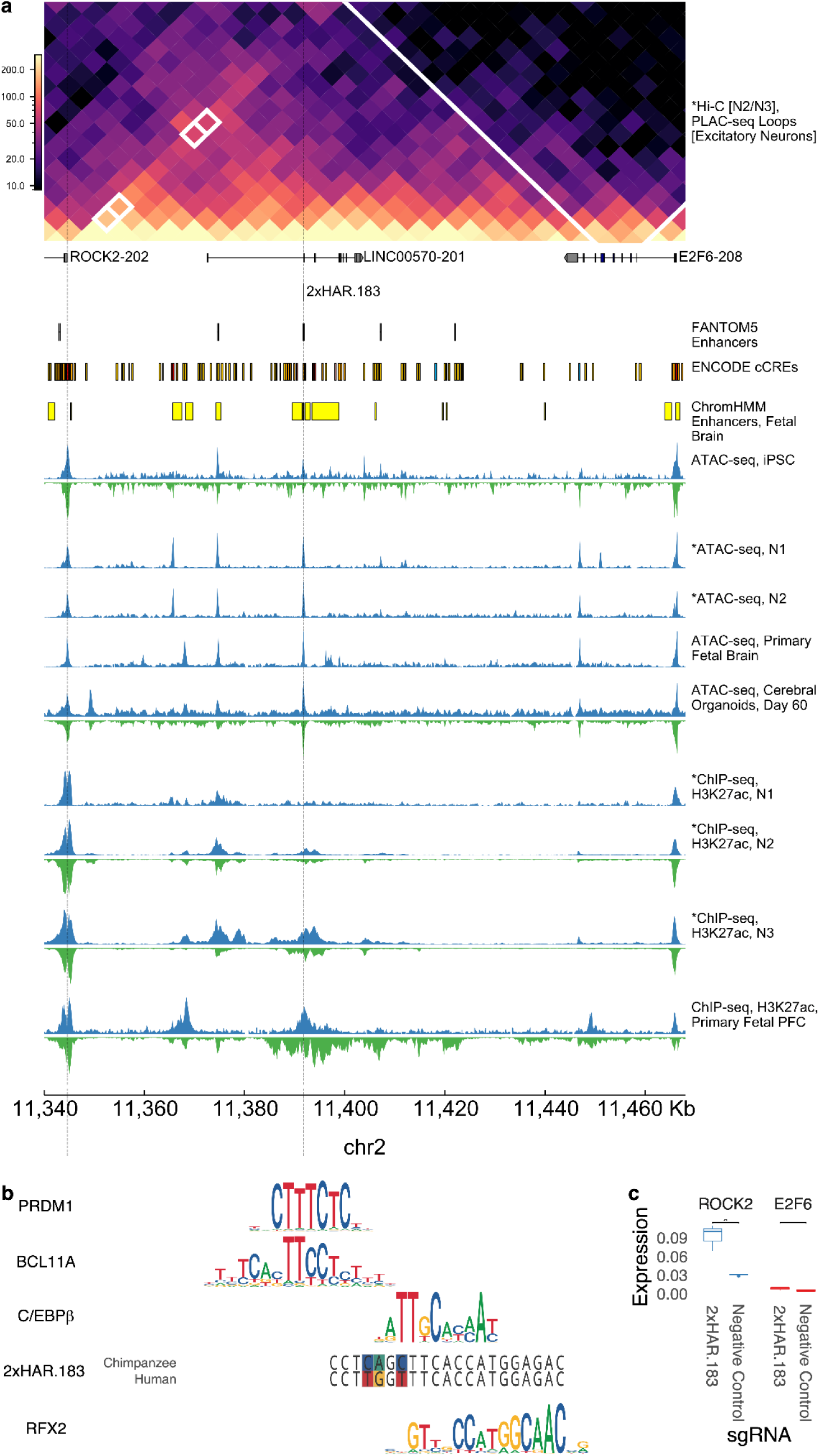
Validation of an active HAR enhancer regulating ROCK2. 2xHAR.183 was selected for further validation due to its high predicted enhancer score in our epigenetic modeling (**Figure 2**). (**A**) 2xHAR.183 has a significant chromatin loop with the *ROCK2* gene in excitatory neuron PLAC-seq data (5kb resolution binary loop call) (Song et al., 2020) and contacts *ROCK2* in our N2/N3 Hi-C. The gene *E2F6* is nearby on the linear genome but has fewer 3D chromatin contacts. 2xHAR.183 overlaps multiple annotations from fetal brain datasets. Chimpanzee and human epigenetic datasets across early neurodevelopment suggest 2xHAR.183 starts and remains accessible in both species, while gaining acetylation beginning at the neural progenitor (N2) stage. The activation signature appears later and stronger in chimpanzee versus human cells. (**B**) The human and chimpanzee alleles of 2xHAR.183 overlap footprints of known neurodevelopmental TFs, some of which overlap human:chimpanzee variants. (**C**) CRISPRa validation (3 replicates per target, 4 per control) shows 2xHAR.183 drives strong expression of *ROCK2*, but not the proximal gene *E2F6*. Variability between replicates is small for low expression values.

### Deep learning predicts that most individual HAR variants alter enhancer activity

We sought to dissect how each nucleotide change that occurred in a HAR during human evolution altered enhancer activity. We first approached this question by utilizing the deep-learning model Sei (Chen et al., 2022) that predicts how human polymorphisms alter tissue-specific regulatory activity. By instead presenting human-chimpanzee fixed differences within HARs to Sei, we were able to predict whether each variant alters chromatin states in various tissues, including brain enhancer activity (**Supplemental Table S5**). This revealed that most HAR variants shift enhancer activity in at least one tissue (**Figure 4A**). Chromatin state changes for HAR variants are generally correlated across different tissues (**Supplemental Figure S3**), indicating that the variant increases or decreases enhancer activity consistently. However, we found several variants with tissue-specific effects such as trade-offs between brain and B-cell activity enhancer activity in HAR3 and HAR166, as well as a 2xHAR.170 variant predicted to decrease enhancer activity in brain tissue while increasing activity in all other tissues (**Figure 4E**). These results demonstrate that deep learning can be used to generate testable hypotheses about HAR variant function.

**Figure 4.**
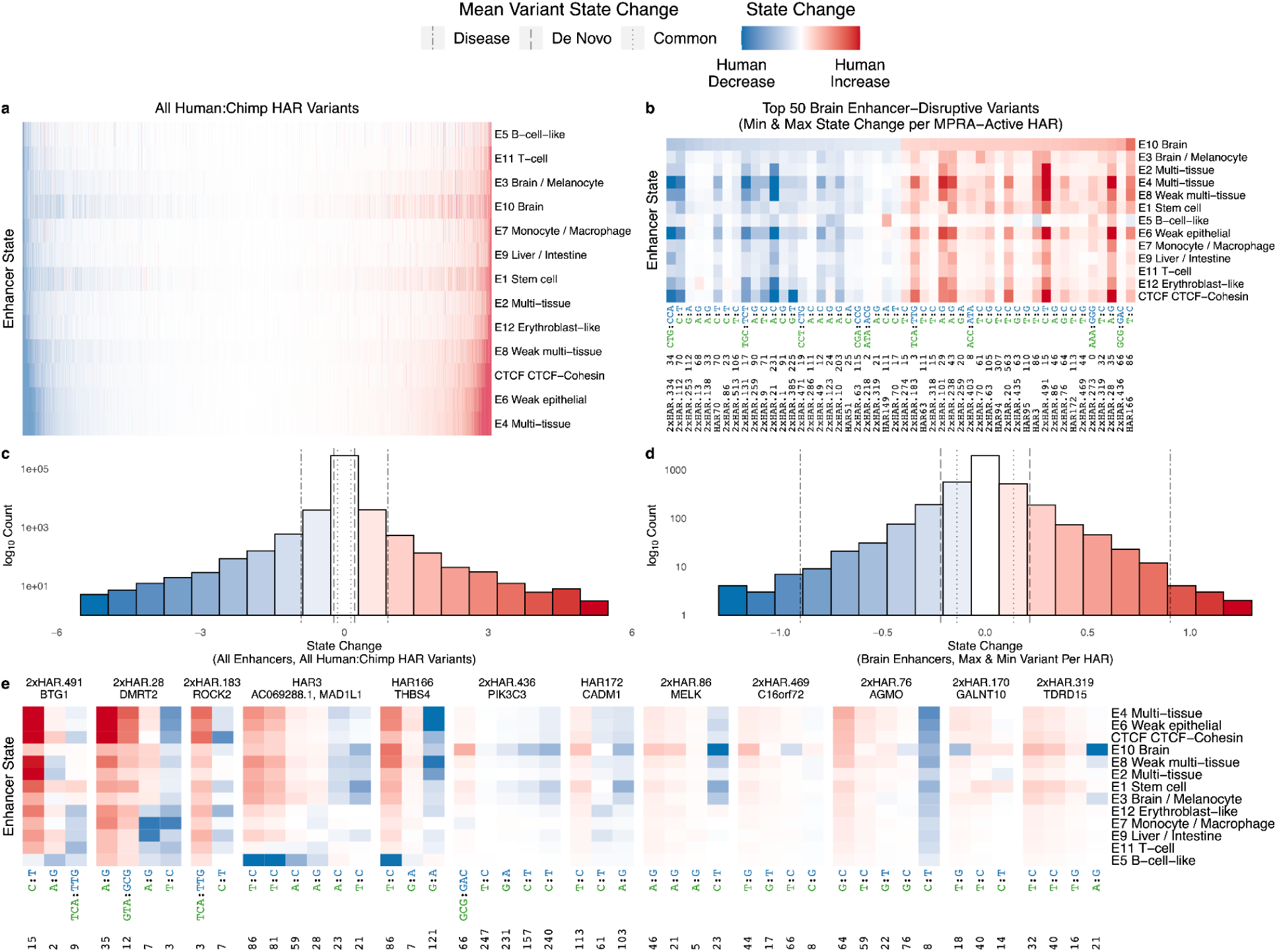
Human-specific variants shift HAR enhancer profiles in a deep-learning model. Every human-specific variant in each HAR was evaluated using the deep-learning model Sei (Chen et al., 2022). Variants where the human nucleotide decreases the chromatin state are blue (shade denotes amount of decrease), variants where the human nucleotide increases the chromatin state are red, and complex variants that can not be scored by Sei are white. (**A**) The landscape of chromatin state changes (y-axis) induced by all human:chimpanzee variants across all HARs (x-axis), sorted by predicted impact on brain enhancer state. (**B**) The 50 HAR variants that most increase or decrease brain enhancer state for all HARs that were active in our MPRA. The x-axis shows the HAR name, the offset of the variant from the HAR’s start position, and the human and chimpanzee alleles colored by species and separated by a colon. (**C**) Histogram of predicted enhancer state changes for all HARs variants from (**A**). Mean state changes for different classes of variants (Chen et al., 2022) are shown via vertical lines: 1000 Genomes common variants, de novo mutations in healthy individuals, disease-causing mutations (from smallest to largest mean change). Many HAR variants have effects that exceed those of phenotype-associated human polymorphisms. (**D**) Histogram of predicted brain enhancer state changes for the most disruptive HAR variants in active HARs. Mean state changes for different classes of variants (Chen et al., 2022) as in (**C**). (**E**) For 12 HARs containing variants with the largest effects on brain enhancer activity in our Sei analysis, we observed a mix of variants predicted by Sei to increase and decrease enhancer activity. Variants (x-axis) are annotated with their offset from the start of the HAR plus the human and chimpanzee alleles separated by a colon. HARs are annotated with the closest protein-coding gene.

To contextualize these results, we evaluated the magnitude of the HAR enhancer state changes predicted by Sei. Suggestive of functionally important changes, the mean of the largest tissue-specific shift per HAR variant (0.54) exceeds that of both common variants from the 1000 Genomes Project (0.139) and *de novo* mutations in healthy individuals (0.217), but is less than disease mutations in the Human Gene Mutation Database (0.903) (Chen et al., 2022). Using these averages as thresholds, we identified 2121 HAR variants (16%) with predicted absolute effects on brain enhancer activity (Sei state E10) greater than expected compared to common variants, 1226 (9%) compared to *de novo* variants, and 61 (< 1%) greater than disease mutations (**Figure 4B** **& D**; **Supplemental Table S5**). Variants predicted to increase activity are more common than those predicted to decrease activity, though effect sizes are slightly larger for decreases overall (**Figure 4C**) and when considering only the most brain enhancer disruptive variant per HAR (**Figure 4D**). Thus, deep learning enabled us to computationally screen all individual HAR variants for effects on enhancer activity across tissues and predict that a substantial number of individual HAR variants changed enhancer activity during human evolution.

### Many HARs contain variants predicted to have opposing effects on enhancer activity

Moving beyond testing individual HAR variants, we examined Sei predictions for all variants within the same HAR. To our surprise, 43% of HARs contain a mix of variants predicted to increase and decrease enhancer activity beyond the average effect of common variants genome-wide (**Figure 4E**), and 14% of HARs contain variants with opposing effects on neurodevelopmental enhancer activity. This is significantly more than expected by chance (bootstrap p=0.03). Limiting this analysis to variants whose effects exceed the mean of *de novo* or disease-causing variants, we observe two or more strongly opposing variants in 30% and 3% of HARs, respectively. Furthermore, many HARs contain individual variants whose effect on enhancer activity is greater than the net effect of all the variants in that HAR. This signature led us to hypothesize that compensatory evolution to fine tune enhancer activity and possibly maintain ancestral activity levels, rather than recurrent selection to successively increase or decrease activity, drove rapid evolution of some HAR enhancers. It is not currently possible to test for variant interactions in the Sei framework, motivating us to move from *in silico* to *in vitro* characterization of HAR variants.

### MPRA characterization of HAR variants in primate NPCs

Performing a massive ML integration of data from epigenetic assays, transgenic mice, transcription factor motifs, and human genetic variants generated several testable hypotheses about HAR enhancer function. First, we predicted that a substantial number of HARs function as enhancers in the developing brain, consistent with prior work. Second, species differences in H3K27ac and transcription factor footprints suggested that the human and chimpanzee sequences of many HARs are differentially active. Going beyond other studies, we additionally hypothesized that many HARs contain variants with opposing effects on enhancer activity. Finally, we found evidence suggesting that HAR variants interact rather than having additive effects on enhancer function, raising the question of whether there might also be interactions with the chimpanzee versus human cellular environment. To gather additional data regarding each of these aspects of HAR enhancer function, we used MPRAs to compare the activity of homologous human and chimpanzee sequences (*cis* effects) in the *trans* environments of chimpanzee and human NPCs. Interrogating different permutations of HAR variants in cells from both species distinguish this experiment from prior MPRA studies.

To quantitatively dissect the effects of nucleotide variants in HARs, we designed an oligonucleotide (oligo) library containing the human and chimpanzee sequences of 714 HARs from our prior studies (Lindblad-Toh et al., 2011; Pollard et al., 2006a, 2006b), all potential evolutionary intermediates between the human and chimpanzee sequences (“permutations”) of three HARs (2xHAR.164, 2xHAR.170, 2xHAR.238) with prior evidence of differences in neurodevelopmental enhancer activity between human and chimpanzee sequences (Capra et al., 2013), 118 positive controls, and 142 negative controls (**Methods**). We performed lentivirus based MPRA (lentiMPRA) with this library in two different human and chimpanzee N2 and N3 cell lines (**Supplemental Figure S4**). For each condition, we generated three technical replicates, yielding 18 measurements of enhancer activity for each sequence after quality control. We observed high correlation (median R^2^ = 0.91) between replicates (**Figure 5A**).

**Figure 5.**
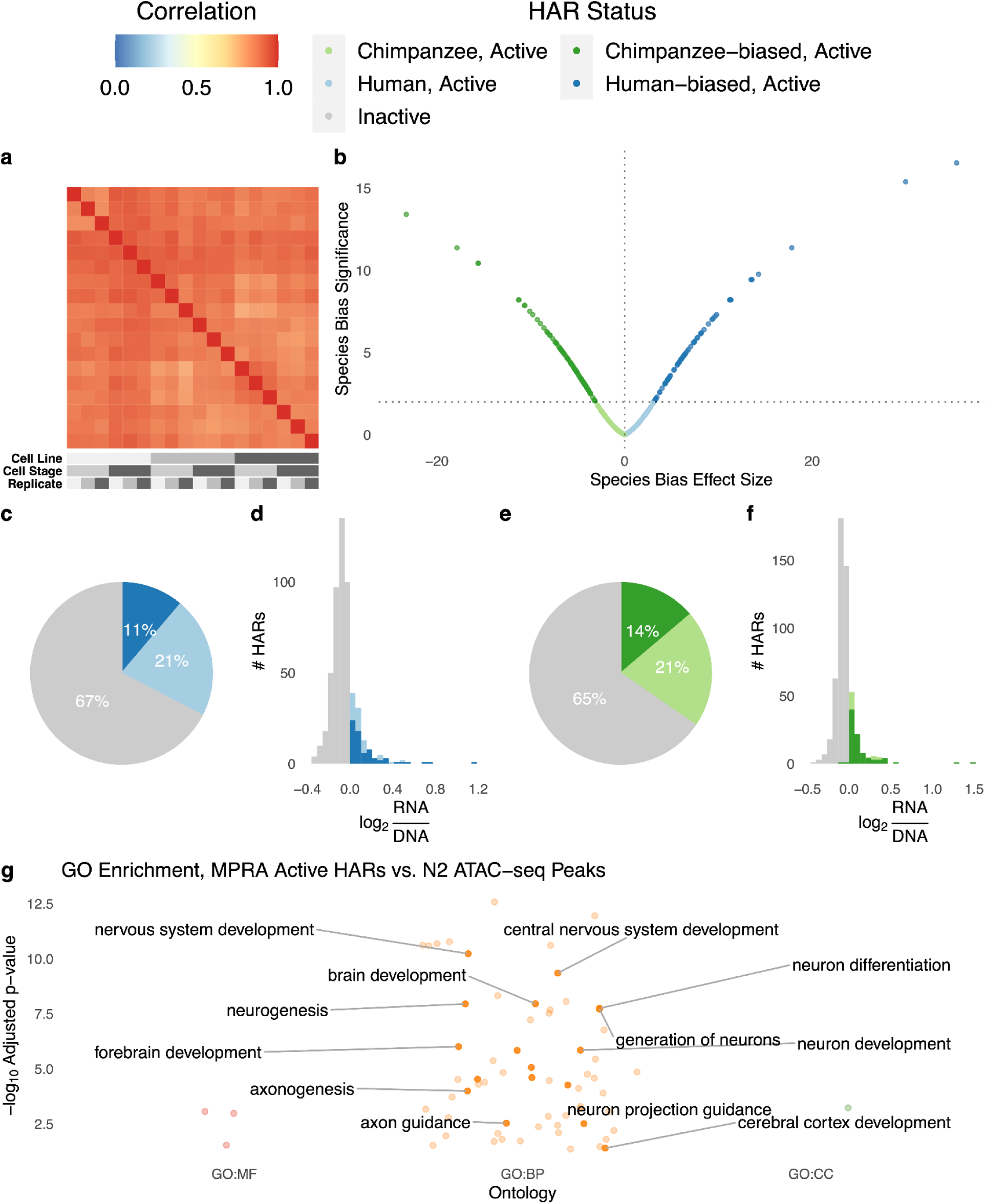
Species-biased HAR enhancers identified in chimpanzee and human NPCs. We performed MPRAs in chimpanzee and human cell lines at the N2 and N3 stages of differentiation. (**A**) Enhancer activity (RNA/DNA ratios batch corrected and normalized for sequencing depth) was highly correlated between technical and biological replicates for eighteen samples passing quality control: 3 replicates (shades of grey) of Pt2a (chimpanzee; dark grey), WTC (human; medium grey), and HS1-11 (human; light grey) iPSC lines differentiated into N2 (medium grey) and N3 (dark grey) cells. (**B**) Effect size (t-statistic) vs significance (-log10 q-value) for the ratio of human and chimpanzee HAR sequence activity for active HAR enhancers. HARs with species-biased activity are plotted in dark green (chimpanzee sequence more active) or dark blue (human sequence more active). (**C**) Roughly a third of human HAR sequences are active across samples (log RNA/DNA > median of positive controls in at least 9/18 replicates), and 11% are human-biased (differentially active with human:chimpanzee ratio > 1). (**D**) Distribution of human HAR sequence enhancer activity for inactive (grey) or active HARs, with active split into human-biased (dark blue) versus not (light blue). (**E**) Roughly a third of chimpanzee HAR sequences are active across samples, and 14% are chimpanzee-biased. (**F**) Histogram of chimpanzee sequence activity as in (**D**). (**G**) HARs active in the MPRA are enriched for many neurodevelopmental GO terms. Colors indicate the type of term: red = molecular function (MF), orange = biological process (BP), green: cellular compartment (CC).

Before comparing human and chimpanzee alleles, we first identified a subset of 293 HARs with activity above the median of positive controls in at least 50% of samples for either the human or chimpanzee sequence. These constitute about one-third of both human and chimpanzee HAR sequences (**Figure 5C-F**) and include 2xHAR.183. The majority of active HARs (233/293) are in a chromatin domain or loop with a neurodevelopmental gene (**Supplemental Table S7**), and these loci are enriched for roles in neurodevelopment, transcription, cell adhesion, axon guidance and neurogenesis (**Figure 5G**, **Supplemental Table S6**).

To validate our lentiMPRA, we compared active HARs to published mouse transgenic reporter assays, mostly performed at embryonic day (E) 11.5, a developmental time point similar to N2 (**Supplemental Table S4**). We found significant concordance with *in vivo* expression for embryonic brain (odds ratio = 3.79, Fisher’s exact test p=0.005) and telencephalon (odds ratio = 7.44, p = 0.00012). We performed mouse reporter experiments for an additional four HARs (HAR152, 2xHAR.133, 2xHAR.518, 2xHAR.548) at developmental stages chosen based on expression of nearby genes and observed enhancer activity for all four (**Supplemental Figure S5**). Next, we performed luciferase assays in one human and one chimpanzee cell line for nine active HARs and observed that six were more active than an empty vector (**Supplemental Figure S6**). Finally, we quantified activity of H3K27ac versus H3K27me3 peaks included as controls in our lentiMPRA, and we observed significantly higher activity for H3K27ac in all samples as expected (**Supplemental Figure S7**). These data indicate that our lentiMPRA identified *bona fide* neuronal enhancers.

Nonetheless, we observed only moderate correlation between neurodevelopmental enhancer scores from our ML model trained on VISTA (**Figure 2D**) and activity levels in NPC lentiMPRA. To investigate this expected difference (Kwasnieski et al., 2014; Lindhorst and Halfon, 2022) (see Discussion), we used the fact that HARs cluster based on their epigenetic profiles (**Figure 2C**) to explore datasets that delineate HARs where ML and lentiMPRA results are concordant versus discordant. We also trained a classifier to distinguish HARs with high ML scores (top 25%) but low lentiMPRA activity (bottom 25%) from HARs with low ML scores but high MPRA activity (auPR 0.96) based on their epigenetic profiles. Analyzing predictive features revealed that lentiMPRA is more permissive, allowing some sequences with closed chromatin in the brain or activating marks outside the brain to show activity in NPCs, whereas the ML model is more tissue-specific (**Supplemental Figure S8**). We also found that HARs prioritized by ML but not lentiMPRA tend to have active marks in whole fetal brain or brain cell types distinct from forebrain neurons (e.g., astrocytes, hippocampal neurons). These results are consistent with lentiMPRA having been performed *in vitro* in NPCs and the ML model having been trained to identify VISTA *in vivo* brain enhancers, which reflect a variety of cell types and regions within the brain. We conclude that it is important to consider the complementary sets of HAR enhancers identified via each approach, as we have done here, with the 48 HARs in the top quartile of both ML and lentiMPRA being particularly high-confidence neurodevelopmental enhancers.

### HAR sequence variants alter enhancer activity while the cellular environment does not

Leveraging the fact that we assayed human and chimpanzee HAR sequences side-by-side across cellular environments, we next assessed evidence that lentiMPRA activity levels differed between chimpanzee and human NPCs. We observed strikingly similar activity of HAR enhancers across not only technical but also biological replicates (**Supplemental Figure S9**), including different cell species and stages (**Figure 5A**). In contrast to these limited *trans* effects, many HARs show consistent differences in activity between human and chimpanzee sequences (**Supplemental Figure S10**). These *cis* effects were statistically significant for 159 HARs (54% of active HARs) at a false discovery rate (FDR) < 1% (**Figure 5B**, **Supplemental Table S7**). HARs where the human sequence has increased activity (70 human-biased HARs; **Figure 5C-D**) are slightly less common than those with decreased activity (89 chimpanzee-biased; **Figure 5E-F**), though effect sizes are generally similar (**Figure 5B**). 2xHAR.548, located in a chromatin domain with the neurodevelopmental regulator and disease gene *FOXP1*, was the most species-biased HAR, showing much higher activity for the human compared to chimpanzee sequence. These results quantitatively demonstrate that *cis* regulatory features are stronger drivers of HAR enhancer activity than the cellular environment. This observation was possible because we performed lentiMPRA in both human and chimpanzee cells.

To validate species-biased HARs, we tested nine homologous chimpanzee and human HAR sequences with luciferase and confirmed statistically significant bias in the expected direction for six (**Supplemental Figure S6**). Furthermore, out of six active HARs that have previously shown differential activity between the human and chimpanzee sequence in mouse transgenics (Aldea et al., 2021; Capra et al., 2013; Norman et al., 2021; Prabhakar et al., 2008) (2xHAR.20, 2xHAR.114, 2xHAR.164, 2xHAR.170, 2xHAR.238), all except 2xHAR.164 were also species-biased in our lentiMPRA (**Supplemental Table S7**). These results are strong evidence that our lentiMPRA accurately detected species-biased HAR enhancer activity.

### Species differences in HAR enhancer activity can be predicted from transcription factor footprints

Most brain-expressed TFs have footprints overlapping multiple HAR variants (**Supplemental Table S2**), and some of these also have large brain enhancer activity decreases (**Figure 6A**) or increases (**Figure 6B**) in our Sei analysis. Furthermore, despite the *trans* environment (e.g., expression level of TFs) being similar, there are species differences in the footprints within orthologous HARs (**Figure 1P**) due to sequence and epigenetic changes in human versus chimpanzee NPCs. This motivated us to model the species-biased enhancer activity we observed in lentiMPRA using the human and chimpanzee footprints of each HAR. A supervised gradient boosting regressor (**Methods**) was able to predict human:chimpanzee lentiMPRA log ratios from human and chimpanzee footprints with very low error (R^2^=0.8, RMSE=0.04), indicating that loss and gain of TF binding sites is a plausible mechanism through which HAR enhancer activity changed during human evolution. This *cis* mechanism is consistent with our observing similar activity for HAR sequences in human versus chimpanzee NPCs.

**Figure 6.**
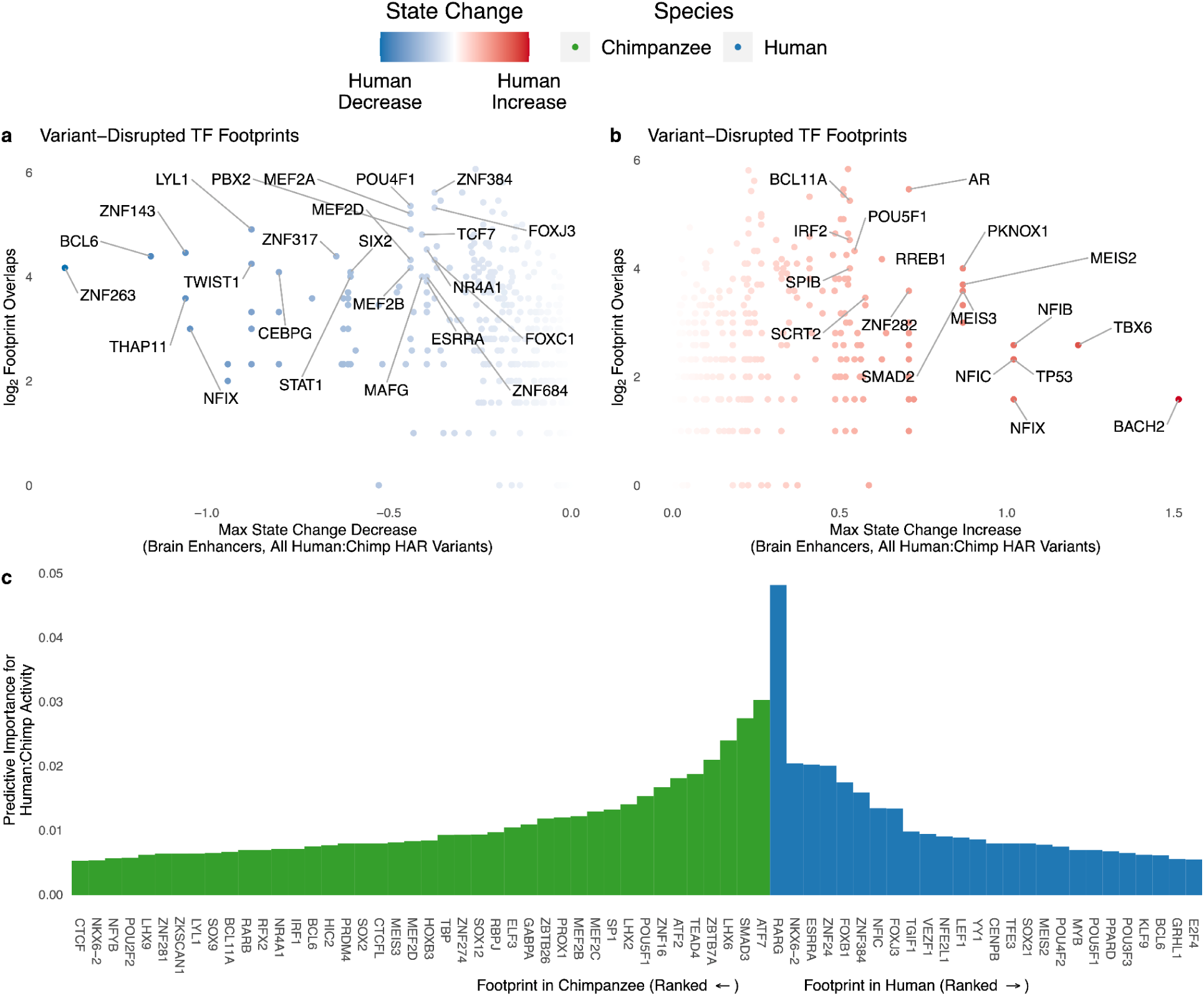
Variants in transcription factor footprints predict HAR species bias. (**A**) The effects of HAR variants in TF footprints (in human N2 H3K27ac ChIP-seq) on brain enhancer activity were predicted using Sei (Chen et al., 2022). For each TF, the largest decrease in brain enhancer state over all variants (x-axis) is shown against the number of variant-containing footprints (y-axis). Select TFs expressed in NPCs (TPM > 1) and scoring high on one or both metrics are labeled. (**B**) TFs with the largest predicted increase in brain enhancer activity in the analysis from (**A**). (**C**) The species-bias of HAR MPRA activity can be predicted accurately using human and chimpanzee N2 H3K27ac footprints for TFs expressed in NPCs as features in a gradient boosting model. The most important TFs for accurate predictions are shown along with their variable importance scores.

Next, we used variable importance to assess which TF footprints contribute most to this predictive accuracy (**Figure 6C**). A TF can be important due to its human footprints, its chimpanzee footprints, or both. In each case, the model may leverage a positive or negative association with human:chimpanzee lentiMPRA activity. This analysis highlighted genes associated with neurological disease (MEF2C, NKX6-2) and brain development (FOXB1, ZNF24), as well as TEAD4, which regulates organ size. Other functions represented amongst the top TFs were regulation of cell proliferation and differentiation (SMAD3, LHX2, LHX6, ZNF16, ZBTB7A, POU5F1, FOXJ3, SP1, MEF2B), retinoic acid and estrogen dependent regulation (RARG, ESRRA), chromatin regulation (ATF2, ATF7), and extracellular matrix regulation (ZNF384). Collectively, these results show that changes in HAR neurodevelopmental enhancer activity during human evolution can be accurately recapitulated by the losses and gains of TF footprints.

### HAR enhancers are linked to neurodevelopmental gene expression and psychiatric disorders

To aid with interpretation of HAR enhancers in the context of neurobiology and disease, we used linkage disequilibrium to associate HAR variants with neuropsychiatric disorder single nucleotide polymorphisms (SNPs) (Sullivan et al., 2018) and brain expression and chromatin quantitative trait loci (QTLs) (GTEx Consortium, 2013; Liang et al., 2021; Wang et al., 2018; Werling et al., 2020) (**Supplemental Table S7**). We found 55 HAR enhancers with genetic associations to psychiatric disease and/or brain gene expression, many of which also have chromatin interactions with neurodevelopmental genes (**Supplemental Figure S11**). We discovered that 2xHAR.170 (i) has a long-range chromatin interaction with *GALNT10*, (ii) harbors a SNP (rs2434531) that is an expression QTL for *GALNT10* (Wu et al., 2020), and (iii) is in linkage disequilibrium with a schizophrenia associated SNP (rs11740474) (Hormozdiari et al., 2017). Along with our lentiMPRA results, these data strongly implicate 2xHAR.170 as a *GALNT10* enhancer with variable activity across people that may be linked to schizophrenia risk. Other notable examples of disease associated HARs include 2xHAR.502, which lies in an intron of the language and schizophrenia associated gene *FOXP2* and contains a SNP associated with attention deficit hyperactivity disorder.In addition, 2xHAR.262 lies in a contact domain with *CPSF2*, *RIN3*, and *SLC24A4* and contains a SNP associated with bipolar disorder. Collectively, we linked the majority of active HARs to neurodevelopmental genes and associated 20 with psychiatric diseases, underscoring the phenotypic consequences of altering the activity of these deeply conserved enhancers.

### Variants within individual HARs interact to tune enhancer activity

To dissect HAR lentiMPRA activity at the single-nucleotide level, we used the permutation oligos. We first considered all oligos that represent a single chimpanzee variant inserted into the human HAR sequence. This parallels our deep learning variant interpretation with Sei, enabling a direct comparison. Across all three HARs, Sei’s brain enhancer state correlated loosely with lentiMPRA activity (**Figure 7A**, **Supplemental Figure S12**). We observed the strongest concordance for 2xHAR.170, a HAR that is species-biased in lentiMPRA and in transgenic mouse embryos (**Figure 7D,E**). Next, we used the permutation lentiMPRA to look for evidence of compensatory evolution (i.e., negative interactions between variants). We fit a model (**Methods**) to predict enhancer activity from the unique combination of human:chimpanzee sequence differences present in each oligo. We observed that all three tested HARs contain opposing variants. To confirm this finding, we generated a second lentiMPRA library containing only permutation oligos, observing highly correlated activity measurements (**Supplemental Figure S13**) and concordant results. Thus, permutation lentiMPRA revealed that human-specific mutations in HARs interact non-additively both to buffer and to amplify each other’s effects on enhancer activity during neurodevelopment.

**Figure 7.**
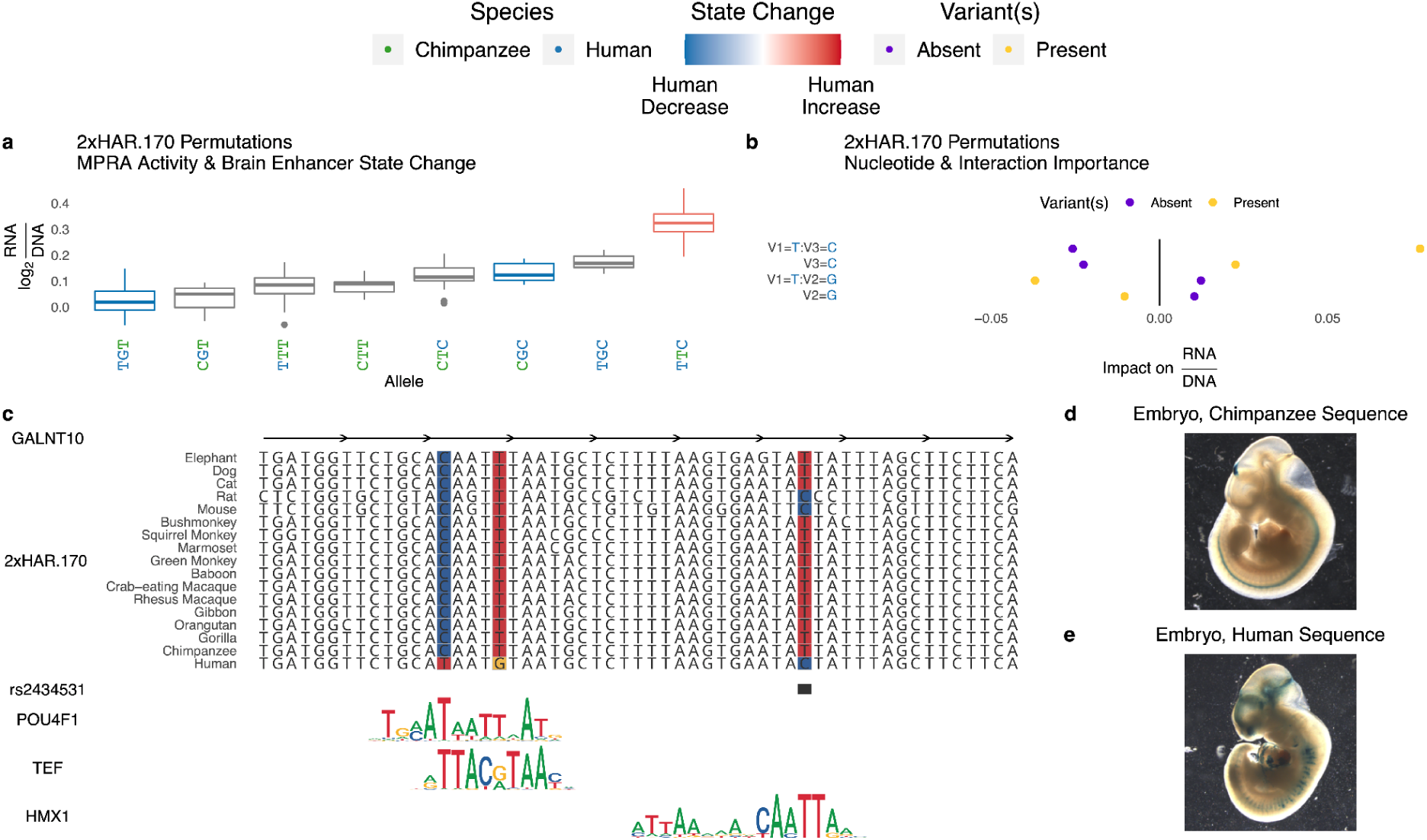
Variants in HARs interact to tune enhancer activity. (**A**) All evolutionary intermediates between chimpanzee and human alleles of 2xHAR.170 were tested via MPRA. Individual variants showed a range of effects on MPRA activity (y-axis; log2(RNA/DNA)) that correlated with Sei predicted effects on brain enhancer activity (red = human increase, blue = human decrease, no color = untestable). MPRA oligos containing permutations of variants revealed interactions between variants. (**B**) We assessed the importance of each variant using a gradient boosting model that predicts MPRA activity of each permutation using the presence or absence of the human allele at each of the three variants. Interactions between multiple variants (separated by colons on the y-axis) were included as predictors alongside main effects (no colon) to assess their predictive importance. This model confirmed the importance of specific variant interactions (x-axis, positive = higher predicted activity, negative = lower activity). Variant names consist of a V followed by the variant number (1, 2 or 3 ordered from 5’ to 3’), and the allele is shown after the equal sign. Present (yellow) indicates the expected change in enhancer activity for oligos that have the allele denoted on the y-axis, while absent (purple) shows the expected change for oligos that lack the allele. Yellow points at positive impact on RNA/DNA values means that variant or variant combination increases enhancer activity on average across oligos with other variant combinations, while purple points at positive values mean the variant or variant combination decreases activity (i.e., activity is higher when absent). (**C**) 2xHAR.170 is a candidate intronic enhancer of *GALNT10*, acquiring a human-specific change from C to T that enhances POU4F1 and TEF binding in our footprint analysis. The human polymorphism rs2434531 enhances binding of the repressor HMX1. The HMX1 and TEF footprints were detected in an independent brain footprinting study (Funk et al., 2020). Both results are supported by Sei predictions (**4E**), MPRA activity (**A**), and differential activity between the chimpanzee (**D**) and human (**E**) sequences of 2xHAR.170 in the forebrain and midbrain of transgenic mouse embryos. Adapted from (Capra et al., 2013).

We next dissected variant effects in individual HARs, focusing on 2xHAR.170 as it showed strong interactions between its three human variants (**Figure 7B**). The chimpanzee haplotype (CTT) is near the middle of activity levels across all permutations, with some variant combinations decreasing and others increasing activity up to 24%. The derived C allele at the third variant has the largest individual effect in lentiMPRA, which is consistent with it having the largest effect of the three 2xHAR.170 variants in our Sei analysis (**Figure 7A**). Footprint analysis (**Figure 7C**) shows that this C decreases binding affinity of HMX1 (Furlan et al., 2013), a repressor of neural differentiation-driver TLX3 (Divya et al., 2016), consistent with increased enhancer activity. Since the third variant is polymorphic, some humans have the least active permutation (log2(RNA/DNA)=0.0184) while others have one of the most active (log2(RNA/DNA)=0.169), with possible phenotypic consequences (see Discussion). In contrast, the derived alleles at the other two variants individually decrease enhancer activity in lentiMPRA and Sei analysis and have the lowest activity when tested together in combination with the chimpanzee T allele at the third variant. This is consistent with their being proximal and changing high information content positions in a POU4F1 footprint, which is supported by POU4F1 ChIP-seq in fibroblasts (Hammal et al., 2022). However, the activity-increasing effect of the derived C allele at the third variant is amplified, not reduced, in the presence of the derived T allele at the first variant. Thus, the human-specific variants in 2xHAR.170 clearly have interacting effects on brain enhancer activity.

## DISCUSSION

In this work, we used ML models coupled with lentiMPRA and epigenetic (ChIP-seq, ATAC-seq, Hi-C) experiments in chimpanzee and human NPCs, dozens of which we generated, to functionally profile 714 HARs at single-nucleotide resolution in neurodevelopment. We discovered a much greater effect on enhancer activity for HAR sequence variants as compared to species-specific differences in the cellular environment. We showed that this *cis* regulatory divergence can be predicted using differences in TF footprints within HARs. Finally, we dissected the contribution of all nucleotide variants in each HAR, revealing pervasive interactions between sites, in many cases suggestive of compensatory evolution to maintain ancestral enhancer activity. Altogether, our results prioritize dozens of HARs with evidence of differential neurodevelopmental enhancer activity in humans compared to chimpanzees and other mammals.

The key novelty of our study is our use of ML modeling to efficiently integrate hundreds of epigenetic datasets and screen thousands of HAR variants *in silico.* To our knowledge, this study is the first to use ML to characterize HARs at the level of single nucleotides and their interactions. Epigenomic data has been used previously, through overlap analyses (Girskis et al., 2021; Uebbing et al., 2021) and supervised learning (Capra et al., 2013), to predict which HARs function as developmental enhancers and which ones are active in the developing brain. We significantly extended these prior analyses in four ways. First, we used hundreds of epigenetic data sets from primary tissues, organoids and cell lines, many generated in this study or in the past few years, which vastly expanded the features available for ML modeling. Second, we performed supervised learning of these epigenetic features for HARs alongside *in vivo* validated enhancers, which showed clear clustering of a subset of HARs with tissue-specific enhancers.

Third, we leveraged methods for transcription factor footprinting to model how human-specific variants in HARs alter enhancer activity. Many prior studies looked at how HAR sequence variants changed transcription factor binding motifs. But footprinting based on our chimpanzee NPC H3K27ac ChIP-seq provided a more precise measurement of which transcription factors bind human versus chimpanzee HAR sequences, which in turn allowed us to very accurately predict species-biased enhancer activity in lentiMPRA. This highlighted specific transcription factors with the largest roles in differential enhancer activity and generated testable hypotheses about HAR variants with causal roles in enhancer evolution.

Finally, we used ML to dissect HARs at the single-nucleotide level through the deep-learning framework Sei. Using this tool for predicting how variants change the epigenetic state of genome sequences, we found evidence that individual variants within the same HAR often have opposite effects on enhancer activity. This prompted us to perform a lentiMPRA experiment to test all combinations of variants in each of a small set of HAR enhancers, revealing both negative and positive interactions. With lentiMPRA and CRISPRa we directly measured activity for a small number of HARs in NPCs, whereas Sei and gradient boosting enabled us to screen and prioritize *in silico* many more HAR variants with much lower cost and effort. Thus, ML is highly complementary to experimental methods, and together they advanced understanding of HAR function.

Our lentiMPRA experimental design was unique compared to prior HAR MPRA studies (Girskis et al., 2021; Uebbing et al., 2021), because we used chimpanzee NPCs alongside human ones. This enabled us to disentangle *trans* effects of the cellular environment from *cis* effects of sequence variants in lentiMPRAs, leading to our discovery that *cis* effects are common and highly consistent between human and chimpanzee cells, as well as two stages of neural differentiation (N2 and N3). This result is concordant with prior literature on *in vivo* enhancer assays being sensitive to sequence changes but robust for a given sequence tested across different vertebrate species (Mattioli et al., 2020; Ritter et al., 2010). Another benefit of using chimpanzee NPCs was our ability to generate epigenomic data in both species, which enabled our comparative transcription factor footprint analyses and generation of a ML model that predicts *cis* effects in HAR enhancers from human and chimpanzee transcription factor footprints, pointing to the importance of differential regulators binding HARs in driving their species-biased enhancer activity. This compendium of chromatin state and interaction data, collected in parallel for chimpanzee and human NPCs (**Table 1**), is freely available to enable future ML modeling.

While our lentiMPRA was highly reproducible in NPCs, it did not perfectly agree with our ML model trained on epigenetic profiles and VISTA brain enhancers. This is consistent with observations that only 26% of ENCODE enhancer predictions based on epigenetic marks in K562 cells validated in MPRAs (Kwasnieski et al., 2014) and that fruit fly enhancers identified via epigenetic marks and MPRA activity are largely non-overlapping (Lindhorst and Halfon, 2022). Indeed, such differences are expected. MPRAs test sequences outside their native chromatin context, often using insulators, and therefore tend to be permissive (i.e., reporting potential regulatory activity). Conversely, epigenetic marks typically associated with enhancers do not alone indicate enhancer function. The two approaches are also affected by differences between our lentiMPRA and the reporter assays in VISTA, such as measuring activity in NPCs versus whole embryonic brains, primate versus mouse cells, *in vivo* versus *in vitro* reporter assays, and ∼100-bp versus ∼1,000-bp sequences. Despite these differences, ML and lentiMPRA consistently prioritized dozens of HARs as neurodevelopmental enhancers, and each approach likely revealed some HAR enhancers missed by the other method.

Iteratively combining ML and experimentation shed light on a major question regarding HARs: why did they acquire so many mutations in the human lineage after being conserved throughout mammalian evolution? This question has been hard to tackle, because most HAR sequence changes occurred before our common ancestor with Neanderthals and other archaic hominins (Hubisz and Pollard, 2014), which means we cannot directly link sequence changes to phenotypes. We hypothesized that functional data for individual variants and variant combinations in both human and chimpanzee cells would provide some insight. For example, if a human-specific variant changed enhancer activity relative to the chimpanzee allele in chimpanzee cells but not human cells, we might conclude that it evolved to maintain ancestral activity in the presence of altered trans factors in human cells. We found little evidence for such trans effects and instead concluded that species-biased HARs are driven primarily by sequence changes leading to differential use of regulatory pathways and proteins present in NPCs from both species. While investigating effects of individual variants versus variant combinations, we expected that variants within the same HAR would alter enhancer activity in the same direction, potentially interacting synergistically to generate large differences between the human and chimpanzee reference genome sequences. In contrast, we found that variants in the same HAR have both positive and negative interactions, and often some variants individually increase activity while others decrease it. This suggests the possibility that compensatory evolution played a role in the rapid evolution of HARs. Combining these results, we speculate that a typical HAR enhancer may have evolved through initial variants with large changes in activity that were then moderated back towards ancestral levels by subsequent nearby variants.

Our analysis of the genetic and three-dimensional chromatin interactions between HARs and genes provides some insight into why HARs might have evolved in this forward-and-back way. Using our NPC Hi-C data and publicly available brain Hi-C data, we showed that most HARs with enhancer activity in NPCs have 3D chromatin interactions with neurodevelopmental genes, often across multiple brain cell types. Furthermore, many HAR variants are in linkage disequilibrium with QTLs, establishing a connection to differential gene expression and chromatin accessibility. Finally, many active HARs are genetically linked to neuropsychiatric disease SNPs. These results suggest that differential enhancer activity of the HAR could affect brain development and phenotypes. It has been suggested that changes in the human brain that enable our unique cognitive abilities are “Achilles’ heels” that also contribute to schizophrenia and other psychiatric disorders (Crow, 2000). Thus, it is plausible that opposing selection pressures for new cognitive traits and against neurological disease were amongst the evolutionary forces that contributed to interacting, compensatory variants in HAR enhancers and their many sequence differences between humans and chimpanzees.

Through our integrative analyses, 2xHAR.170 emerged as a HAR that could have accelerated through compensatory evolution. We do not know the order in which the three variants in 2xHAR.170 occurred. One intriguing possibility is that the third variant increased activity so much that there was strong selection to reduce activity back towards the ancestral level, leading to the first and second variants. While this is possible, the third variant is polymorphic (rs2434531), suggesting that it may be the most recent. This polymorphism is an eQTL for *GALNT10* in lymphoblastoid cells, where the derived allele is associated with higher expression (Wu et al., 2020). This upregulation is directly relevant to neurological disease, because *GALNT10* is overexpressed in individuals with schizophrenia (Voisey et al., 2017). In neuronal cells, 2xHAR.170 is bound by the transcription factor FOXP2, as well as other enhancer-associated proteins (ISL1, HAND2, PHOX2B, FOSL2) and chromatin modifiers (EZH2, SMARCA2, SMARCC1) (Boyle et al., 2012). These connections to *GALNT10* and neuronal function support an important role for 2xHAR.170 in regulating neurodevelopment and are consistent with compensatory evolution in this HAR.

No doubt the true evolutionary trajectory of HARs is more complex than this one hypothesis. It is also likely that our conclusions are influenced by biases in currently available epigenomic data and in reporter technologies. For instance, MPRA, luciferase and transgenic animal enhancer assays test candidate enhancers outside their native locus and its chromatin environment. Therefore, it is possible that HAR enhancers in their genomic loci would show trans effects that we could not detect in this study. This is an important direction for future work. Another caveat is that our NPCs represent only one cell type and two differentiation time points, whereas our unsupervised learning analysis indicates that HARs likely function broadly across tissues, cell types, and developmental stages. In this study, we addressed the shortcoming that prior HAR MPRA and epigenetic studies used human cells, and we created an *in vitro* system to assay HARs in chimpanzee neurodevelopment. But it is entirely possible that HAR variants are driven by functional effects in other cellular contexts. Our Sei analysis revealed examples where a variant has differential effects across tissues, although most HAR variants do not show this pattern. Nonetheless, it will be critical to evaluate how human-specific sequence changes affect HAR enhancer function beyond early neuronal differentiation. The integrated ML and experimental strategy presented here provides a framework for these investigations.

## SUPPLEMENTARY FIGURE CAPTIONS

**Figure S1. ChIP-seq signal across cell lines at HARs and genome-wide.** (a-e) Normalized ChIP-seq read coverage (CPM) at HARs. For each HAR, reads are centered on the position with maximum coverage. Rows in all panels are ordered by coverage in panel a. One representative replicate is shown for each condition. (a) H3K27ac in human N2, (b) H3K27ac in human N3, (c) H3K27me3 in human N2, (d) H3K27ac in chimpanzee N2, (e) H3K27ac in chimpanzee N3. (f-g) Normalized read coverage (CPM) genome-wide at transcription start sites (TSS). Rows in both panels are ordered by coverage at the TSS (offset 0) in panel f. All TSS shared between the human and chimpanzee genomes are analyzed. (f) H3K27ac in human N2, (g) H3K27ac in chimpanzee N2.

**Figure S2. Overlap of HARs with epigenetic datasets for heart, brain, and other primary tissues.**

**Figure S3. Correlation of Sei-predicted chromatin state changes across all human-chimpanzee HAR variants.**

**Figure S4. LentiMPRA study design.**

**Figure S5. Transgenic mouse embryos for 2xHAR.133, HAR152, 2xHAR.518 and 2xHAR.548.** Pictures of all PCR positive embryos from mouse transgenic enhancer assays. The name of the tested HAR is noted at the top left of each set of images, and the sequence origin (human or chimpanzee) is given in each row. Tissue names are abbreviated as notochord (NT), forebrain (FB), midbrain (MB), limb (LB), and facial mesenchyme (FM). A plus sign (+) in the row of the tissue name means LacZ expression was observed for that tissue. Embryos that did not show LacZ expression but were LacZ positive by PCR, are marked as negative (NEG).

**Figure S6. Luciferase assays for validation of lentiMPRA.** (a-b) Relative luciferase activity in human (HS1, panel a) and chimpanzee (Pt2a, panel b) N3 cells for human (blue) or chimpanzee (green) HAR sequences. We selected HARs across a range of RNA/DNA levels (mean of human and chimpanzee alleles) and tested whether relative activity of the human and chimpanzee alleles was similar in luciferase assays compared to lentiMPRA. 2xHAR.273, 2xHAR.434, 2xHAR.11, 2xHAR.417, and 2xHAR.176 (bold font) had higher activity of the human allele in lentiMPRA. 2xHAR.518, 2xHAR.401, 2xHAR.35, 2xHAR.53, and 2xHAR.364 (plain font) had higher activity of the chimpanzee allele in lentiMPRA. 2xHAR.401, 2xHAR.518, 2xHAR.176, 2xHAR.417, 2xHAR.11, and 2xHAR.173 each have at least one allele (human and/or chimpanzee sequence) significantly higher than the empty vector. We also tested seven “inactive” sequences with low activity in lentiMPRA (neg, N01, N06, N10, N12, N15, N17), empty pLS-mP-luc vector (Empty), and pLS-SV40-mP-luc (SV40). Asterisks indicate statistically significant differences between the human and chimpanzee alleles by Student’s t-test. (c-e) Relative activity of human versus chimpanzee alleles for HARs from (a-b) that showed species-biased activity in lentiMPRA. (c) HS1 luciferase, (d) Pt2a luciferase, and (e) lentiMPRA.

**Figure S7. LentiMPRA activity of control sequences.** RNA/DNA ratios of negative (H3K27me3, red) and positive (H3K27ac, blue) controls by cell line (columns) and cell stage (rows). The activity distribution is significantly higher for positive controls in all panels.

**Figure S8. UMAP plot of HARs co-embedded with validated VISTA enhancers.** Each HAR and VISTA enhancer is described by overlaps with epigenetic datasets in primary brain, heart, and limb tissue and co-embedded in UMAP space. Validated non-brain (primarily heart and limb) enhancers cluster in the upper right corner, while brain enhancers cluster in the lower left. A ML model (L1-penalized classifier) was trained to distinguish HARs with high lentiMPRA activity (top 25%) and low ML scores (bottom 25%) from those with low lentiMPRA activity (bottom 25%) and high ML scores (top 25%). The model achieved high accuracy (0.96 auPR), indicating that these groups of discordant HARs have distinct combinations of epigenetic features. Examples of several of the most important epigenetic features are shown in panels a-d. The most discordant HARs (top quartile with only one method) that overlap each feature are colored to indicate if the HAR has a high score in lentiMPRA only (red) or ML only (blue). (a-b) HARs with high lentiMPRA activity but low ML scores have marks of active chromatin in non-brain tissues. These are a mix of enhancers with active marks in NPCs and other tissues, plus enhancers inactive in NPCs that nonetheless show high activity in lentiMPRAs due to being tested outside their native chromatin environment that is silent in the embryonic brain. (a) Embryonic (Day 81) forelimb DNase-seq. (b) Adult (46 years) heart ventricle H3K27ac. (c-d) HARs with high ML scores but low lentiMPRA activity overlap open chromatin in samples from the whole fetal brain. These appear to be mostly embryonic brain enhancers active in cell types other than forebrain neurons. (c) Embryonic (Day 58) brain DNase-seq. (d) Embryonic (Day 72) brain DNase-seq.

Figure S9. LentiMPRA activity measurements are highly reproducible across technical and biological replicates. Correlation scatterplot of RNA/DNA ratios for all combinations of cell line, cell stage, and replicate.

**Figure S10. Heatmap of lentiMPRA activity for all species-biased HARs.** Chimpanzee and human sequences of the same HAR are plotted next to each other. The stripes present across all cell lines, cell stages, and replicates show consistent differences in activity between the alleles from the two species.

**Figure S11. Associating active HARs with neurodevelopmental genes.** Number of active HARs (lentiMPRA) interacting with GWAS variants via chromatin loops, interacting with QTLs via chromatin loops, sharing an LD block with GWAS variants, sharing an LD block with QTLs, interacting with the promoter of protein coding genes via chromatin loops, and sharing a contact domain (sub-TAD) with promoters of protein-coding genes.

**Figure S12. LentiMPRA activity of HAR permutations.** Activity of all evolutionary intermediates for (a) 2xHAR.164 and (b) 2xHAR.238. Human bases of the allele are shown in blue and chimpanzee bases in green. Single nucleotide changes are colored with the predicted brain enhancer state change from Sei, showing moderate concordance with lentiMPRA activity.

**Figure S13. Permutation lentiMPRA measurements are highly concordant.** We observed high correlation between lentiMPRA activity levels of permutation oligos in library 1 and library 2 after batch correction. Correlation is high across all cell lines, cell stages, and replicates.

**Figure S14. Distribution of the number of 171-bp oligos required to tile each HAR.**

## SUPPLEMENTARY TABLE CAPTIONS

Table S1. Single-cell RNA-seq cell counts.

Table S2. Transcription factor footprints called in each HAR using the human genome and human H3K27ac or the chimpanzee genome and chimpanzee H3K27ac, as well as footprints disrupted by human:chimpanzee variants.

Table S3. In vivo epigenetic datasets used in this study.

Table S4. Telencephalon expression in HARs characterized in published mouse enhancer assays.

Table S5. Sei predicted epigenetic state changes for each HAR variant. Table S6. Gene set enrichment analysis results using gProfiler.

Table S7. Annotations of HARs active in lentiMPRA. Table S8. LentiMPRA oligonucleotide library.

Table S9. Primers used in this study.

## METHODS

### Cell Lines

We performed lentiMPRA in N2 and N3 cells derived from four separate iPSC lines from two human and two chimpanzee males. All lines were reprogrammed from fibroblasts using episomal plasmids according to a recently published protocol(Okita et al., 2013). One iPSC line was previously described (WTC;(Miyaoka et al., 2014)), and three were generated from low passage fibroblasts (P3 – P7) from Coriell Cell Repository (Hs1: 2 year old human male, catalog AG07095; Pt2: 6 year old chimpanzee male, Maverick, catalog: S003611; Pt5: 8 year old chimpanzee male, catalog PR00738)(Pollen et al., 2019). We electroporated three micrograms of episomal expression plasmid mixture encoding OCT3/4, SOX2, KLF4, L-MYC, LIN28, and shRNA for TP53 into 300,000 fibroblasts from each individual with a Neon Electroporation Device (Invitrogen), using a 100 μL kit, with setting of 1,650V, 10ms, and three pulses(Bershteyn et al., 2017; Pollen et al., 2019). After 5–8 days, cells were detached and seeded onto irradiated SNL feeder cells. The culture medium was replaced the next day with primate ESC medium (Reprocell) containing 5 – 20 ng/mL of βFGF. Colonies were picked after 20 – 30 days, and selected for further cultivation. After three to five passages, colonies were transferred to Matrigel-coated dishes and maintained in mTeSR1 medium (Stem Cell Technologies, 05850) supplemented with Penicillin/Streptomycin/Gentomycin. Further passaging was performed using calcium- and magnesium-free PBS to gently disrupt colonies. Each line showed a normal karyotype, and was recently described(Pollen et al., 2019). The UCSF Committee on Human Research and the UCSF GESCR (Gamete, Embryo, and Stem Cell Research) Committee approved all human iPSC experiments. HepG2 cells were cultured in Dulbecco’s Modified Eagle Medium (Corning) supplemented with 10% FBS and Penicillin-Streptomycin, and passaged every 4-5 days using StemPro Accutase (Thermo Fisher Scientific).

### Neural differentiation of human and chimpanzee iPSCs

Human and chimpanzee iPSCs were cultured in Matrigel-coated plates with mTeSR media in an undifferentiated state. Cells were propagated at a 1:3 ratio by treatment using calcium and magnesium free PBS to gently disrupt colonies by mechanical dissection. To trigger neural induction, iPSCs were split with EDTA at 1:5 ratios in culture dishes coated with matrigel and culture in N2B27 medium (comprised of DMEM/F12 medium (Invitrogen) supplemented with 1% MEM-nonessential amino acids (Invitrogen), 1 mM L-glutamine, 1% penicillin-streptomycin, 50 ng/mL bFGF (FGF-2) (Millipore), 1x N2 supplement, and 1 x B27 supplement (Invitrogen)) supplemented with 100 ng/ml mouse recombinant Noggin (R&D systems). N1 cells were collected eleven days after initiating neural induction. Cells at passages 1-3 were split by collagenase into small clumps, similar to iPSC culture, and continuously cultured in N2B27 medium with Noggin. After passage 3, cells were plated at the density of 5E4 cells/cm^2^ after disassociation by TrypLE express (Invitrogen) into single-cell suspension, and cultured in N2B27 medium supplemented with 20 ng/mL bFGF and EGF. Cells were maintained under this culture condition for a minimum of three months with a stable proliferative capacity. N2 cells were collected at P12-18 and N3 cells at P20-28.

### Validation of N2 and N3 markers through immunostaining

Human and chimpanzee N2 and N3 cells were examined using immunostaining against neural and glial progenitor markers. Cells were cultured in chambered Millipore EZ slides, rinsed with PBS, fixed with 4% paraformaldehyde in PBS for 15 minutes at room temperature, washed three times with ice cold PBS, and permeabilized through incubation for 10 min with PBS containing 0.1% Triton X-100. Cells were washed in PBS three times and incubated with 10% donkey serum for 30 minutes to block nonspecific binding of antibodies. Cells were next incubated with diluted primary antibodies against Nestin (monoclonal mouse, Abcam, AB6142), Pax6 (polyclonal rabbit, Abcam, AB5790), and GFAP (polyclonal rabbit, Chemicon, AB5804) in 10% donkey serum for 1 hour at room temperature. The cells were then washed three times in PBS, 5 minutes each wash, then incubated with a secondary antibody (Alexa 488 donkey anti rabbit, Life technologies; Alexa 546 donkey anti mouse, Life technologies) in donkey serum for 1 hour at room temperature in the dark. Cells were then washed three times with PBS in the dark, then covered with a coverslip in Cytoseal mounting media (Thermo Scientific).

### Single Cell RNA-Sequencing

To determine the composition of cell types in human and chimpanzee cell lines used for lentiMPRA, we generated single cell gene expression (scRNA-seq) data and clustered cells from each line based on expression. Cells were captured using the C1TM Single-Cell Auto Prep Integrated Fluidic Circuit (IFC), which uses a microfluidic chip to capture the cells, perform lysis, reverse transcription and cDNA amplification in nanoliter reaction volumes. The details of the protocol are described in PN100-7168 (http://www.fluidigm.com/). Sequencing libraries were prepared after the cDNA was harvested from the C1 microfluidic chip using the Nextera XT Sample Preparation Kit (Illumina), following its protocol with minor modifications. The single cell libraries from each C1 capture were then pooled, cleaned twice with 0.9X Agencourt AMPure XP SPRI beads (Beckman Coulter), eluted in DNA suspension buffer (Teknova) or EB buffer (Qiagen) buffer and quantified using High Sensitivity DNA Chip (Agilent). scRNA-seq paired-end reads were generated for ∼50 cells per library (**Supplemental Table S1**). Sequencing data is available through the accession number GSE110760 (chimpanzee cells: GSE110759, human cells: GSE110758). We trimmed reads for quality using cutadapt under the Trim Galore! wrapper (https://www.bioinformatics.babraham.ac.uk/projects/trim_galore/) with the default settings, and Nextera transposase sequences were removed. Reads shorter than 20 bp were discarded. Read level quality control was then assessed using FastQC (http://www.bioinformatics.babraham.ac.uk/projects/fastqc/). Reads were aligned to the NCBI human reference assembly GRCh38 by HiSat2(Kim et al., 2015) using the prefilter-multihits option and a guided alignment via the human Gencode Basic v20 transcriptome. Expression for RefSeq genes was quantified by the featureCounts routine, in the subRead library(Liao et al., 2013), using only uniquely mapping reads and discarding chimeric fragments and unpaired reads. Gene expression values were normalized based on library size as counts per million reads (CPM). We used visual image calls to remove any libraries that originated from C1 chambers with multiple cells. To further identify outlier cells, we removed those with fewer than 1,000 genes detected, or with greater than 20% of reads aligning to mitochondrial or ribosomal genes. Gene expression was analyzed using a threshold of detection for each gene at 2 CPM. We then calculated the percentage of cells expressing regional identity genes (e.g., *FOXG1* for telencephalon, *DLX6-AS1* for GABAergic neurons, *MKI67* for dividing cells, *SLC1A3* for radial glia). In both human and chimpanzee cell lines at the NPC and GPC stage, 50-90% of cells expressed telencephalon (*FOXG1*) and radial glia/astrocyte markers.

### Histone ChIP-seq experiments

Ten million human (HS1) and chimpanzee (Pt2a) N2 and N3 cells were crosslinked with 1% formaldehyde for 5 minutes and quenched with 125 mM glycine for 5 minutes. To obtain antibody-beads conjugate, Dynabeads protein A (Invitrogen) and Dynabeads protein G (Invitrogen) were mixed at 1:1 ratio and washed twice with Buffer A (LowCell# ChIP kit, diagenode). 10 μg of H3K27ac antibody (Abcam, ab4729) was added to the beads, and gently agitated at 4°C for 2 hours. ChIP was performed using LowCell# ChIP kit (Diagenode) according to manufacturer’s protocol. Sequencing libraries were generated using Accel-NGS 2S Plus DNA library kit (Swift Biosciences). DNA was quantified with Qubit DNA HS assay kit and Bioanalyzer (Agilent) using the DNA High Sensitivity kit. Sequencing was performed using an Illumina HiSeq 4000 with 50 bp single reads. Two biological replicates were done for each cell type. ChIP-seq was processed by the ENCODE Transcription Factor and Histone ChIP-seq processing pipeline (https://github.com/ENCODE-DCC/chip-seq-pipeline2) using default parameters. The pipeline configuration file was modified to enable alignment of Pt2a cell line data to the panTro5 genome.

### ATAC-seq experiments

ATAC-seq was performed according to previously described(Inoue et al., 2019). Briefly, 50,000 cells were dissociated using Accutase and precipitated with centrifugation at 500 g for 5 minutes. The cell pellet was washed with PBS, resuspended in 50 mL lysis buffer (10 mM TrisHCl, pH 7.4, 10 mM NaCl, 3 mM MgCl2, 0.1% Igepal CA-630), and precipitated with centrifugation at 500 g for 10 minutes. The nuclei pellet was resuspended in 50 mL transposition reaction mixture which includes 25 μL Tagment DNA buffer (Nextera DNA sample preparation kit; Illumina), 2.5 μL Tagment DNA enzyme (Nextera DNA sample preparation kit; Illumina), and 22.5 μL nuclease-free water, and incubated at 37C for 30 minutes. Tagmented DNA was purified with MinElute reaction cleanup kit (QIAGEN). The DNA was size-selected using SPRIselect (Beckman Coulter) according to the manufacturer’s protocol. 0.6x and 1.5x volume of SPRIselect was used for right and left side selection, respectively. Library amplification was performed as previously described(Buenrostro et al., 2013). Amplified library was further purified with SPRIselect as described above. DNA was quantified on a Bioanalyzer using the DNA High Sensitivity kit (Agilent). Massively parallel sequencing was performed on an Illumina HiSeq4000 with PE150. ATAC-seq was done in 2 biological replicates for each time point.

### Hi-C experiments

Hi-C was performed using the Arima Hi-C kit (Arima Genomics) according to the manufacturer’s instructions. 10 million cells were used. The sequencing library was prepared using Accel-NGS 2S Plus DNA Library Kit (Swift Biosciences) according to the manufacturer’s protocol. Two independent biological replicates were prepared for each cell line. In total eight libraries were pooled and sequenced with paired-end 150-bp reads using two lanes of a NovaSeq6000 S2 (Illumina) at the Chan Zuckerberg Biohub.

### Transcription factor footprints

While traditional footprinting methods operate on open chromatin data, the HINT (Gusmao et al., 2014) method can also compute footprints from histone ChIP-seq data alone, albeit with reduced accuracy. We utilized this strategy, because we have matched H3K27ac ChIP-seq from human and chimpanzee NPCs. Genome-wide footprint analysis was run separately on human N2 H3K27ac ChIP-seq and chimpanzee N2 H3K27ac ChIP-seq, both using HOCOMOCO v11 and JASPAR 2020 TF motifs. Expressed TFs were defined as those with TPM>1 in NPCs (Schwartz et al., 2015) (Kallisto (Bray et al., 2016) v0.48). HINT (v0.13.2) was used to compute enrichment of footprints within HARs compared to H3K27ac peaks (‘rgt-hint footprinting --histonè followed by ‘rgt-motifanalysis matching‘).

### LentiMPRA library design

All HARs from our prior studies (Lindblad-Toh et al., 2011; Pollard et al., 2006a) that were not fully covered in the human (hg19) and chimpanzee (panTro2) reference genome sequences were included in the library design. These 714 HARs have similar lengths and genomic distributions compared to the larger set of 2645 HARs, so we expect them to be representative. For each HAR, we designed 171-bp oligos representing the orthologous human and chimpanzee sequences. Since HARs have median length 227 bp, most could be synthesized using a single oligo or two highly overlapping oligos. Flanking genomic sequence was added to HARs shorter than 171 bp, and HARs longer than 171 bp were tiled with multiple oligos having variable but considerable overlap depending on the length of the HAR (e.g., 67% overlap between the two oligos for HAR sequences between 171 and 342 bp long, which have mean length 233 bp). A third of HARs could be synthesized using a single oligo (**Supplemental Figure S14**). For the remaining HARs, the multiple oligos were separately quantified for enhancer activity (see below) and assessed for agreement. We observed high correlation between multiple oligos per HAR, likely due to their generally high level of overlap, so we merged all oligos per HAR (summed their reads) for downstream analysis. This produced one activity measurement for each human or chimpanzee HAR sequence.

We additionally synthesized 118 sequences that we expected would show little or no enhancer activity in NPCs (negative controls) and 143 sequences that we expected would drive expression in NPCs (positive controls). Negative controls were comprised of 34 sequences used as negatives by the ENCODE consortium (provided by Rick Meyers) plus 84 human genome sequences located in H3K27me3 ChIP-seq peaks from human N2 and N3 cells (data generated in the Ahituv lab, released with this study). Positive controls included 9 positive enhancer elements from ENCODE (provided by Rick Meyers), 124 human genome sequences located in H3K27ac ChIP-seq peaks from human N2 and N3 cells (data generated in the Ahituv lab, released with this study), and 10 human genome sequences we predicted would function as neurodevelopmental enhancers using our EnhancerFinder algorithm (Erwin et al., 2014).

All HAR and control sequences were scanned for restriction sites (for *Sbf*I and *Eco*RI) and modified to avoid problems in synthesis and cloning. We designed our experiments to ideally have 100 unique 15-bp barcodes per variant to build in robustness to barcode dropout and jackpotting issues, variability in activity across integration sites, and other sources of technical error. These barcodes are not random, but rather designed to be at least two substitutions and one insert-deletion (indel) apart from other barcodes and synthesized with the oligos. The final array design included 2,440 unique 171-bp sequences, each with 100 barcodes, for a total of 244,000 oligos inclusive of HARs and controls (**Supplemental Table S8**).

### LentiMPRA library synthesis and cloning

All lentiMPRA sequences were array-synthesized as 230-bp oligos (Agilent Technologies) containing universal priming sites (AGGACCGGATCAACT…CATTGCGTGAACCGA), a 171-bp candidate enhancer sequence, spacer (CCTGCAGGGAATTC), and 15-bp barcode. The amplification and cloning of the enhancers and barcodes into the pLS-mP lentiviral vector was performed as previously described(Inoue et al., 2017). Briefly, pLS-mP was cut with *Sbf*I and *Eco*RI taking out the minimal promoter and EGFP reporter gene. The oligos containing the HAR, spacer, and barcode were amplified with adaptor primers (pLSmP-AG-f and pLSmP-AG-r; **Supplemental Table S9**) that have overhangs complementary to the cut vector backbone, and the products were cloned using NEBuilder HiFi DNA Assembly mix (NEB, E2621). The cloning reaction was transformed into electro-competent cells (NEB C3020) and multiple transformations were pooled and midiprepped (Chargeswitch Pro Filter Plasmid Midi Kit, Invitrogen CS31104). The library was then cut using *Sbf*I and *Eco*RI sites contained within the spacer, so that the minimal promoter and EGFP could be reintroduced via a sticky end ligation (T4 DNA Ligase, NEB M0202). This library was transformed and purified, as previously described, and DNA sequenced to determine complexity.

We estimate that at least 92% of barcodes are correctly synthesized and that per-base substitution errors are about 0.02-0.04% (synthesis and amplification). The observed median number of unique barcodes per variant ranged from 79 to 81 across our 24 replicates, with a 25th percentile of 73 to 76 barcodes and a 75th percentile of 79 to 87 barcodes. The chimpanzee sequence of 2xHAR.335 had particularly low barcode counts, with a median of 17, a minimum of 16, and a maximum of 18. The next worst HAR has a median of 49 barcodes. Thus, barcode count was consistently high for most of the HARs we tested. Also, the number of barcodes was similar across replicates for any given oligo, suggesting synthesis as the source of differential numbers of barcodes. By aiming for 100 barcodes per variant, we generated a library with high numbers of barcodes despite barcode dropout.

### Lentivirus library preparation and infection

Lentivirus packaging of the HAR lentiMPRA library was performed by the UCSF Viracore using standard techniques (Wang and McManus, 2009). Twelve million HEK293T cells were plated in a 15-cm dish and cultured for 24 hours. The cells were co-transfected with 8 μg of the HAR library and 4 μg of packaging vectors using jetPRIME (Polyplus-transfections). The transfected cells were cultured for 3 days and lentiviruses were harvested and concentrated as previously described (Wang and McManus, 2009). For all human and chimpanzee cell lines and cell stages, about twelve million cells were plated in 15-cm dishes and cultured for 24-48 hours. Cells were infected with a multiplicity of infection (MOI) of 50. When infecting the library into HepG2 cells, 8 μg/mL polybrene was added to the cells. The culture medium was refreshed daily. Infected cells were washed with PBS three times before cell lysis in order to remove any non-integrated lentivirus.

### RNA & DNA isolations and sequencing

Genomic DNA and total RNA were extracted using the AllPrep DNA/RNA mini kit (Qiagen). Messenger RNA was purified from the total RNA using Oligotex mRNA mini kit (Qiagen) and treated with Turbo DNAseq to remove contaminating DNA. The RT-PCR, amplification and sequencing of RNA and DNA were performed as previously described(Inoue et al., 2017), with some alterations for adding Unique Molecular Identifiers (UMIs) in the process. In brief, mRNA was reverse transcribed with SuperScript II (Invitrogen) using a primer downstream from the barcode (pLSmP-ass-R-UMI-i#; **Supplemental Table S9**). The resulting cDNA was split into multiple reactions to reduce PCR jack-potting effects and cDNA amplification performed with Kapa Robust polymerase for three cycles, incorporating unique molecular identifiers (UMIs) of 10 bp length. PCR products were cleaned with AMPure XP beads (Beckman Coulter) to remove primers and concentrate samples. These products underwent a second round of amplification in 8 reactions per replicate for 15 cycles, switching from the UMI-incorporating reverse primer to one containing only the P7 flow cell sequence (P7; **Supplemental Table S9**). All reactions were pooled and run on agarose gels for size selection and submitted for sequencing. For DNA, each replicate was amplified for 3 cycles with UMI-incorporating primers, just as the RNA. First round products were cleaned up with AMPure XP beads, and amplified in split reactions, each for 20 cycles. Again, reactions were pooled and gel-purified.

RNA and DNA for all three replicates for all samples were sequenced on an Illumina NextSeq instrument (2x15 bp barcodes + 10bp UMI + 10bp sample index) using custom primers (BARCODE-SEQ-R1-V4, pLSmP-AG-seqIndx, BARCODE-SEQ-R2-V4; **Supplemental Table S9**) and are available through the Short Read Archive (SRA) with BioProject accession numbers PRJNA428580 (chimpanzee cells) and PRJNA428579 (human cells). Illumina Paired End reads sequenced the barcodes from the forward and reverse direction and allowed for adapter trimming and consensus calling of tags (Kircher, 2012). Barcode or UMI sequences containing unresolved bases (N) or not matching the designed length of 15 bp were excluded. In data analysis, each barcode x UMI pair is counted only once and only barcodes matching perfectly to those included in the above oligo design were considered.

### RNA/DNA ratios and quantification of enhancer activity

RNA and DNA counts were first normalized per replicate using counts per million reads mapped (CPM). RNA/DNA ratios per HAR per replicate were calculated by taking the sum of RNA counts for all ∼80 barcodes assigned to all oligo(s) tiling across each HAR, divided by the sum of all DNA counts for all barcodes across all oligo(s) per HAR, and using only barcodes with >0 counts in DNA. Importantly, we do not compute RNA/DNA for each barcode and average these, but rather use the ratio of the sum of RNA counts and the sum of DNA counts over all detected barcodes, which is more robust to over-represented (PCR “jackpot”) barcodes than first taking the ratio per barcode. We also tried using the ratio of the median RNA and median DNA count, rather than sums (equivalent to means when RNA and DNA have the same number of barcodes), and we observed a correlation of ∼98.5%, demonstrating that our quantification method is indeed robust. We summed counts across oligos for HARs tiled using two or more oligos, because we observed generally good agreement between oligos for the same HAR. The resulting RNA/DNA ratios were batch normalized for RNA and DNA library preparation date using limma (Ritchie et al., 2015).

We focused our differential activity analyses on HARs with the highest and most consistent activity across replicates, specifically, 293 “active” HARs (41%) that drive expression above the median of positive controls in at least 50% of samples for either the human or chimpanzee sequence. These HARs also all have activity above the 75th percentile of the negative controls in at least 50% of samples for either the human or chimpanzee sequence. There is no threshold that perfectly separates positive and negative controls, because their activity distributions overlap despite positives being significantly more active than negatives in all cell lines (**Supplemental Figure S7**). This overlap likely represents the permissiveness of MPRAs, which are conducted outside the chromatin environment of the native locus. Importantly, our conclusion that most HARs show small quantitative differential activity between human and chimpanzee sequences is robust to the chosen threshold for active HARs.

### Modeling lentiMPRA *cis* and *trans* effects

To identify HARs with different enhancer activity between human and chimpanzee sequences (“*cis* effects”), we used the R limma (Ritchie et al., 2015) package (3.50.1) to fit a linear model for the mean log2(human [RNA/DNA] / chimpanzee [RNA/DNA]) of each HAR across 18 samples passing QC (human and chimpanzee cells, N2 and N3 stages) with code: lmFit(log2(human_chimp_ratios), model.matrix(∼ prep_date)) %>% eBayes(). This fits a linear model for each HAR that adjusts for the library preparation date, which we used to test for mean log-ratios significantly different from zero. P-values were adjusted for multiple testing using the false discovery rate (FDR eBayes q-value <1%), producing 188 differentially active HARs. We also explored using limma with voom or other variance stabilizing transformations but found that these were not needed because the log2(human[RNA/DNA] / chimpanzee [RNA/DNA]) values do not have a strong mean-variance relationship.

### Luciferase assays

To generate pLS-mP-Luc vector (Addgene 106253), minimal promoter and Luciferase gene fragment was amplified using pGL4.23 (Promega) as a template and inserted into pLS-mP (Addgene 81225) replacing with mP-EGFP. To generate pLS-SV40-mP-Rluc (Addgene106292), renilla luciferase gene was amplified using pGL4.74 (promega) as a template and inserted into pLS-SV40-mP vector (17) replacing with *EGFP* gene. We used an Agilent array to synthesize human and chimpanzee sequences of 2xHAR.11, 2xHAR.35, 2xHAR.53, 2xHAR.176, 2xHAR.273, 2xHAR.364, 2xHAR.401, 2xHAR.417, 2xHAR.434, and 2xHAR.518, and six negative control sequences (hg19 coordinates): N0 = chr1:10755200-10755371 (astrocyte progenitor H3K27me3 peak), N06 = chr7:27118200-27118371 (N2 H3K27me3 peak), N10 = chr4:8852800-8852971 (N1 H3K27me3 peak), N12 = chr17:46740400-46740571 (N2 H3K27me3 peak), N15 = chr19:1744200-1744371 (N1 H3K27me3 peak), N17 = chr14:37219958-37220129 (ENCODE negative control). These were synthesized along with homology arms on both sides (left: AGCCTGCATTTCTGCCAGGGCCCGCTCTAG, right: CTAGACCTGCAGGCACTAGAGGGTATATA), amplified using Agilent-luc.F and Agilent-luc.R primers (**Supplemental Table S9**), and cloned into XbaI site of the pLS-mP-luc using NEBuilder HiFi DNA Assembly Cloning Kit (NEB). Fragments that failed to clone (human 2xHAR.11, chimpanzee 2xHAR.35, human 2xHAR.176, human 2xHAR.273, chimpanzee 2xHAR.364, human and chimpanzee 2xHAR.434, and chimpanzee 2xHAR.518) were synthesized by Twist Bioscience along with homology arms: (left: TGTATATCCGGTCTCTTCTCTGGGTAGTCTCACTCAGCCTGCATTTCTGCCAGGGCCCGCTC TAG, right: CTAGACCTGCAGGCACTAGAGGGTATATAATGGAAGCTCGACTTCCAGCTTGGCAATCCGG TAC), amplified using Twist-luc.F and Twist-luc.R primers (**Supplemental Table S9**) and cloned into the pLS-mP-luc. Lentivirus was generated using standard methods (Wang and McManus, 2009), as described below for the library, individually for each clone with pLS-SV40-mP-Rluc spiked in at 10% of the total amount of plasmid used. 2x10^4^ cells per well (HS1 and Pt2 N3 cells) were seeded in a 96-well plate and were infected with virus 24 hours later. Three independent replicate cultures were transfected per plasmid and two biological replicates were done in different days. Firefly and Renilla luciferase activities were measured on a Synergy 2 microplate reader (BioTek) using the Dual-Luciferase Reporter Assay System (Promega). Enhancer activity was calculated as the fold change of each construct’s firefly luciferase activity normalized to renilla luciferase activity.

### Transgenic mouse reporter assays

We selected HAR152, 2xHAR.133, 2xHAR.518, and 2xHAR.548 for *in vivo* validation with mouse transient transgenic reporter assays based on their lentiMPRA activity, epigenetic profiles, and nearby genes. All HAR sequences were cloned into the Hsp68-LacZ vector (Addgene #37843) and validated by Sanger sequencing. LacZ transgenic mice were generated by Cyagen Biosciences using standard procedures(Pu et al., 2019), harvested and stained for LacZ expression as previously described(Pennacchio et al., 2006). Pictures were taken using an M165FC stereo microscope and a DFC500 12-megapixel camera (Leica).

### CRISPR activation experiment

2xHAR.183 was selected for further functional characterization due to its high predicted enhancer score (see Methods: Supervised and Unsupervised Learning Analysis) and overlaps with multiple chromatin interaction datasets. The HAR shares a TAD and has a significant chromatin loop with the ROCK2 gene in excitatory neuron PLAC-seq data (Song et al., 2020) and contacts ROCK2 in our N2/N3 Hi-C. The gene E2F6 is nearby on the linear genome but has fewer 3D chromatin contacts. Independent from our prediction, 2xHAR.183 overlaps a predicted FANTOM5 enhancer, an ENCODE candidate *cis*-regulatory element, and a ChromHMM enhancer annotated using fetal brain datasets.

Human excitatory neurons were generated using hiPSCs in the WTC11 background containing a doxycycline inducible neurogenin-2 at the AAVS1 safe harbour locus. In their undifferentiated state, cells were plated in Matrigel-coated plates and cultured with mTeSR media. mTeSR media was changed daily. To induce differentiation, cells were dissociated using Accutase and plated in Matrigel-coated plates. Cells were cultured for 3 days in pre-differentiation media containing KnockOut DMEM/F-12 with 2 ug/mL doxycycline supplemented with 1X N-2 Supplement, 1X NEAA, 10 ng/mL brain-derived neurothrophic factor (BDNF), 10 ng/mL NT-3, and 1ug/mL lamininin. On the first day, ROCK inhibitor was added to the predifferentiation media at a concentration of 10uM. Pre-differentiation media was changed daily for 3 days. To induce maturation, precursor cells were dissociated with Accutase and subplated in poly-D-Lysine coated plates. Cells were cultured in maturation media containing Neurobasal A and DMEM/F12 with 2 ug/mL doxycycline supplemented with 1X N-2 Supplement, 0.5 X B-27 Supplement, 1X NEAA, 0.5X GlutaMax, 10 ng/mL BDNF, 10 ng/mL NT-3, and 1ug/mL lamininin. Cells were maintained in the maturation media for the remaining 14 days. Half media changes were conducted on day 7 and day 14 of differentiation with maturation media minus doxycyline. After 14 days, wells were infected with lentivirus containing dCAS9-VP64_Blast (Addgene Plasmid #61425) and sgRNA targeting 2xHAR.183. The sgRNA sequence ATCATAGGATCAACTCGTTA was selected using CHOPCHOP to target 2xHAR.183 and was cloned into the pLG1 expression vector. Experiments were performed in triplicate and compared to wells infected with dCAS9-VP64 and no sgRNA. RNA was isolated after 5 days infection with lentivirus using the QIAGEN RNeasy kit with gDNA elimination column. RNA quality was investigated using the Agilent 2100 Bioanalyzer system and the RNA 6000 Nano kit, and an RNA integrity number over 9.0 was verified for all samples. cDNA was made from 1 microgram total RNA using SuperScript III First-Strand Synthesis SuperMix. qRT-PCR was performed using Maxima SYBR Green / ROX qPCR master mix and the following oligonucleotides:

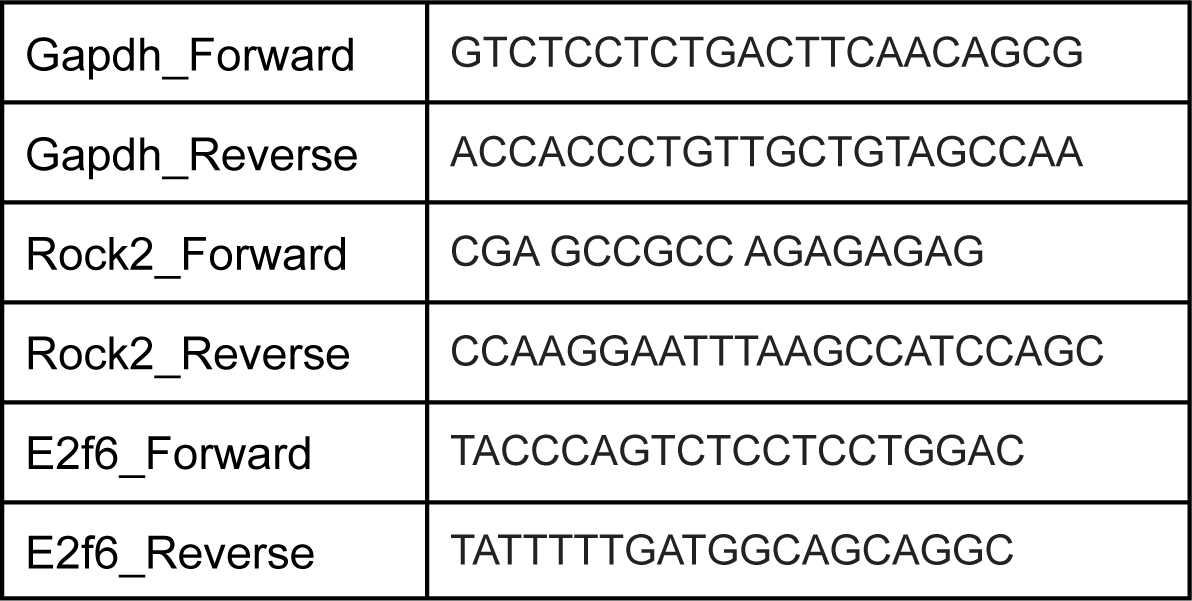

### Gene Ontology analysis

Gene Ontology (GO) terms associated with the 293 active HAR enhancers were separately compared to those of two background sets: all human N2 ATAC-seq peaks, and a random subset of 20k conserved elements identified with phastCons. Enrichment was computed with g:Profiler(Raudvere et al., 2019), using the custom statistical domain scope to provide the appropriate background set. The significance threshold was computed using g:SCS, which accounts for the non-independence of terms in the GO hierarchy.

### Predicting differential activity from footprints

HARs were intersected with TF footprints detected in human and chimpanzee N2 H3K27ac data (HINT (Gusmao et al., 2014) v0.13.2), excluding TFs not expressed in NPCs (TPM < 1, Kallisto (Bray et al., 2016) v0.48). Each HAR was then described by a vector of 762 binary features (presence/absence of 381 human and 381 chimpanzee footprints). A supervised gradient boosting regressor (XGBoost) was trained using these features and the log-scale RNA/DNA ratio from the MPRA as a continuous label. 80 percent of HARs were used for training and 20 percent used to detect overfitting during training (early stopping validation set). Variable importance was computed for each feature (TF x species). In-sample R2 and MSE were reported, as HARs are described by sparse and diverse sets of footprints that result in low R2 for out-of-sample data.

### Variant-disrupted footprints

The predicted chromatin state changes for all human:chimpanzee HAR variants were computed using Sei and overlapped with predicted footprints for NPC-expressed TFs (TPM > 1). For each TF, the number of overlapping variants and the maximum and minimum state change was computed. For visualization, TFs in the upper quartile of # variant overlaps were labeled if their maximum or minimum state change were in the top or bottom 18th percentile, respectively.

### Supervised and unsupervised learning analysis with *in vivo* epigenetic profiles

A large collection of open chromatin and TF binding datasets from multiple primary tissues (49% brain, 48% heart, 2% limb) were intersected with HARs and shown using Upset plots. The marks H3K4me1, H3K4me3, H3K9ac, H3K27ac, and H3K36me3 were labeled as activating. GEO and ENCODE accessions for these datasets are given in **Supplemental Table S3**. Feature vectors encoding the intersection of these datasets with HARs and validated VISTA enhancers(Visel et al., 2007) (all tissues) were projected into two dimensions using UMAP(McInnes et al., 2018) (umap-learn v0.5.3, n_neighbors = 8, metric = russellrao). For supervised learning, a logistic regression model with L1 (LASSO) penalty (scikit-learn (Pedregosa et al., 2011) 1.0.2) was trained using the feature vectors of validated VISTA enhancers and labeling brain enhancers as positives and non-brain enhancers (candidates that failed to validate or were active in other tissues) as negatives. This model then scored the similarity of HARs to neurodevelopmental enhancers.

### Deep learning characterization of HAR variants

All variants between human and chimpanzee HAR alleles were computed by first extracting alignments for each HAR (‘mafsInRegion‘) from the Zoonomia Consortium (Zoonomia Consortium, 2020), the largest multi-species alignment (MSA) to date. Human and chimpanzee alignments were then converted from MAF format to FASTA (‘msa_view‘), and from FASTA to VCF (‘jvarkit msa2vcf‘) (Lindenbaum). The deep learning tool Sei (Chen et al., 2022) then predicted changes in chromatin state for each human:chimpanzee variant across all HARs. Sei utilizes tens of thousands of human datasets which do not exist for chimpanzee; therefore, the predicted score estimates the impact of a chimpanzee variant relative to the human allele. To interpret these changes from an evolutionary perspective, we multiplied predicted scores by -1 so that chimpanzee variants with large negative scores would instead be positive (i.e., the human allele caused an increase) and large positive scores would instead be negative (i.e., the human allele caused a decrease).

### Permutation lentiMPRA

For each of three selected HARs with significant *cis* effects in our lentiMPRAs and prior evidence of enhancer activity (2xHAR.164, 2xHAR.170, 2xHAR.238), we designed oligos carrying all possible evolutionary intermediates (“permutations”) between homologous human and chimpanzee sequences. Some human HAR sequences differ in length from their homologous chimpanzee sequence due to short insertions and deletions. So to truly isolate the effects of individual human mutations in HARs, permutation oligos were created by mutating these sites and combinations thereof in the chimpanzee sequence. Thus, for HARs with insertions or deletions the oligo containing all human alleles is not the exact human genome sequence, but rather the chimpanzee sequence with all these mutations introduced. Permutation oligos were assayed, quantified and normalized alongside the main lentiMPRA library described above. The one-hot encoded oligo sequence, along with cell species and stage, were used to model the log RNA/DNA ratios for a given HAR using gradient boosting (XGBoost). The model estimates the importance of each nucleotide with a human-chimpanzee sequence difference, interactions between these nucleotides, and interactions between nucleotides and cell species or stage for predicting MPRA activity. We also assayed the permutation oligos as a separate library (“library 2”) in three technical replicates of two human (WTC, HS1-11) and two chimpanzee (Pt2A, Pt5C) cell lines, each at two stages of neural differentiation (N2, N3). For library 2, RNA and DNA count data was quantified as above, including normalizing for sequencing depth and batch correcting for library preparation date using limma.

### Analysis of HAR genetic and physical linkage to genes

Raw Hi-C data was aligned to hg38 and processed with juicer and distiller. Contact domains were called with the arrowhead algorithm in juicer (Durand et al., 2016), while chromatin loops were called using mustache (Roayaei Ardakany et al., 2020) on output from distiller (Goloborodko et al., 2019). LD blocks were computed using plink (Purcell et al., 2007) using 1000 Genomes (1000 Genomes Project Consortium et al., 2015) super-populations. HARs were annotated with multiple data sources including neurodevelopmental enhancer score (machine learning), closest protein coding gene (bedtools), neuropsychiatric variants in the same LD block for all (plink, bedtools), protein coding gene promoters sharing a contact domain in NPC/GPC or CP/GZ (Won et al., 2016) Hi-C data (juicer, bedtools), variants or protein coding gene promoters interacting with the HAR via chromatin looping in NPC/GPC or CP/GZ (Won et al., 2016) Hi-C data (mustache, bedtools), and genes interacting with the HAR via chromatin looping in primary tissue PCHi-C (Song et al., 2019) or PLAC-seq (Song et al., 2020) data (bedtools).

Variant datasets from the Psychiatric Genomics Consortium included ADHD (Demontis et al., 2019), Alzheimer’s Disease (Jansen et al., 2019), Autism Spectrum Disorder (Autism Spectrum Disorders Working Group of The Psychiatric Genomics Consortium, 2017), Bipolar Disorder (Mullins et al., 2021), Cross-Disorder (Cross-Disorder Group of the Psychiatric Genomics Consortium, 2019), Major Depressive Disorder (Wray et al., 2018), Obsessive-Compulsive Disorder (International Obsessive Compulsive Disorder Foundation Genetics Collaborative (IOCDF-GC) and OCD Collaborative Genetics Association Studies (OCGAS), 2018), Schizophrenia (in review), and Tourette’s Syndrome (Yu et al., 2019). Variant datasets from the PsychENCODE Consortium include expression QTLs (FDR < 0.05, > 1 FPKM in >= 20% of samples) and chromatin QTLs. Other variant datasets included chromatin QTLs in neurons and neural progenitors (Liang et al., 2021), expression QTLs in prefrontal cortex (Werling et al., 2020), and GTEx v8 fine-mapped brain expression QTLs (GTEx Consortium, 2013). Datasets using hg19 coordinates were mapped to hg38 using their rsid in combination with dbSNP build 155.

### Data and materials availability

All sequencing data is available through the Gene Expression Omnibus (GEO), accession number GSE110760 (chimpanzee cells: GSE110759, human cells: GSE110758). The unprocessed data is also available in the short read archive (SRA), BioProject accession numbers PRJNA428580 (chimpanzee cells) and PRJNA428579 (human cells).

## ACKNOWLEDGEMENTS

This work was funded by the Gladstone Institutes (K.S.P.), Schmidt Futures Foundation (A.A.P), and NIH grants: DP2MH122400-01 (A.A.P.), R35NS097305 (A.R.K.), FHG011569A (T.F.), R01MH109907 (N.A. and K.S.P.), U01MH116438 (N.A. and K.S.P.), UM1HG009408 (J.S. and N.A.), UM1HG011966 (J.S. and N.A.), and 2R01NS099099 (J.L.R.). K.S.P. is a Chan Zuckerberg Investigator. A.A.P is a New York Stem Cell Foundation Robertson Investigator and member of the UCSF Kavli Institute for Fundamental Neuroscience.

## AUTHOR CONTRIBUTIONS

S.W., H.R., F.I., N.A., and K.S.P. conceived and designed the study. S.W., A.W., S.T., and K.S.P. designed the lentiMPRA library. S.W., K.K., M.K., M.A.H.S., and K.S.P. analyzed data and performed modeling. F.I., H.R., T.F., A.A.P., E.M.P., B.M., B.A., O.E., D.L.C., and R.K. performed experiments. S.W., F.I., H.R., E.M.P., J.S., A.A.P., N.A., and K.S.P. interpreted results. E.U., J.L.R., A.R.K., J.S., A.A.P., N.A., and K.S.P. supervised and provided funding. S.W. and K.S.P. drafted the manuscript. All authors reviewed and edited the manuscript.

## References

1000 Genomes Project Consortium, Auton, A., Brooks, L.D., Durbin, R.M., Garrison, E.P., Kang, H.M., Korbel, J.O., Marchini, J.L., McCarthy, S., McVean, G.A., et al. (2015). A global reference for human genetic variation. Nature 526, 68–74.

Aldea, D., Atsuta, Y., Kokalari, B., Schaffner, S.F., Prasasya, R.D., Aharoni, A., Dingwall, H.L., Warder, B., and Kamberov, Y.G. (2021). Repeated mutation of a developmental enhancer contributed to human thermoregulatory evolution. Proc. Natl. Acad. Sci. U. S. A. 118. https://doi.org/10.1073/pnas.2021722118.

Autism Spectrum Disorders Working Group of The Psychiatric Genomics Consortium (2017). Meta-analysis of GWAS of over 16,000 individuals with autism spectrum disorder highlights a novel locus at 10q24.32 and a significant overlap with schizophrenia. Mol. Autism 8, 21.

Avsec, Ž., Agarwal, V., Visentin, D., Ledsam, J.R., Grabska-Barwinska, A., Taylor, K.R., Assael, Y., Jumper, J., Kohli, P., and Kelley, D.R. (2021). Effective gene expression prediction from sequence by integrating long-range interactions. Nat. Methods 18, 1196–1203.

Babbitt, C.C., Warner, L.R., Fedrigo, O., Wall, C.E., and Wray, G.A. (2011). Genomic signatures of diet-related shifts during human origins. Proceedings of the Royal Society B: Biological Sciences 278, 961–969. https://doi.org/10.1098/rspb.2010.2433.

Bae, B.-I., Jayaraman, D., and Walsh, C.A. (2015). Genetic changes shaping the human brain. Dev. Cell 32, 423–434.

Bershteyn, M., Nowakowski, T.J., Pollen, A.A., Di Lullo, E., Nene, A., Wynshaw-Boris, A., and Kriegstein, A.R. (2017). Human iPSC-Derived Cerebral Organoids Model Cellular Features of Lissencephaly and Reveal Prolonged Mitosis of Outer Radial Glia. Cell Stem Cell 20, 435–449.e4.

Boyd, J.L., Skove, S.L., Rouanet, J.P., Pilaz, L.-J., Bepler, T., Gordân, R., Wray, G.A., and Silver, D.L. (2015). Human-chimpanzee differences in a FZD8 enhancer alter cell-cycle dynamics in the developing neocortex. Curr. Biol. 25, 772–779.

Boyle, A.P., Hong, E.L., Hariharan, M., Cheng, Y., Schaub, M.A., Kasowski, M., Karczewski, K.J., Park, J., Hitz, B.C., Weng, S., et al. (2012). Annotation of functional variation in personal genomes using RegulomeDB. Genome Res. 22, 1790–1797.

Bray, N.L., Pimentel, H., Melsted, P., and Pachter, L. (2016). Near-optimal probabilistic RNA-seq quantification. Nat. Biotechnol. 34, 525–527.

Buenrostro, J.D., Giresi, P.G., Zaba, L.C., Chang, H.Y., and Greenleaf, W.J. (2013). Transposition of native chromatin for fast and sensitive epigenomic profiling of open chromatin, DNA-binding proteins and nucleosome position. Nat. Methods 10, 1213–1218.

Burns, J.K. (2004). An evolutionary theory of schizophrenia: cortical connectivity, metarepresentation, and the social brain. Behav. Brain Sci. 27, 831–855; discussion 855–885.

Capra, J.A., Erwin, G.D., McKinsey, G., Rubenstein, J.L.R., and Pollard, K.S. (2013). Many human accelerated regions are developmental enhancers. Philosophical Transactions of the Royal Society B: Biological Sciences 368, 20130025. https://doi.org/10.1098/rstb.2013.0025.

Castelijns, B., Baak, M.L., Timpanaro, I.S., Wiggers, C.R.M., Vermunt, M.W., Shang, P., Kondova, I., Geeven, G., Bianchi, V., de Laat, W., et al. (2020). Hominin-specific regulatory elements selectively emerged in oligodendrocytes and are disrupted in autism patients. Nat. Commun. 11, 301.

Chen, K.M., Wong, A.K., Troyanskaya, O.G., and Zhou, J. (2022). A sequence-based global map of regulatory activity for deciphering human genetics. Nat. Genet. 54, 940–949.

Cross-Disorder Group of the Psychiatric Genomics Consortium (2019). Genomic Relationships, Novel Loci, and Pleiotropic Mechanisms across Eight Psychiatric Disorders. Cell 179, 1469–1482.e11.

Crow, T.J. (1997). Is schizophrenia the price that Homo sapiens pays for language? Schizophrenia Research 28, 127–141. https://doi.org/10.1016/s0920-9964(97)00110-2.

Crow, T.J. (2000). Schizophrenia as the price that homo sapiens pays for language: a resolution of the central paradox in the origin of the species. Brain Res. Brain Res. Rev. 31, 118–129.

Demontis, D., Walters, R.K., Martin, J., Mattheisen, M., Als, T.D., Agerbo, E., Baldursson, G., Belliveau, R., Bybjerg-Grauholm, J., Bækvad-Hansen, M., et al. (2019). Discovery of the first genome-wide significant risk loci for attention deficit/hyperactivity disorder. Nat. Genet. 51, 63–75.

Divya, T.S., Lalitha, S., Parvathy, S., Subashini, C., Sanalkumar, R., Dhanesh, S.B., Rasheed, V.A., Divya, M.S., Tole, S., and James, J. (2016). Regulation of Tlx3 by Pax6 is required for the restricted expression of Chrnα3 in Cerebellar Granule Neuron progenitors during development. Sci. Rep. 6, 30337.

Doan, R.N., Bae, B.-I., Cubelos, B., Chang, C., Hossain, A.A., Al-Saad, S., Mukaddes, N.M., Oner, O., Al-Saffar, M., Balkhy, S., et al. (2016). Mutations in Human Accelerated Regions Disrupt Cognition and Social Behavior. Cell 167, 341–354.e12.

Durand, N.C., Shamim, M.S., Machol, I., Rao, S.S.P., Huntley, M.H., Lander, E.S., and Aiden, E.L. (2016). Juicer Provides a One-Click System for Analyzing Loop-Resolution Hi-C Experiments. Cell Systems 3, 95–98. https://doi.org/10.1016/j.cels.2016.07.002.

Dutrow, E.V., Emera, D., Yim, K., Uebbing, S., Kocher, A.A., Krenzer, M., Nottoli, T., Burkhardt, D.B., Krishnaswamy, S., Louvi, A., et al. (2022). Modeling uniquely human gene regulatory function via targeted humanization of the mouse genome. Nat. Commun. 13, 304.

Ernst, J., and Kellis, M. (2017). Chromatin-state discovery and genome annotation with ChromHMM. Nat. Protoc. 12, 2478–2492.

Erwin, G.D., Oksenberg, N., Truty, R.M., Kostka, D., Murphy, K.K., Ahituv, N., Pollard, K.S., and Capra, J.A. (2014). Integrating diverse datasets improves developmental enhancer prediction. PLoS Comput. Biol. 10, e1003677.

Franchini, L.F., and Pollard, K.S. (2017). Human evolution: the non-coding revolution. BMC Biol. 15, 89.

Funk, C.C., Casella, A.M., Jung, S., Richards, M.A., Rodriguez, A., Shannon, P., Donovan-Maiye, R., Heavner, B., Chard, K., Xiao, Y., et al. (2020). Atlas of Transcription Factor Binding Sites from ENCODE DNase Hypersensitivity Data across 27 Tissue Types. Cell Rep. 32, 108029.

Furlan, A., Lübke, M., Adameyko, I., Lallemend, F., and Ernfors, P. (2013). The transcription factor Hmx1 and growth factor receptor activities control sympathetic neurons diversification. EMBO J. 32, 1613–1625.

Girskis, K.M., Stergachis, A.B., DeGennaro, E.M., Doan, R.N., Qian, X., Johnson, M.B., Wang, P.P., Sejourne, G.M., Nagy, M.A., Pollina, E.A., et al. (2021). Rewiring of human neurodevelopmental gene regulatory programs by human accelerated regions. Neuron 109, 3239–3251.e7.

Goloborodko, A., Venev, S., Abdennur, N., azkalot, and Di Tommaso, P. (2019). mirnylab/distiller-nf: v0.3.3.

GTEx Consortium (2013). The Genotype-Tissue Expression (GTEx) project. Nat. Genet. 45, 580–585.

Gusmao, E.G., Dieterich, C., Zenke, M., and Costa, I.G. (2014). Detection of active transcription factor binding sites with the combination of DNase hypersensitivity and histone modifications. Bioinformatics 30, 3143–3151.

Hammal, F., de Langen, P., Bergon, A., Lopez, F., and Ballester, B. (2022). ReMap 2022: a database of Human, Mouse, Drosophila and Arabidopsis regulatory regions from an integrative analysis of DNA-binding sequencing experiments. Nucleic Acids Res. 50, D316–D325.

Hormozdiari, F., Zhu, A., Kichaev, G., Ju, C.J.-T., Segrè, A.V., Joo, J.W.J., Won, H., Sankararaman, S., Pasaniuc, B., Shifman, S., et al. (2017). Widespread Allelic Heterogeneity in Complex Traits. Am. J. Hum. Genet. 100, 789–802.

Hubisz, M.J., and Pollard, K.S. (2014). Exploring the genesis and functions of Human Accelerated Regions sheds light on their role in human evolution. Curr. Opin. Genet. Dev. 29, 15–21.

Inoue, F., and Ahituv, N. (2015). Decoding enhancers using massively parallel reporter assays. Genomics 106, 159–164.

Inoue, F., Kircher, M., Martin, B., Cooper, G.M., Witten, D.M., McManus, M.T., Ahituv, N., and Shendure, J. (2017). A systematic comparison reveals substantial differences in chromosomal versus episomal encoding of enhancer activity. Genome Research 27, 38–52. https://doi.org/10.1101/gr.212092.116.

Inoue, F., Kreimer, A., Ashuach, T., Ahituv, N., and Yosef, N. (2019). Identification and Massively Parallel Characterization of Regulatory Elements Driving Neural Induction. Cell Stem Cell 25, 713–727.e10.

International Obsessive Compulsive Disorder Foundation Genetics Collaborative (IOCDF-GC) and OCD Collaborative Genetics Association Studies (OCGAS) (2018). Revealing the complex genetic architecture of obsessive-compulsive disorder using meta-analysis. Mol. Psychiatry 23, 1181–1188.

Jagoda, E., Xue, J.R., Reilly, S.K., Dannemann, M., Racimo, F., Huerta-Sanchez, E., Sankararaman, S., Kelso, J., Pagani, L., Sabeti, P.C., et al. (2022). Detection of Neanderthal Adaptively Introgressed Genetic Variants That Modulate Reporter Gene Expression in Human Immune Cells. Mol. Biol. Evol. 39. https://doi.org/10.1093/molbev/msab304.

Jansen, I.E., Savage, J.E., Watanabe, K., Bryois, J., Williams, D.M., Steinberg, S., Sealock, J., Karlsson, I.K., Hägg, S., Athanasiu, L., et al. (2019). Genome-wide meta-analysis identifies new loci and functional pathways influencing Alzheimer’s disease risk. Nat. Genet. 51, 404–413.

Kamm, G.B., Pisciottano, F., Kliger, R., and Franchini, L.F. (2013). The developmental brain gene NPAS3 contains the largest number of accelerated regulatory sequences in the human genome. Mol. Biol. Evol. 30, 1088–1102.

Kanton, S., Boyle, M.J., He, Z., Santel, M., Weigert, A., Sanchís-Calleja, F., Guijarro, P., Sidow, L., Fleck, J.S., Han, D., et al. (2019). Organoid single-cell genomic atlas uncovers human-specific features of brain development. Nature 574, 418–422.

Kim, D., Langmead, B., and Salzberg, S.L. (2015). HISAT: a fast spliced aligner with low memory requirements. Nature Methods 12, 357–360. https://doi.org/10.1038/nmeth.3317.

Kircher, M. (2012). Analysis of High-Throughput Ancient DNA Sequencing Data. Methods in Molecular Biology 197–228. https://doi.org/10.1007/978-1-61779-516-9_23.

Kostka, D., Hubisz, M.J., Siepel, A., and Pollard, K.S. (2012). The role of GC-biased gene conversion in shaping the fastest evolving regions of the human genome. Mol. Biol. Evol. 29, 1047–1057.

Kwasnieski, J.C., Fiore, C., Chaudhari, H.G., and Cohen, B.A. (2014). High-throughput functional testing of ENCODE segmentation predictions. Genome Res. 24, 1595–1602.

Lein, E.S., Hawrylycz, M.J., Ao, N., Ayres, M., Bensinger, A., Bernard, A., Boe, A.F., Boguski, M.S., Brockway, K.S., Byrnes, E.J., et al. (2007). Genome-wide atlas of gene expression in the adult mouse brain. Nature 445, 168–176.

Liang, D., Elwell, A.L., Aygün, N., Krupa, O., Wolter, J.M., Kyere, F.A., Lafferty, M.J., Cheek, K.E., Courtney, K.P., Yusupova, M., et al. (2021). Cell-type-specific effects of genetic variation on chromatin accessibility during human neuronal differentiation. Nat. Neurosci. 24, 941–953.

Liao, Y., Smyth, G.K., and Shi, W. (2013). The Subread aligner: fast, accurate and scalable read mapping by seed-and-vote. Nucleic Acids Research 41, e108–e108. https://doi.org/10.1093/nar/gkt214.

Lindblad-Toh, K., Garber, M., Zuk, O., Lin, M.F., Parker, B.J., Washietl, S., Kheradpour, P., Ernst, J., Jordan, G., Mauceli, E., et al. (2011). A high-resolution map of human evolutionary constraint using 29 mammals. Nature 478, 476–482.

Lindenbaum, P. jvarkit: Java utilities for Bioinformatics (Github).

Lindhorst, D., and Halfon, M.S. (2022). Reporter gene assays and chromatin-level assays define substantially non-overlapping sets of enhancer sequences.

Lizio, M., Harshbarger, J., Shimoji, H., Severin, J., Kasukawa, T., Sahin, S., Abugessaisa, I., Fukuda, S., Hori, F., Ishikawa-Kato, S., et al. (2015). Gateways to the FANTOM5 promoter level mammalian expression atlas. Genome Biol. 16, 22.

Markenscoff-Papadimitriou, E., Whalen, S., Przytycki, P., Thomas, R., Binyameen, F., Nowakowski, T.J., Kriegstein, A.R., Sanders, S.J., State, M.W., Pollard, K.S., et al. (2020). A Chromatin Accessibility Atlas of the Developing Human Telencephalon. Cell 182, 754–769.e18.

Mattioli, K., Oliveros, W., Gerhardinger, C., Andergassen, D., Maass, P.G., Rinn, J.L., and Melé, M. (2020). Cis and trans effects differentially contribute to the evolution of promoters and enhancers. Genome Biol. 21, 210.

McInnes, L., Healy, J., Saul, N., and Großberger, L. (2018). UMAP: Uniform Manifold Approximation and Projection. J. Open Source Softw. 3, 861.

Miyaoka, Y., Chan, A.H., Judge, L.M., Yoo, J., Huang, M., Nguyen, T.D., Lizarraga, P.P., So, P.-L., and Conklin, B.R. (2014). Isolation of single-base genome-edited human iPS cells without antibiotic selection. Nature Methods 11, 291–293. https://doi.org/10.1038/nmeth.2840.

Mullins, N., Forstner, A.J., O’Connell, K.S., Coombes, B., Coleman, J.R.I., Qiao, Z., Als, T.D., Bigdeli, T.B., Børte, S., Bryois, J., et al. (2021). Genome-wide association study of more than 40,000 bipolar disorder cases provides new insights into the underlying biology. Nat. Genet. 53, 817–829.

Norman, A.R., Ryu, A.H., Jamieson, K., Thomas, S., Shen, Y., Ahituv, N., Pollard, K.S., and Reiter, J.F. (2021). A Human Accelerated Region is a Leydig cell GLI2 Enhancer that Affects Male-Typical Behavior.

Okita, K., Yamakawa, T., Matsumura, Y., Sato, Y., Amano, N., Watanabe, A., Goshima, N., and Yamanaka, S. (2013). An efficient nonviral method to generate integration-free human-induced pluripotent stem cells from cord blood and peripheral blood cells. Stem Cells 31, 458–466.

Pedregosa, F., Varoquaux, G., Gramfort, A., Michel, V., Thirion, B., Grisel, O., Blondel, M., Prettenhofer, P., Weiss, R., Dubourg, V., et al. (2011). Scikit-learn: Machine Learning in Python. Journal of Machine Learning Research 12, 2825–2830.

Pennacchio, L.A., Ahituv, N., Moses, A.M., Prabhakar, S., Nobrega, M.A., Shoukry, M., Minovitsky, S., Dubchak, I., Holt, A., Lewis, K.D., et al. (2006). In vivo enhancer analysis of human conserved non-coding sequences. Nature 444, 499–502.

Pollard, K.S., Salama, S.R., King, B., Kern, A.D., Dreszer, T., Katzman, S., Siepel, A., Pedersen, J.S., Bejerano, G., Baertsch, R., et al. (2006a). Forces shaping the fastest evolving regions in the human genome. PLoS Genet. 2, e168.

Pollard, K.S., Salama, S.R., Lambert, N., Lambot, M.-A., Coppens, S., Pedersen, J.S., Katzman, S., King, B., Onodera, C., Siepel, A., et al. (2006b). An RNA gene expressed during cortical development evolved rapidly in humans. Nature 443, 167–172.

Pollen, A.A., Bhaduri, A., Andrews, M.G., Nowakowski, T.J., Meyerson, O.S., Mostajo-Radji, M.A., Di Lullo, E., Alvarado, B., Bedolli, M., Dougherty, M.L., et al. (2019). Establishing Cerebral Organoids as Models of Human-Specific Brain Evolution. Cell 176, 743–756.e17.

Prabhakar, S., Noonan, J.P., Pääbo, S., and Rubin, E.M. (2006). Accelerated evolution of conserved noncoding sequences in humans. Science 314, 786.

Prabhakar, S., Visel, A., Akiyama, J.A., Shoukry, M., Lewis, K.D., Holt, A., Plajzer-Frick, I., Morrison, H., Fitzpatrick, D.R., Afzal, V., et al. (2008). Human-specific gain of function in a developmental enhancer. Science 321, 1346–1350.

Pu, X.-A., Young, A.P., and Kubisch, H.M. (2019). Production of Transgenic Mice by Pronuclear Microinjection. In Microinjection: Methods and Protocols, C. Liu, and Y. Du, eds. (New York, NY: Springer New York), pp. 17–41.

Purcell, S., Neale, B., Todd-Brown, K., Thomas, L., Ferreira, M.A.R., Bender, D., Maller, J., Sklar, P., de Bakker, P.I.W., Daly, M.J., et al. (2007). PLINK: a tool set for whole-genome association and population-based linkage analyses. Am. J. Hum. Genet. 81, 559–575.

Raudvere, U., Kolberg, L., Kuzmin, I., Arak, T., Adler, P., Peterson, H., and Vilo, J. (2019). g:Profiler: a web server for functional enrichment analysis and conversions of gene lists (2019 update). Nucleic Acids Res. 47, W191–W198.

Ritchie, M.E., Phipson, B., Wu, D., Hu, Y., Law, C.W., Shi, W., and Smyth, G.K. (2015). limma powers differential expression analyses for RNA-sequencing and microarray studies. Nucleic Acids Res. 43, e47.

Ritter, D.I., Li, Q., Kostka, D., Pollard, K.S., Guo, S., and Chuang, J.H. (2010). The importance of being cis: evolution of orthologous fish and mammalian enhancer activity. Mol. Biol. Evol. 27, 2322–2332.

Roayaei Ardakany, A., Gezer, H.T., Lonardi, S., and Ay, F. (2020). Mustache: multi-scale detection of chromatin loops from Hi-C and Micro-C maps using scale-space representation. Genome Biol. 21, 256.

Schwartz, M.P., Hou, Z., Propson, N.E., Zhang, J., Engstrom, C.J., Santos Costa, V., Jiang, P., Nguyen, B.K., Bolin, J.M., Daly, W., et al. (2015). Human pluripotent stem cell-derived neural constructs for predicting neural toxicity. Proc. Natl. Acad. Sci. U. S. A. 112, 12516–12521.

Song, M., Yang, X., Ren, X., Maliskova, L., Li, B., Jones, I.R., Wang, C., Jacob, F., Wu, K., Traglia, M., et al. (2019). Mapping cis-regulatory chromatin contacts in neural cells links neuropsychiatric disorder risk variants to target genes. Nat. Genet. 51, 1252–1262.

Song, M., Pebworth, M.-P., Yang, X., Abnousi, A., Fan, C., Wen, J., Rosen, J.D., Choudhary, M.N.K., Cui, X., Jones, I.R., et al. (2020). Cell-type-specific 3D epigenomes in the developing human cortex. Nature 587, 644–649.

Sullivan, P.F., Agrawal, A., Bulik, C.M., Andreassen, O.A., Børglum, A.D., Breen, G., Cichon, S., Edenberg, H.J., Faraone, S.V., Gelernter, J., et al. (2018). Psychiatric Genomics: An Update and an Agenda. Am. J. Psychiatry 175, 15–27.

Uebbing, S., Gockley, J., Reilly, S.K., Kocher, A.A., Geller, E., Gandotra, N., Scharfe, C., Cotney, J., and Noonan, J.P. (2021). Massively parallel discovery of human-specific substitutions that alter enhancer activity. Proc. Natl. Acad. Sci. U. S. A. 118. https://doi.org/10.1073/pnas.2007049118.

Vaishnav, E.D., de Boer, C.G., Molinet, J., Yassour, M., Fan, L., Adiconis, X., Thompson, D.A., Levin, J.Z., Cubillos, F.A., and Regev, A. (2022). The evolution, evolvability and engineering of gene regulatory DNA. Nature 603, 455–463.

Visel, A., Minovitsky, S., Dubchak, I., and Pennacchio, L.A. (2007). VISTA Enhancer Browser--a database of tissue-specific human enhancers. Nucleic Acids Res. 35, D88–D92.

Voisey, J., Mehta, D., McLeay, R., Morris, C.P., Wockner, L.F., Noble, E.P., Lawford, B.R., and Young, R.M. (2017). Clinically proven drug targets differentially expressed in the prefrontal cortex of schizophrenia patients. Brain Behav. Immun. 61, 259–265.

Wang, X., and McManus, M. (2009). Lentivirus Production. Journal of Visualized Experiments https://doi.org/10.3791/1499.

Wang, C., Ward, M.E., Chen, R., Liu, K., Tracy, T.E., Chen, X., Xie, M., Sohn, P.D., Ludwig, C., Meyer-Franke, A., et al. (2017). Scalable Production of iPSC-Derived Human Neurons to Identify Tau-Lowering Compounds by High-Content Screening. Stem Cell Reports 9, 1221–1233.

Wang, D., Liu, S., Warrell, J., Won, H., Shi, X., Navarro, F.C.P., Clarke, D., Gu, M., Emani, P., Yang, Y.T., et al. (2018). Comprehensive functional genomic resource and integrative model for the human brain. Science 362. https://doi.org/10.1126/science.aat8464.

Weiss, C.V., Harshman, L., Inoue, F., Fraser, H.B., Petrov, D.A., Ahituv, N., and Gokhman, D. (2021). The cis-regulatory effects of modern human-specific variants. Elife 10. https://doi.org/10.7554/eLife.63713.

Werling, D.M., Pochareddy, S., Choi, J., An, J.-Y., Sheppard, B., Peng, M., Li, Z., Dastmalchi, C., Santpere, G., Sousa, A.M.M., et al. (2020). Whole-Genome and RNA Sequencing Reveal Variation and Transcriptomic Coordination in the Developing Human Prefrontal Cortex. Cell Rep. 31, 107489.

Won, H., de la Torre-Ubieta, L., Stein, J.L., Parikshak, N.N., Huang, J., Opland, C.K., Gandal, M.J., Sutton, G.J., Hormozdiari, F., Lu, D., et al. (2016). Chromosome conformation elucidates regulatory relationships in developing human brain. Nature 538, 523–527.

Wray, N.R., Ripke, S., Mattheisen, M., Trzaskowski, M., Byrne, E.M., Abdellaoui, A., Adams, M.J., Agerbo, E., Air, T.M., Andlauer, T.M.F., et al. (2018). Genome-wide association analyses identify 44 risk variants and refine the genetic architecture of major depression. Nat. Genet. 50, 668–681.

Wu, Y., Li, X., Liu, J., Luo, X.-J., and Yao, Y.-G. (2020). SZDB2.0: an updated comprehensive resource for schizophrenia research. Hum. Genet. 139, 1285–1297.

Yu, D., Sul, J.H., Tsetsos, F., Nawaz, M.S., Huang, A.Y., Zelaya, I., Illmann, C., Osiecki, L., Darrow, S.M., Hirschtritt, M.E., et al. (2019). Interrogating the Genetic Determinants of Tourette’s Syndrome and Other Tic Disorders Through Genome-Wide Association Studies. AJP 176, 217–227.

Zhou, J., Theesfeld, C.L., Yao, K., Chen, K.M., Wong, A.K., and Troyanskaya, O.G. (2018). Deep learning sequence-based ab initio prediction of variant effects on expression and disease risk. Nat. Genet. 50, 1171–1179.

Zoonomia Consortium (2020). A comparative genomics multitool for scientific discovery and conservation. Nature 587, 240–245.

