## Supplementary figures and images for "Machine-learning dissection of Human Accelerated Regions in primate neurodevelopment"

### Figure S1

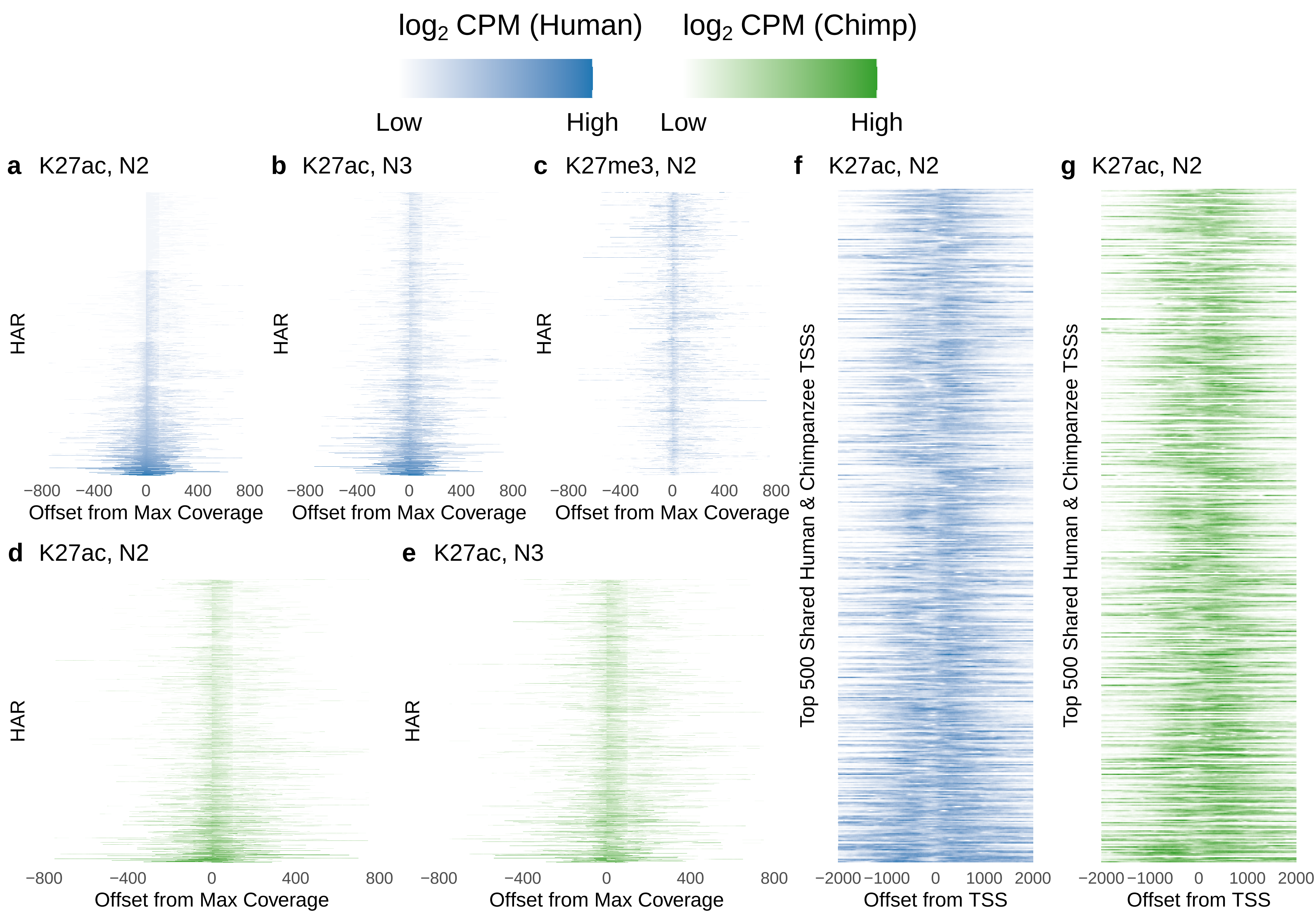

### Figure S2

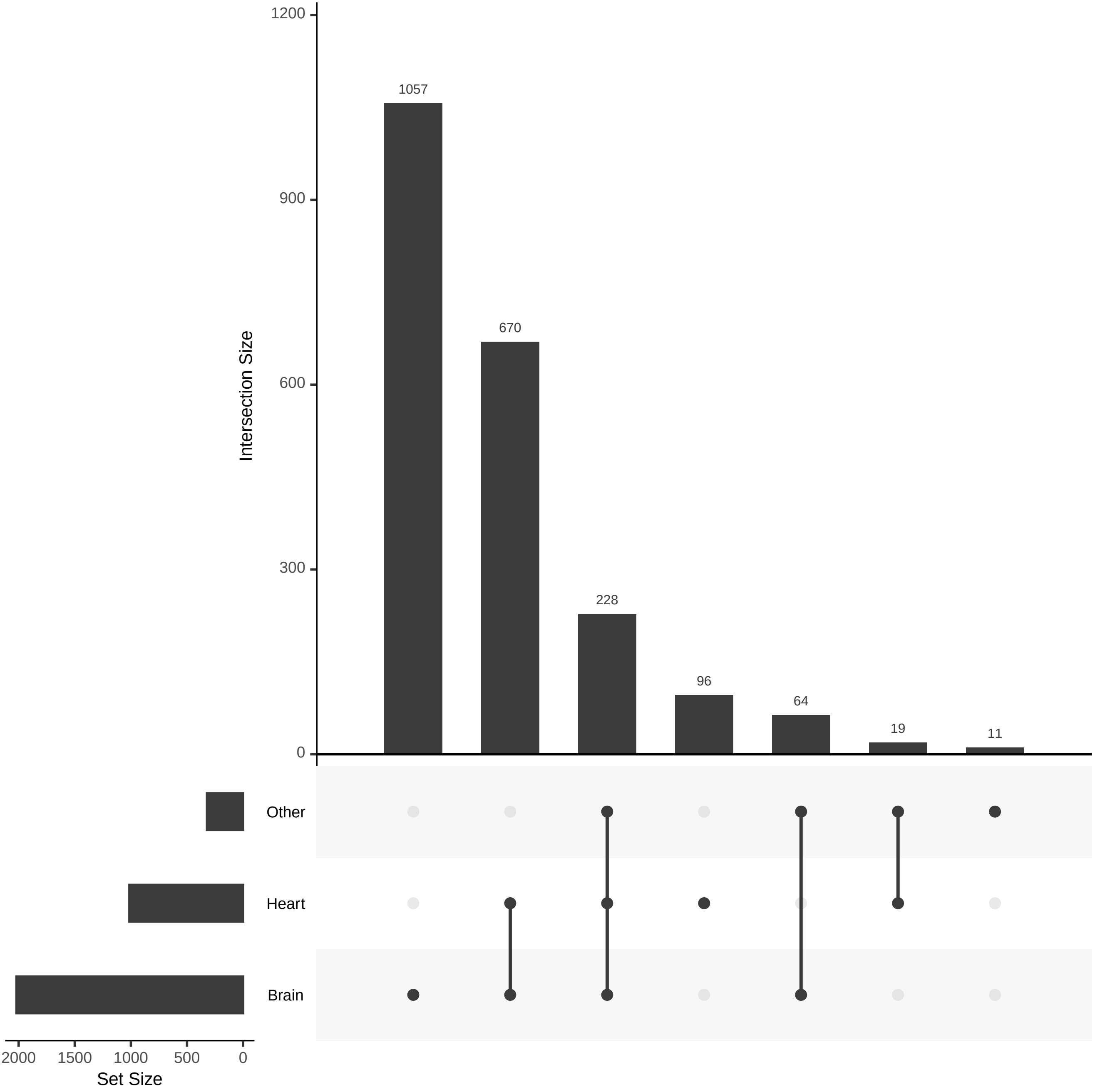

### Figure S3

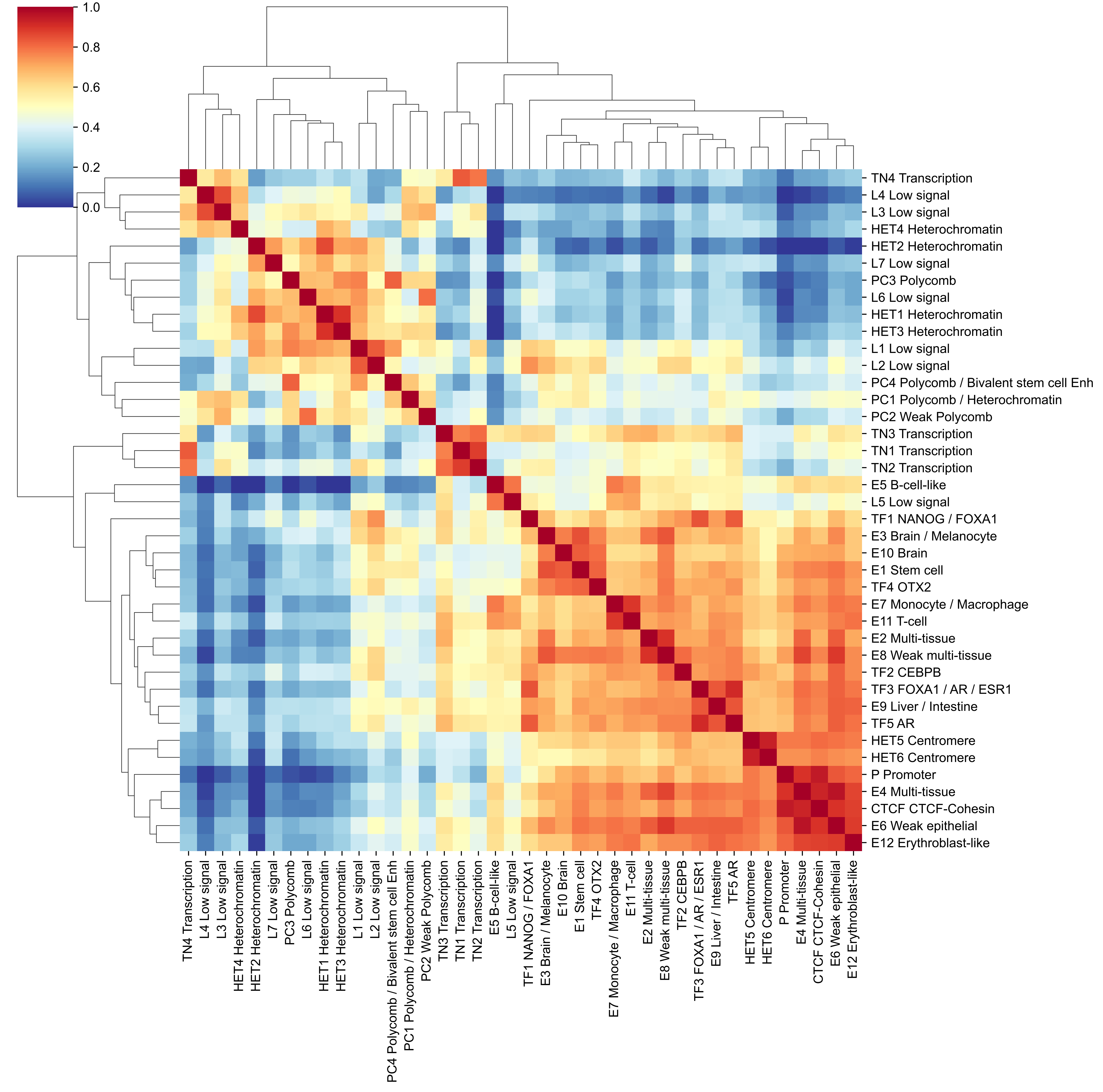

### Figure S4

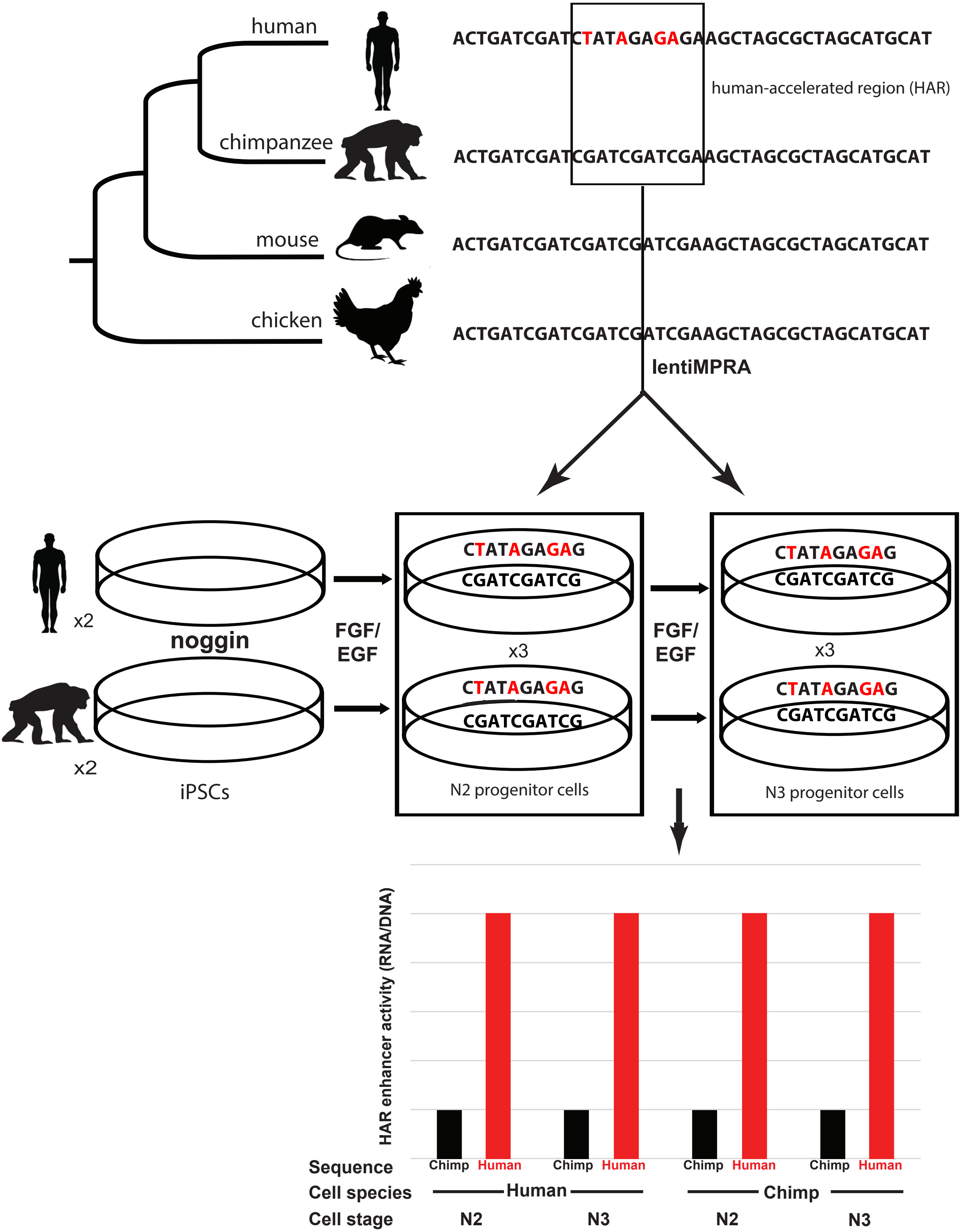

### Figure S5

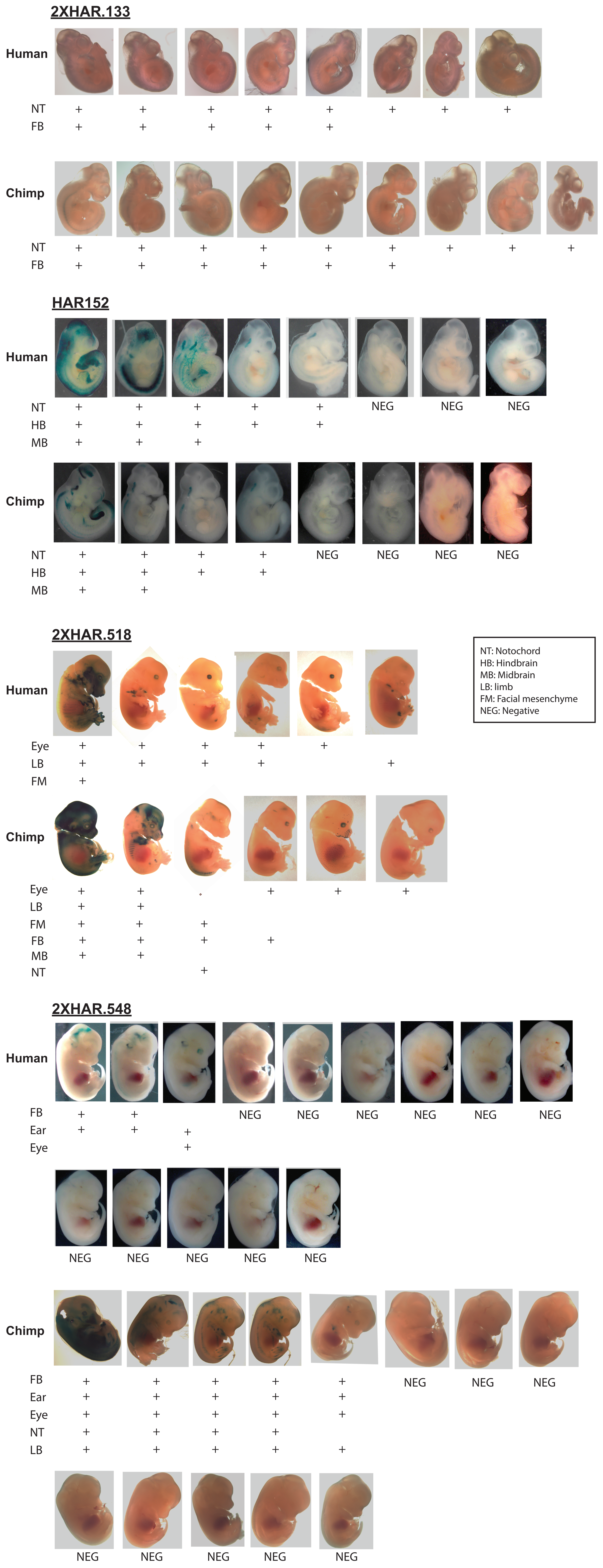

### Figure S6

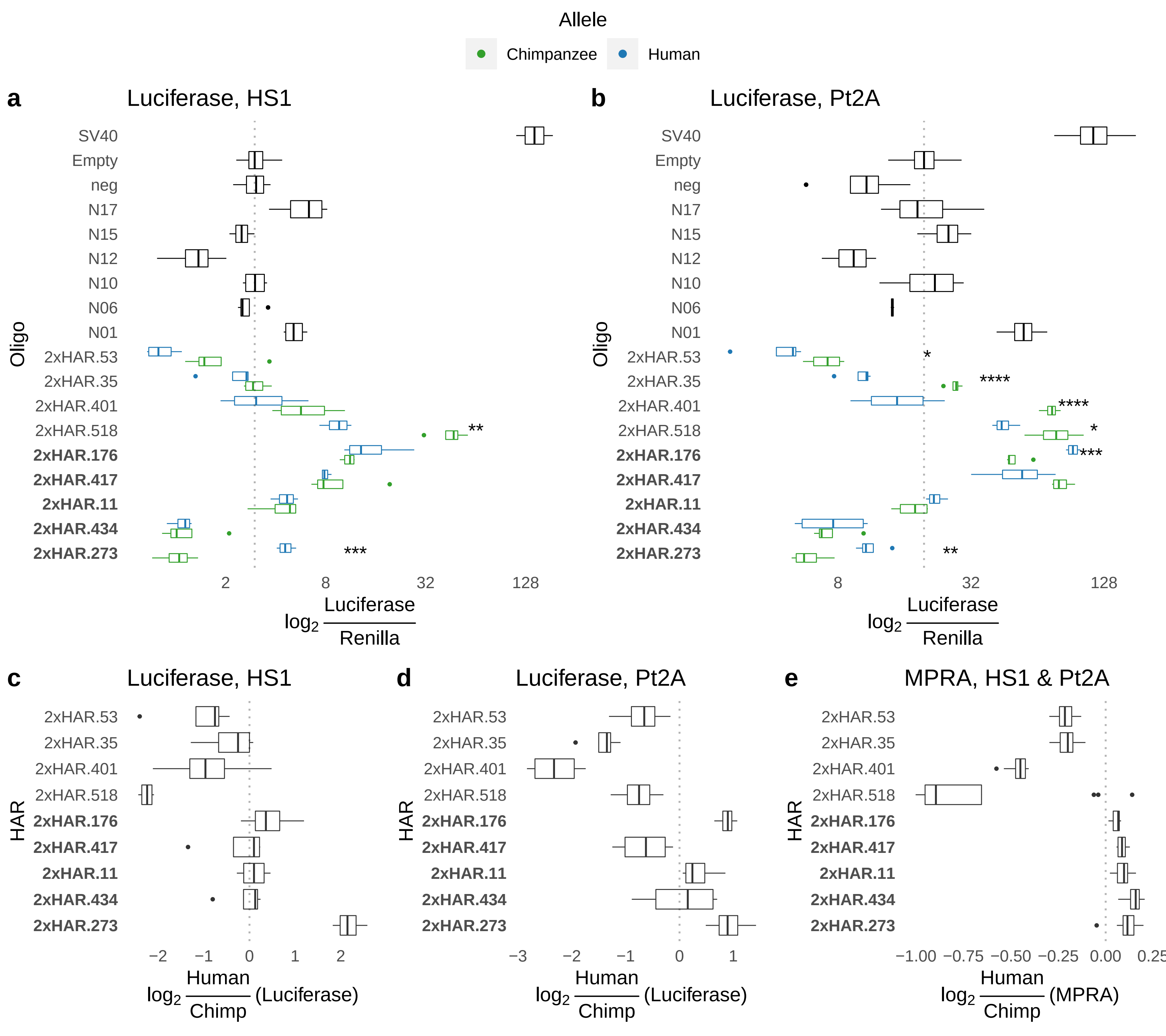

### Figure S7

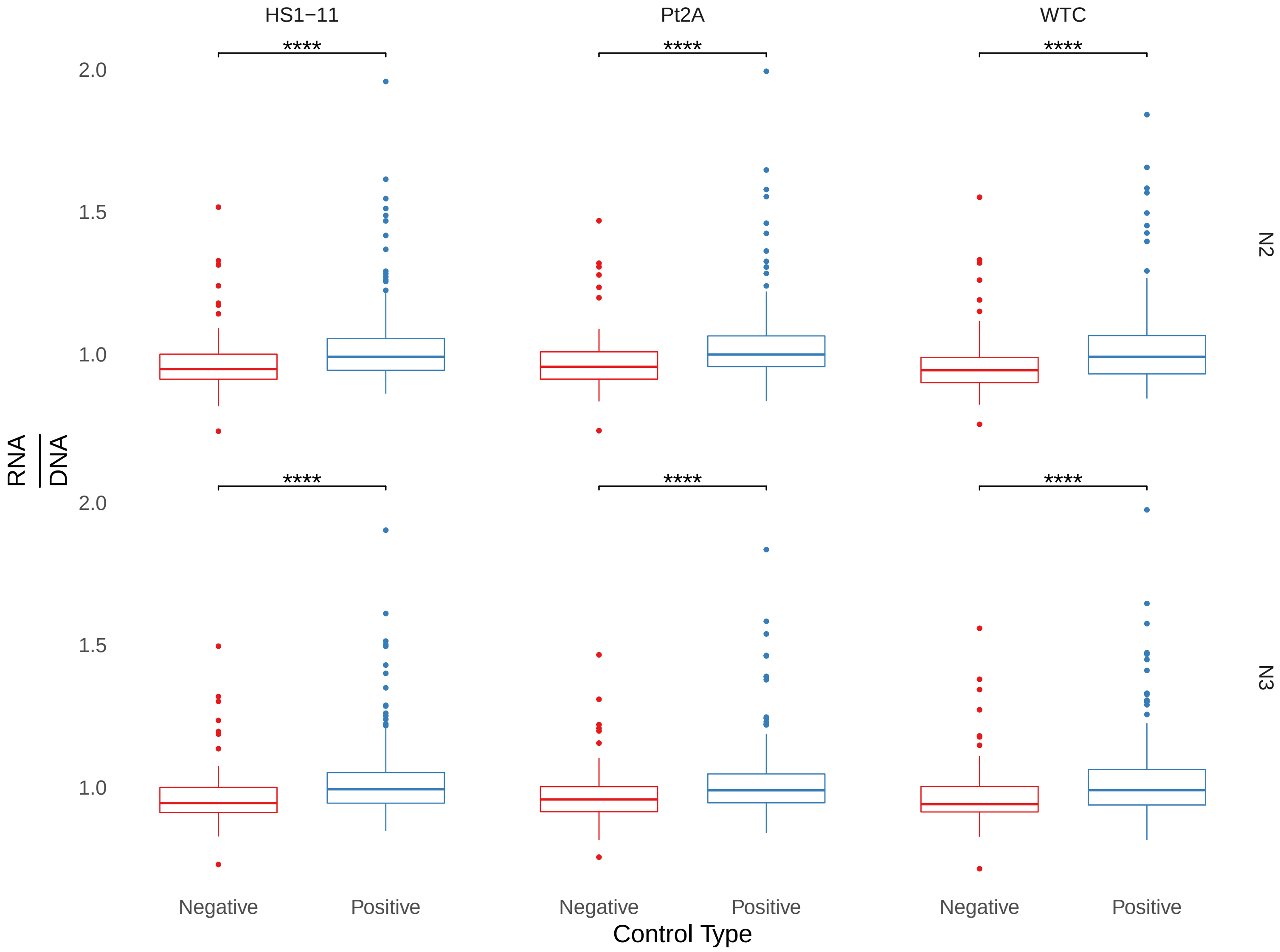

### Figure S8

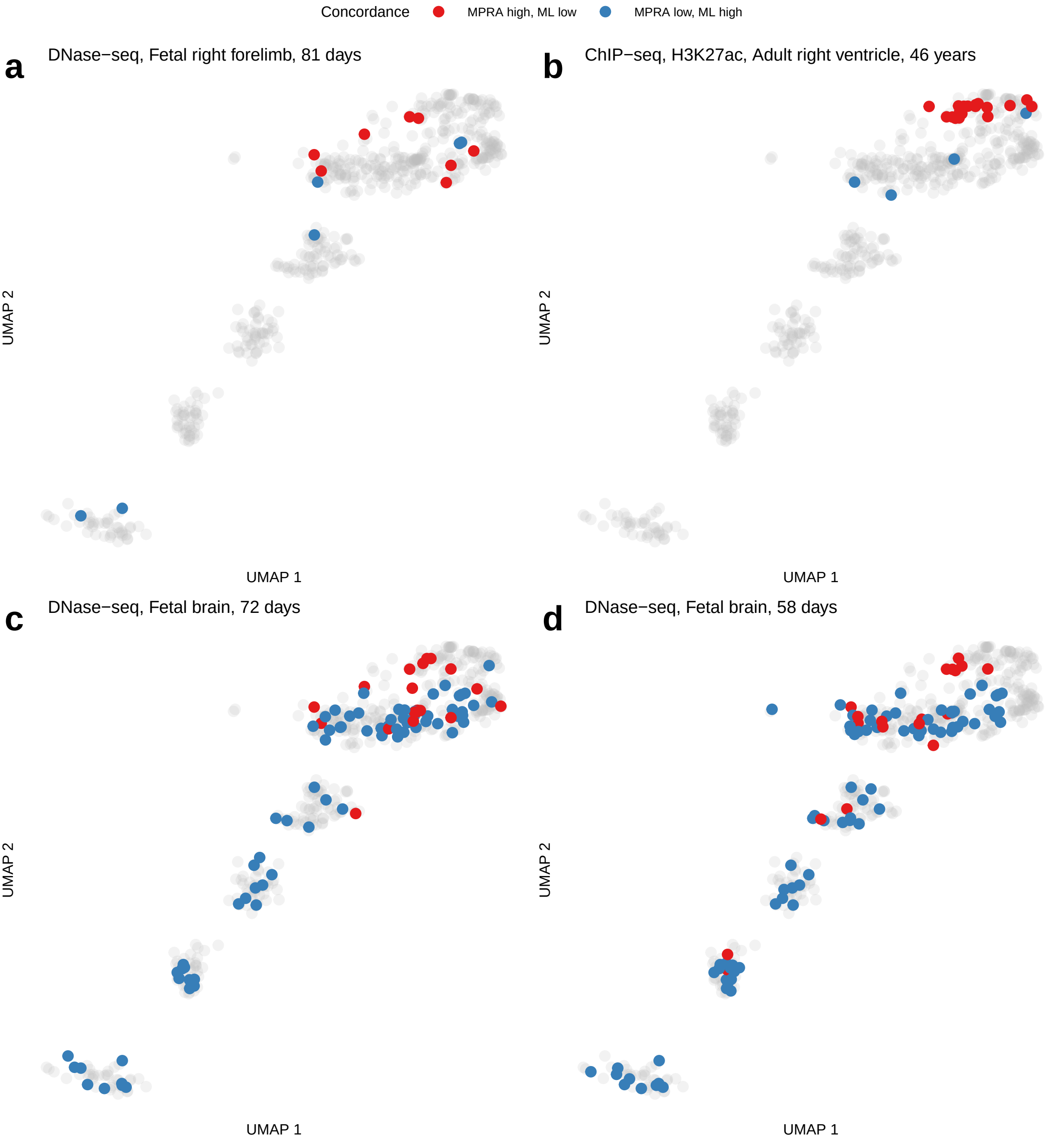

### Figure S9

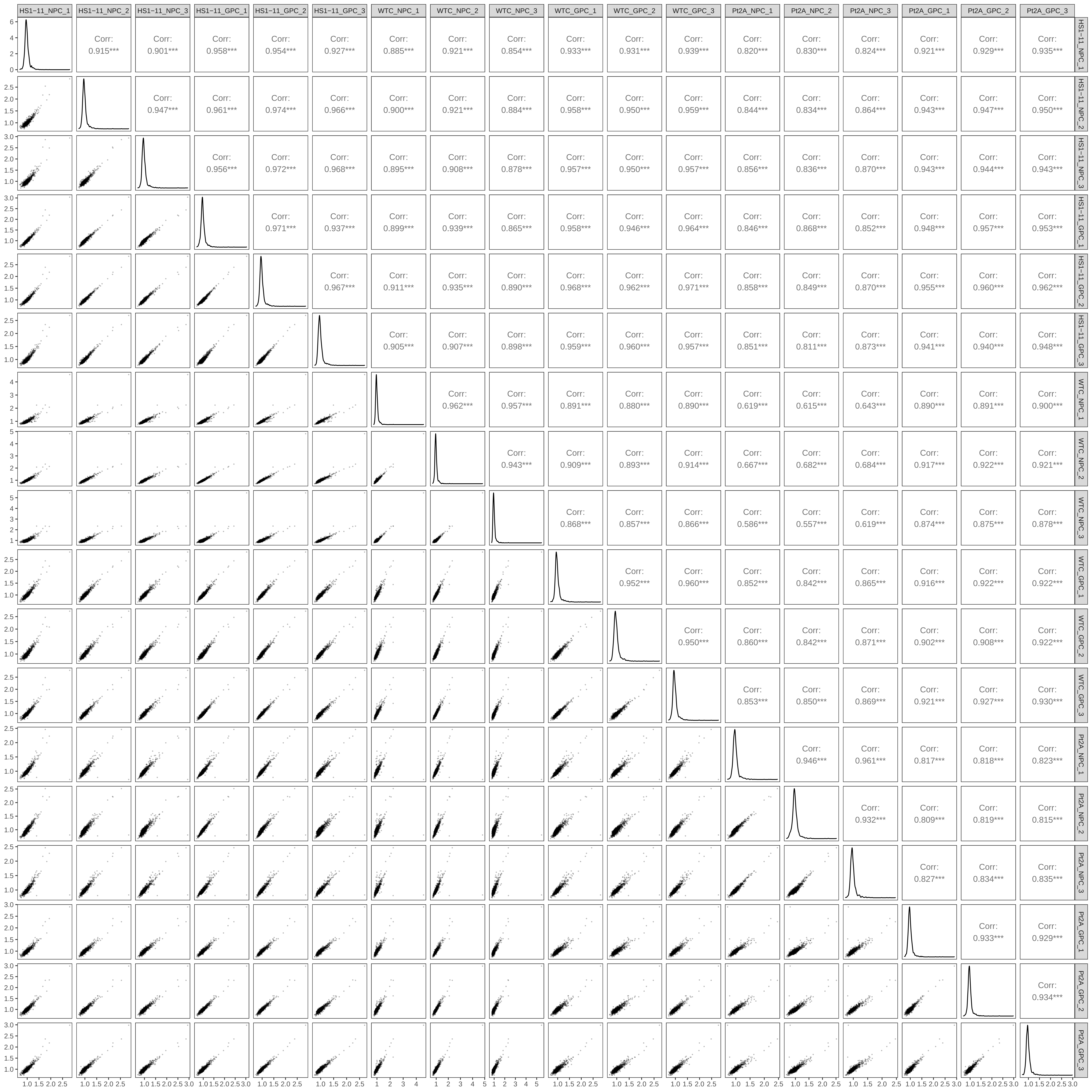

### Figure S10

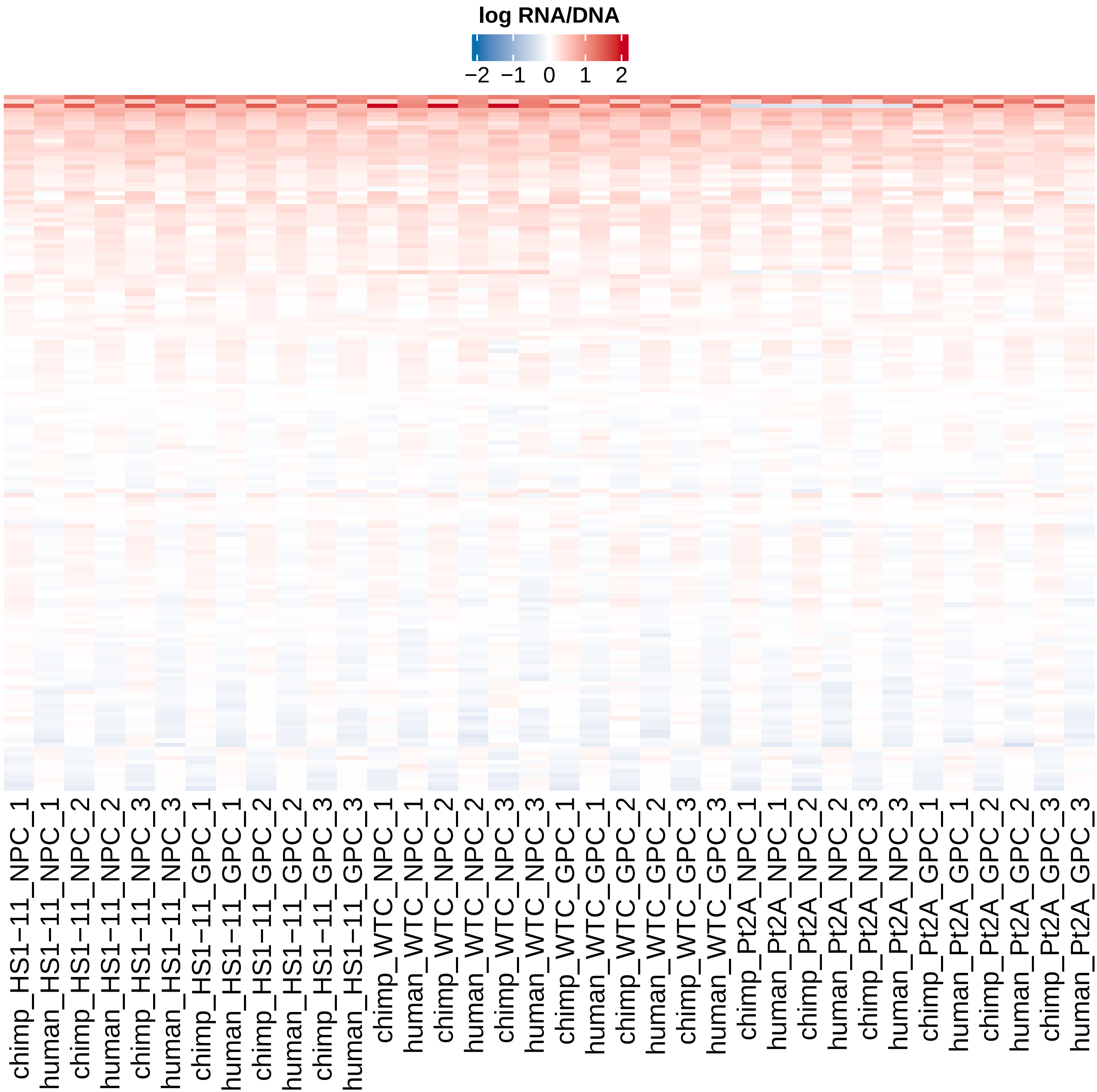

### Figure S11

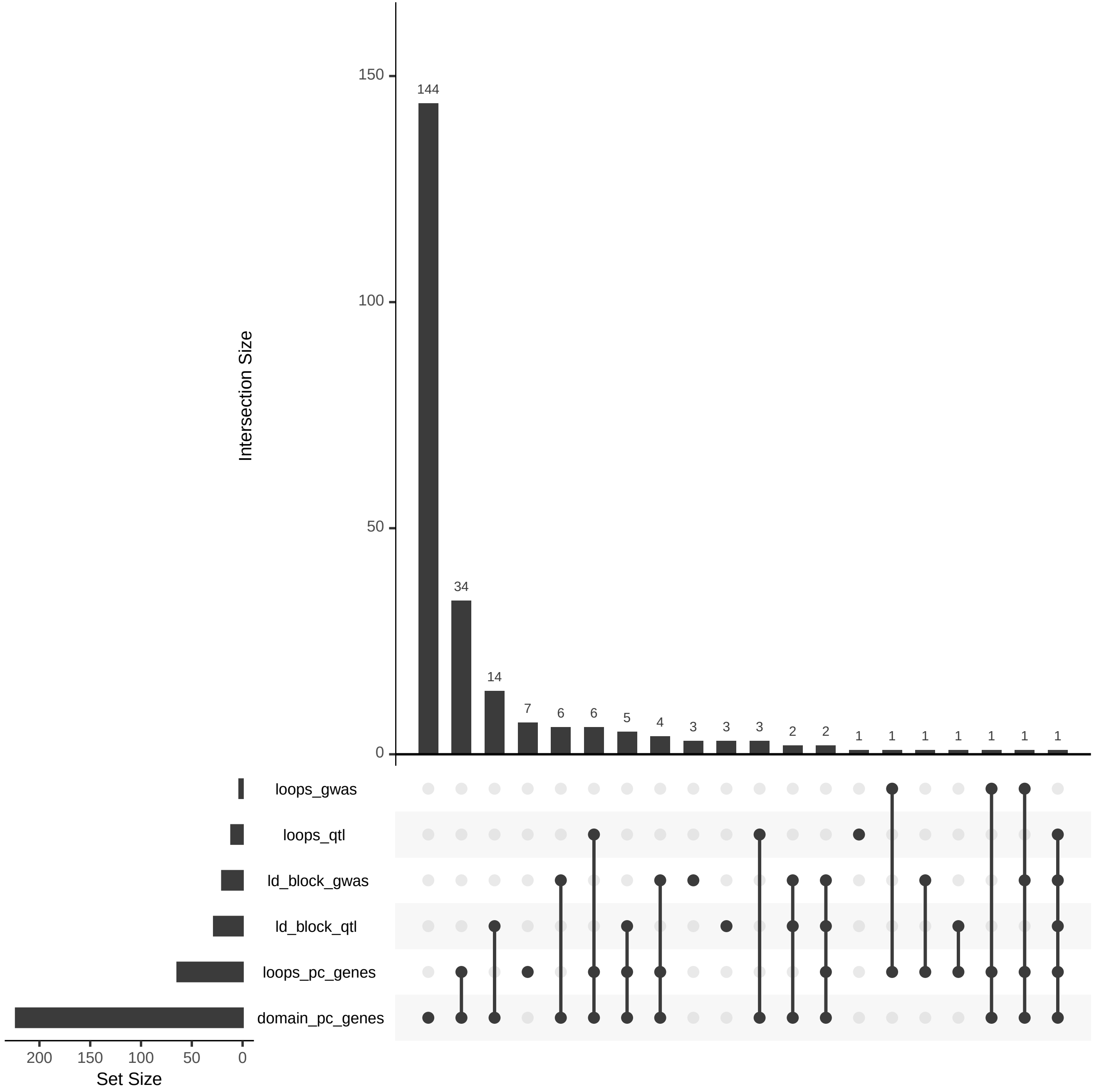

### Figure S12

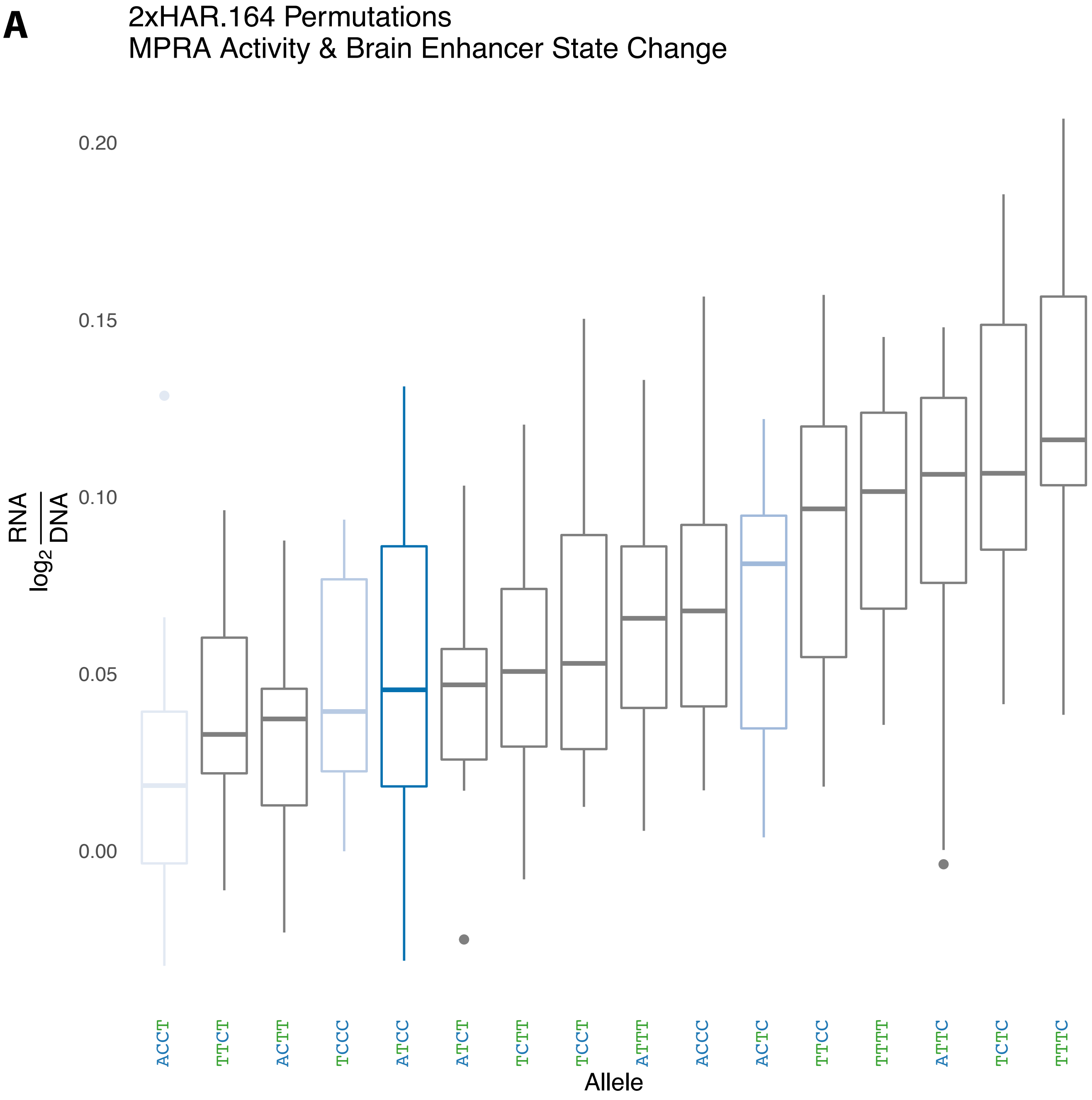

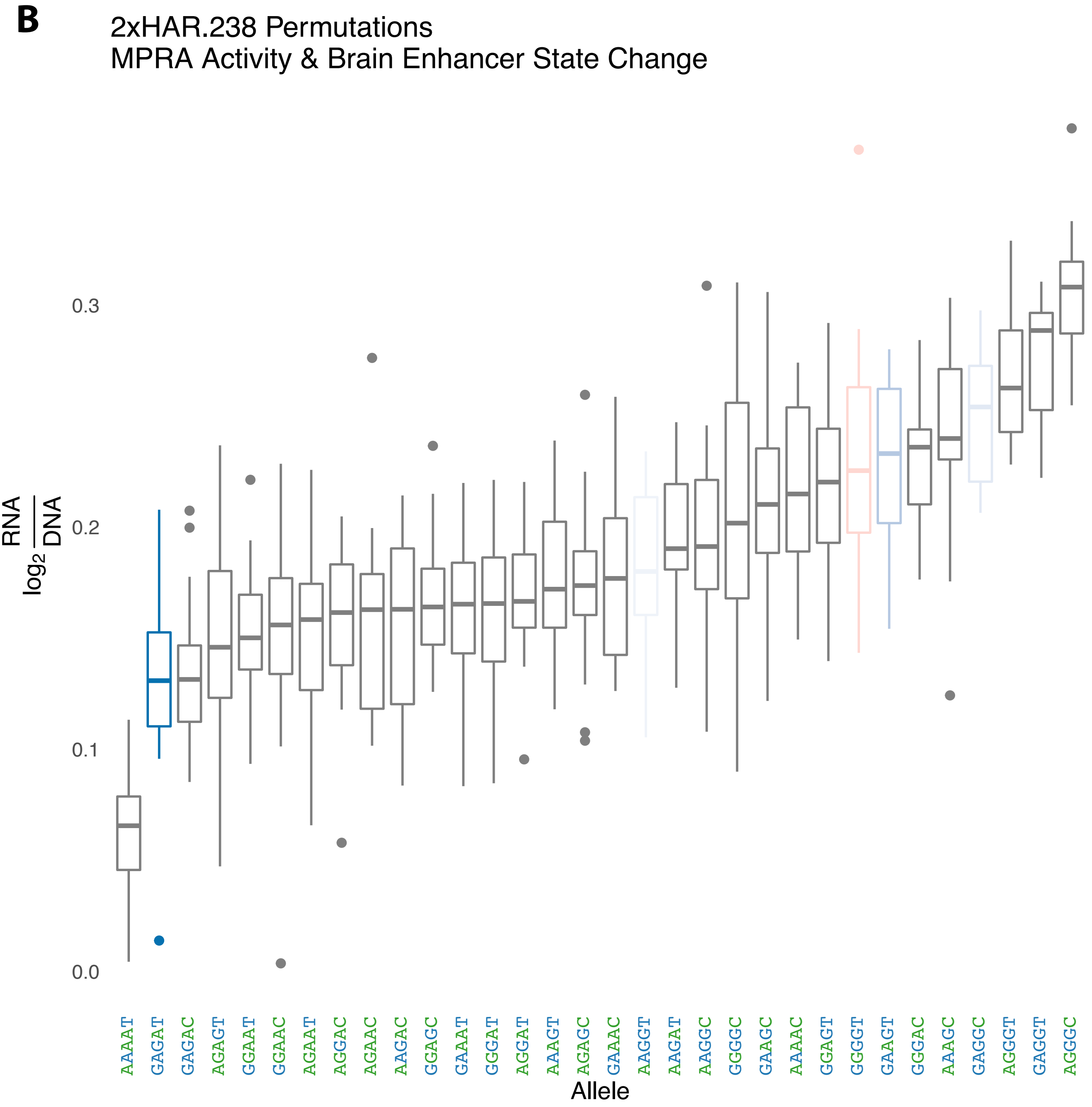

### Figure S13

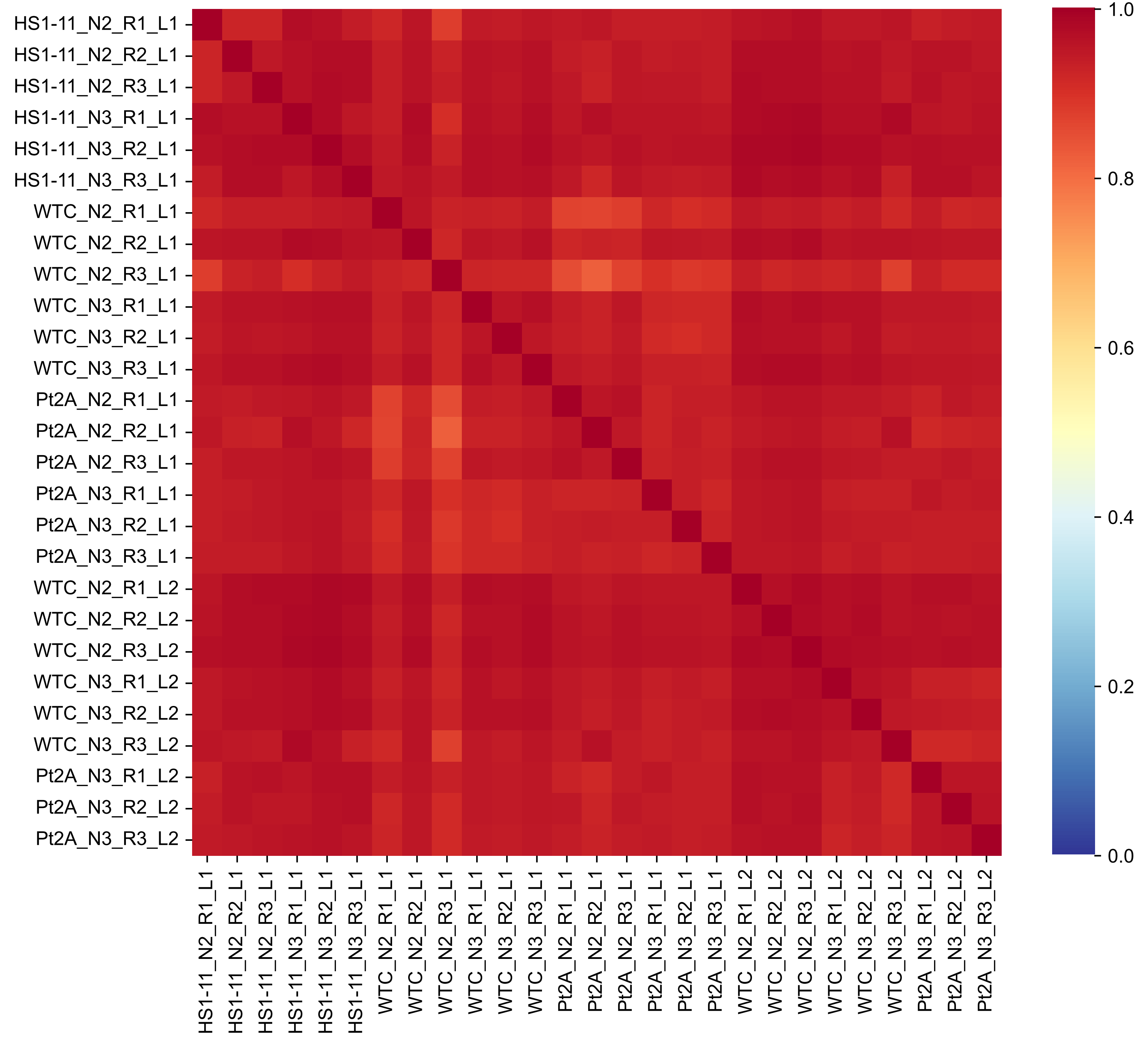

### Figure S14

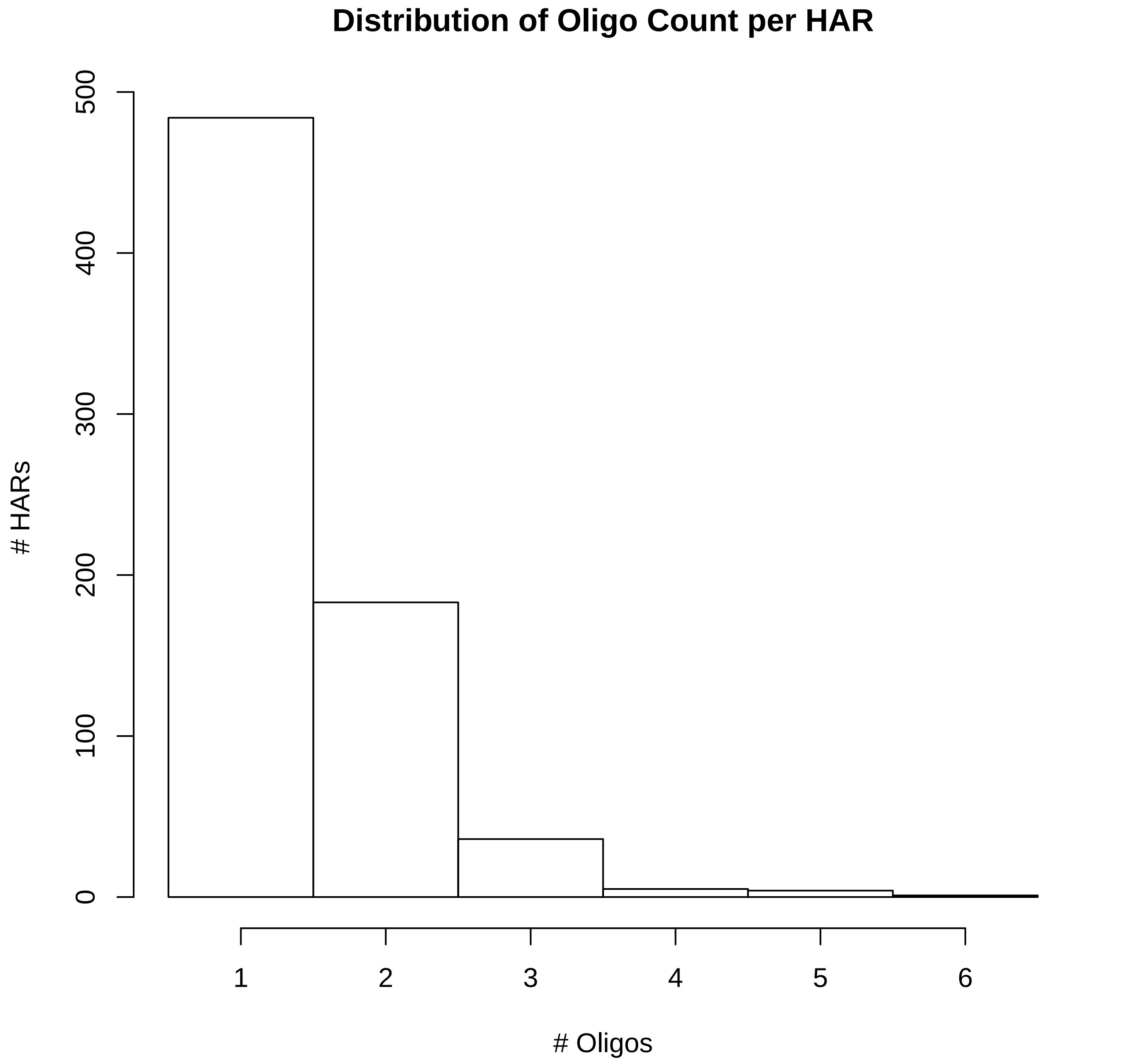
