## Supplementary material for "Machine-learning dissection of Human Accelerated Regions in primate neurodevelopment": Table S1

| Species | Stage | Donor | CapturePlateID | Number of Cells |
| --- | --- | --- | --- | --- |
| Human | N2 | WTC | 1771064068 | 49 |
| Chimp | N2 | Pt2 | 1771064069 | 27 |
| Chimp | N2 | Pt5 | 1784059104 | 20 |
| Human | N2 | Hs1 | 1784060057 | 50 |
| Chimp | N3 | Pt5 | 1784060058 | 84 |
| Chimp | N3 | Pt2 | 1784073115 | 69 |
| Human | N3 | WTC | 1784073116 | 57 |
