## Supplementary material for "Machine-learning dissection of Human Accelerated Regions in primate neurodevelopment": Table S4

**Table S4.** Telencephalon expression in HARs characterized in published mouse enhancer assays.

|  |  | **Any Tissue** | | **Brain** | | **Telencephalon** | | **NPC MPRA** |
| --- | --- | --- | --- | --- | --- | --- | --- | --- |
| **HAR** | **Refs** | **Human** | **Chimp** | **Human** | **Chimp** | **Human** | **Chimp** | **(this study)** |
| HAR2/2xHAR.3/  HACNS1 | 6,7 | Yes | Yes | No | No | No | No | Inactive |
| HAR5/2xHAR.557 | 7 | No | Not tested | No | Not tested | No | Not tested | Chimp biased |
| 2xHAR.5 | 2 | No | Not tested | No | Not tested | No | Not tested | Inactive |
| 2xHAR.20 | 1,2 | Yes | Yes | Yes | Yes | Yes | Yes | Human biased |
| HAR25 | 2 | Yes | Yes | Yes | Yes | No | No | Human biased |
| HAR31 | 5,7 | Yes | Not tested | No | Not tested | No | Not tested | Inactive |
| HAR34 | 2,7 | Yes | Yes | Yes | Yes | Yes | Yes | Inactive |
| 2xHAR.46 | 2 | Yes | Yes | No | No | No | No | Chimp biased |
| 2xHAR.59 | 7 | No | Not tested | No | Not tested | No | Not tested | Inactive |
| HAR69 | 2 | No | Not tested | No | Not tested | No | Not tested | Inactive |
| 2xHAR.82 | 7 | No | Not tested | No | Not tested | No | Not tested | Inactive |
| 2xHAR.90 | 2 | No | No | No | No | No | No | Active |
| 2xHAR.92 | 7 | No | Not tested | No | Not tested | No | Not tested | Active |
| 2xHAR.93 | 2 | Yes | Yes | No | No | No | No | Inactive |
| 2xHAR.97 | 7 | Yes | Not tested | No | Not tested | No | Not tested | Active |
| 2xHAR.99 | 2 | Yes | Yes | No | No | No | No | Inactive |
| HAR104 | 7 | Yes | Not tested | Yes | Not tested | Yes | Not tested | Inactive |
| 2xHAR.114 | 2 | Yes | Yes | Yes | Yes | No | No | Human biased |
| 2xHAR.116 | 2 | Yes | Yes | Yes | Yes | Yes | Yes | Inactive |
| HAR118 | 7 | Yes | Not tested | Yes | Not tested | Yes | Not tested | Chimp biased |
| HAR119/2xHAR.18 | 7 | No | Not tested | No | Not tested | No | Not tested | Chimp biased |
| HAR122 | 7 | Yes | Not tested | No | Not tested | No | Not tested | Active |
| 2xHAR.122 | 2 | No | No | No | No | No | No | Inactive |
| 2xHAR.128 | 2 | Yes | Yes | No | No | No | No | Inactive |
| 2xHAR.138 | 2 | Yes | Yes | Yes | Yes | Yes | Yes | Chimp biased |
| 2xHAR.142 | 3 | Yes | Yes | Yes | Yes | Yes | Yes | Inactive |
| HAR143 | 7 | Yes | Not tested | Yes | Not tested | Yes | Not tested | Inactive |
| HAR157 | 7 | No | Not tested | No | Not tested | No | Not tested | Inactive |
| HAR164 | 7 | Yes | Not tested | No | Not tested | No | Not tested | Inactive |
| 2xHAR.164 | 2 | Yes | Yes | Yes | Yes | Yes | Yes | Active |
| 2xHAR.170 | 2 | Yes | Yes | Yes | Yes | Yes | Yes | Human biased |
| HAR180/  2xHAR.427 | 7 | No | Not tested | No | Not tested | No | Not tested | Inactive |
| HAR196 | 7 | Yes | Not tested | Yes | Not tested | Yes | Not tested | Chimp biased |
| 2xHAR.222 | 2 | Yes | Yes | Yes | Yes | No | No | Active |
| 2xHAR.238 | 2,4,7 | Yes | Yes | Yes | Yes | Yes | Yes | Human biased |
| 2xHAR.240 | 2 | Yes | Yes | No | No | No | No | Inactive |
| 2xHAR.243 | 2 | No | No | No | No | No | No | Human biased |
| 2xHAR.247 | 7 | No | Not tested | No | Not tested | No | Not tested | Inactive |
| 2xHAR.274 | 2 | Yes | Yes | Yes | Yes | No | No | Human biased |
| 2xHAR.283 | 2 | No | No | No | No | No | No | Inactive |
| 2xHAR.287 | 2 | No | No | No | No | No | No | Inactive |
| 2xHAR.332 | 2 | No | No | No | No | No | No | Chimp biased |
| 2xHAR.349 | 2 | Yes | Yes | No | No | No | No | Active |
| 2xHAR.374 | 2 | Yes | Yes | No | No | No | No | Inactive |
| 2xHAR.377 | 2 | No | No | No | No | No | No | Inactive |
| 2xHAR.384 | 2 | No | No | No | No | No | No | Active |
| 2xHAR.408 | 2 | Yes | Yes | Yes | Yes | Yes | Yes | Human biased |
| 2xHAR.427 | 7 | No | Not tested | No | Not tested | No | Not tested | Inactive |
| 2xHAR.447 | 7 | Yes | Not tested | Yes | Not tested | Yes | Not tested | Inactive |
| 2xHAR.482 | 2 | No | No | No | No | No | No | Inactive |
| 2xHAR.499 | 7 | No | Not tested | No | Not tested | No | Not tested | Active |
| 2xHAR.514 | 7 | Yes | Not tested | Yes | Not tested | No | Not tested | Active |
| **TOTAL from tested** |  | 31/52 | 20/30 | 19/52 | 13/30 | 14/52 | 9/30 |  |
| **Active Percentage** |  | 59.61% | 66.67% | 36.54% | 43.33% | 26.92% | 30.00% |  |
| **Telecephalon:**  **Brain Percentage** |  |  |  |  |  | 73.68% | 69.23% |  |

**References**

[1] Aldea et al 2021: <https://doi.org/10.1073/pnas.2021722118>

[2] Capra et al 2013: <https://royalsocietypublishing.org/doi/10.1098/rstb.2013.0025>

[3] Kamm et al 2013b: <https://royalsocietypublishing.org/doi/10.1098/rstb.2013.0019>

[4] Norman et al 2021: <https://www.biorxiv.org/content/10.1101/2021.01.27.428524v1>

[5] Oskenberg et al 2013: <https://doi.org/10.1371/journal.pgen.1003221>

[6] Prabhakar et al 2008: <https://pubmed.ncbi.nlm.nih.gov/18772437/>

[7] VISTA: [http://enhancer.lbl.gov](http://enhancer.lbl.gov/)
